# Responses of the hyper-diverse community of canopy-dwelling Hymenoptera to oak decline

**DOI:** 10.1101/2023.07.25.550070

**Authors:** E. Le Souchu, J. Cours, T. Cochenille, C. Bouget, S. Bankhead-Dronnet, Y. Braet, P. Burguet, C. Gabard, C. Galkowski, B. Gereys, F. Herbrecht, B. Joncour, E. Marhic, D. Michez, P. Neerup Buhl, T. Noblecourt, D. G. Notton, W. Penigot, J.-Y. Rasplus, T. Robert, A. Staverlokk, C. Vincent-Barbaroux, A. Sallé

## Abstract

1. Forest decline and dieback are growing phenomena worldwide, resulting in severe, large-scale degradation of the canopy. This can profoundly alter the provision of trophic resources and microhabitats for canopy-dwelling arthropods.
2. In 2019, we assessed the effect of oak decline on the community of canopy-dwelling Hymenoptera. We selected 21 oak stands, and 42 plots, located in three forests in France, presenting contrasting levels of decline. Insects were sampled at the canopy level with green multi-funnel and flight-interception traps.
3. We collected a particularly diverse community of 19,289 insect individuals belonging to 918 taxa, ten larval trophic guilds and five nesting guilds.
4. Oak decline had no effect on the abundance or richness of the overall community, but significantly reshaped the community assemblages. Decline had contrasting effects depending on the taxa and guilds considered. Specialist parasitoids were more abundant at intermediate levels of decline severity while generalists were negatively affected. Taxa depending on ground-related resources and microhabitats were promoted. Saproxylic taxa were more abundant while xylophagous insects were negatively impacted.
5. Reduced leaf area index promoted several guilds, and the diversity of the overall community. While an increasing tree mortality rate enhanced the abundance and diversity of deadwood resources, it had negative impacts on several Hymenoptera guilds. Our results suggest that micro-environmental changes at the ground-level due to canopy decline have major cascading effects on the communities of canopy-dwelling Hymenoptera.
6. Our study highlights the relevance of studying Hymenoptera communities to investigate the outcomes of disturbances on forest biodiversity.

## INTRODUCTION

Global change is currently increasing the frequency, severity and spatial extent of major forest disturbances in Europe such as droughts, windstorms and wildfires (Seidl et al., 2017; Samaniego et al., 2018; Spinoni et al., 2018). Disturbances can markedly affect the amount, diversity and distribution of key trophic resources and microhabitats for forest arthropods (Cours et al., 2023). One of the main drivers of these changes is the degradation of the forest canopy (Sallé et al., 2021; Cours et al., 2023). A reduction in canopy cover can result from direct disturbance impacts on the trees themselves, but also from subsequent forest dieback or decline, as a progressive loss of tree vigour generally translates into crown dieback (Sallé et al., 2021). Decline and dieback can consequently affect the quality and quantity of foliage, and reduce the production of flowers, fruits and seeds in the crown (Ishii et al., 2004; Günthardt-Goerg et al., 2013; Hu et al., 2013). On the other hand, they may also increase the amount and diversity of deadwood resources and tree-related microhabitats such as fruiting bodies of opportunistic fungi, trunk cavities or perched deadwood (Sallé et al., 2021; Larrieu et al., 2022; Bouget et al., 2023; Zemlerová et al., 2023). Increased canopy openness also considerably affects microclimates and trophic resources for arthropods in the lower strata of forest ecosystems for instance, by promoting the accumulation and diversification of floral resources in the herbaceous layer, or of deadwood resources on the ground (Cours et al., 2023). Consequently, forest dieback and decline can promote forest biodiversity to a certain extent by increasing structural complexity at multiple scales, as predicted by the pulse dynamics theory (Cours et al., 2022). This theory postulates that pulse events like disturbances affect resource ratios, storage and availability, energy fluxes, spatiotemporal patch dynamics and biotic trait diversity, and predicts that species and trait diversity at the landscape scale should increase with patch distribution and resource heterogeneity (Jentsch and White, 2019). These modifications can profoundly reshape communities of forest arthropods (Viljur et al., 2022; Cours et al., 2023). The impact of canopy dieback on forest arthropods has been investigated for several taxonomic groups and functional guilds, including some specifically dwelling in the canopy (Martel & Mauffette, 1997; Stone et al., 2010; Sallé et al., 2020; Vincent et al., 2020). While the responses of canopy arthropods are largely mediated by their functional and/or trophic guild (*e.g.*, Sallé et al., 2020), idiosyncratic responses still occur among the taxa of a same guild or taxonomic group (*e.g.*, Vincent et al., 2020). Consequently, the impacts of disturbance-driven changes in canopy structure on arthropod communities are still difficult to predict, especially in the largely under-studied temperate forest canopy (Sallé et al., 2021; Cours et al., 2023). Considering that canopy mortality rate has been on the rise for several decades in Europe (Senf et al., 2018), further studies on how decline and dieback affect canopy-dwelling arthropods are urgently needed, especially for the taxonomic groups that have received limited attention to date, like the Hymenoptera.

Hymenoptera is a hyper-diverse group of insects, sometimes regarded as the most speciose animal order (Forbes et al., 2018). This group plays a wide variety of functional roles in temperate forests. Ants and predatory and parasitic wasps help to regulate forest pests (Hilszczański, 2018). Bees and wasps pollinate forest plants (Motten, 1986; Yumoto, 1987), while ants can disperse seeds (Rico-Gray and Oliveira, 2008). Hymenoptera nests, galleries and galls also provide microhabitats for many species (Stone et al., 2002). Wood-nesting bees and ants, as well as the larvae of xylophagous species, contribute to the decomposition process of wood (Ulyshen, 2016). Phyllophagous and xylophagous species can be major pests for trees (Lyytikäinen-Saarenmaa and Tomppo, 2002; Slippers et al., 2015). The Hymenoptera, especially parasitic wasps, can be used as bioindicators in forest ecosystems (Maleque et al., 2009). Their communities can be largely influenced by stand-related variables such as stand composition (Fraser et al., 2007; Ulyshen et al., 2011a), deadwood resources (Hilszczański et al., 2005; Ulyshen et al., 2011a; Jonsell et al., 2023) and silvicultural practices (Lewis and Whitfield, 1999; Hilszczański et al., 2005; Rappa et al., 2023). Pollinators in particular are sensitive to canopy opening and the subsequent positive impacts on the floral resources in the herbaceous layer and on ground-nesting sites (Wermelinger et al., 2017; Burkle et al., 2019; Viljur et al., 2022; Cours et al., 2023).

Many Hymenoptera use the canopy as a hunting ground, a nesting site or a breeding site (Ulyshen, 2011). The canopy can also act as a dispersal corridor for species with a preference for open habitats (Pucci, 2008; Di Giovanni et al., 2017). As for other forest insects, the Hymenoptera community exhibits a conspicuous vertical stratification in temperate forests (Smith et al., 2012). Certain ecological guilds like the predatory and parasitic wasps, and the pollinators, differ significantly between the ground level and the canopy (Sobek et al., 2009; Ulyshen et al., 2011b; Urban-Mead et al., 2021), and several taxa are specific to the canopy layer (Ulyshen et al., 2010; Di Giovanni et al., 2017). While some studies have investigated the vertical stratification of certain taxonomic groups or ecological guilds of bees and wasps in temperate forests, few have taken into account the whole Hymenoptera community (*e.g.*, Vance et al., 2007), and none have considered this community’s response to forest decline.

In the present study, we investigated the impact of oak decline on the community of canopy-dwelling Hymenoptera. We chose to focus on oak forests because they are the dominant forest ecosystem in western Europe; they shelter a diverse entomofauna (Kennedy and Southwood, 1984); finally, they have suffered from historic decline in France (Nageleisen, 2008), and have recently undergone severe summer droughts (in 2018 and 2019) (Saintonge and Goudet, 2020). Our first objective was to characterise the little-known community of canopy-dwelling Hymenoptera in temperate oak forests. Our second objective was to assess how the severity of forest decline affected the richness and taxonomic composition of this community. Our third objective was to evaluate the effects of decline on certain ecological guilds of Hymenoptera, and how changes in stand-related variables drive those effects. We focused on the trophic guilds of larvae and nesting guilds. For these guilds, we formulated three hypotheses. First (H1), we expected to find positive effects of increasing decline severity on pollinivorous / nectarivorous and phyllophagous species associated with host plants other than oak due to the increased diversity and amount of plant-resources in the herbaceous and shrub layers, commonly observed when canopy closure decreases (Romey et al., 2007; Lu et al., 2019; Cacciatori et al., 2022). Likewise, we expected that soil-nesters would be promoted by more open conditions resulting in warmer soil temperatures. We also hypothesised that xylophagous, saproxylic and wood-nester taxa would benefit from oak dieback and the subsequent increased amount and diversity of deadwood. Conversely (H2), we expected to find negative effects on the guilds that feed on oak foliage, for example, gall-inducing and phyllophagous taxa, due to the decrease in the quality and quantity of leaves. Finally (H3), we hypothesised that decline severity would have contrasting effects on parasitoids and polyphagous taxa, depending on their hosts and trophic resources. In addition, we expected that specialist parasitoid taxa, highly sensitive to habitat modification (Hilszczański, 2018), would exhibit pronounced changes along the gradient of decline severity. On the other hand, we anticipated that generalist taxa would be favoured by the disturbed environmental conditions in severely declining stands (Devictor et al., 2008).

## MATERIAL AND METHODS

### Study sites

Three oak-dominated (*Quercus petraea* (Matt.) Liebl. and *Quercus robur* L.) French forests were sampled in 2019 (Fig. 1, Tab. S1): one near Orléans (Forêt domaniale d’Orléans, 47°98’97”81N, 1°95’44”E), one near Vierzon (Forêt domaniale de Vierzon, 47°26’89”N, 02°10’74”E) and one near Marcenat (Forêt domaniale de l’Abbaye, 46°21’12”N, 3°36’13”E). All the forests were managed by the French National Forest Service. The Orléans forest covers 35,000 ha and is dominated by oaks (47%) and pines (*Pinus sylvestris* L., 51%). The Vierzon forest covers 7,500 ha and is also dominated by oaks (61%) and pines (*P. sylvestris* and *P. pinaster* Ait., 31%). The Marcenat forest covers 2,100 ha and is dominated by oaks (71%), pines (11%) and Douglas fir (8%). The Vierzon forest suffered the most from decline. Several decline events had been recorded there since 1920, and massive oak mortality had occurred in several stands (Douzon, 2006). In the Marcenat forest, two summer droughts (2018-2019) had led to severe decline in several stands (Saintonge and Goudet, 2020). Conversely, no major decline event had been reported in the Orléans forest.

**Figure 1.**
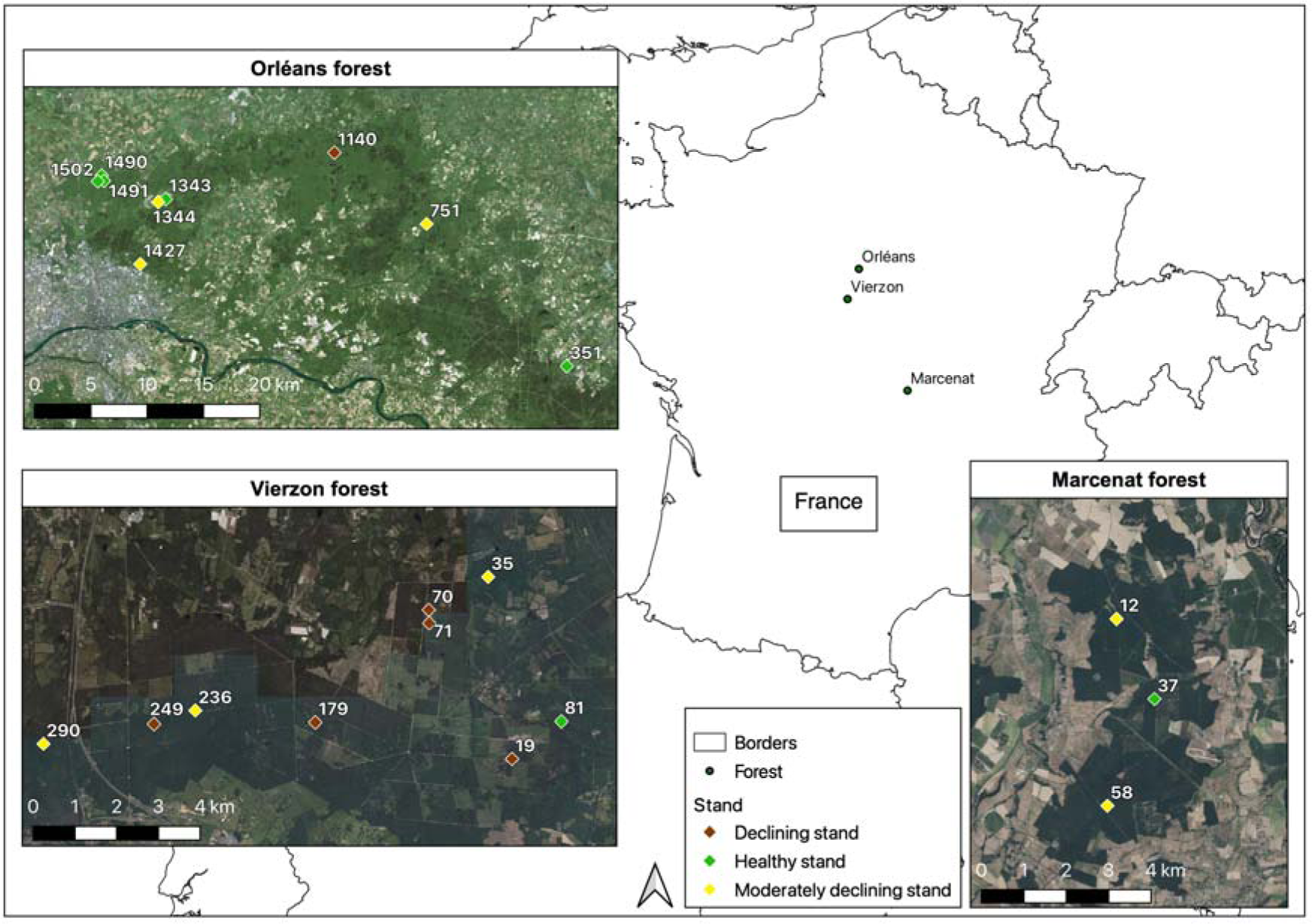
Map of France showing the location of the three oak forests studied, and the 21 stands sampled. Colors indicate the level of stand decline, according to the proportion of declining trees: < 30%: healthy stand; 30-60%: moderately declining stand; > 60%: severely declining. Details on the stands can be found in Tab. S1.

In each forest, we first selected oak-dominated stands exhibiting contrasting levels of decline (nine in Orléans, nine in Vierzon and three in Marcenat), on the basis of local forester knowledge and visual examination. Even in forests with no major decline, declining trees could be found, in stands with temporarily waterlogged soils for instance. These 21 stands were separated by at least 500 m, with a minimum surface area of 3 ha (Fig. 1). In each stand we selected two living trees to hang our traps, at least 50 m apart, one healthy and one declining if both tree types were present. These two trees were at the center of two plots where we secondly quantified the decline severity (see below). In each stand, we recorded average tree height, density and diameter, and the stand tree species composition (Tab. S1). We also evaluated the level of decline according to the DEPERIS protocol (Goudet and Nageleisen, 2019). This protocol makes it possible to quantify decline at the tree level by evaluating the percentage of dead branches and ramification loss in the crown. Based on these criteria, each tree was assigned to a decline class ranging from A (no decline) to F (severe decline). Trees in the A–C classes were considered healthy. Trees in the D–F classes were considered in decline. We used this protocol to describe tree decline at two embedded spatial scales: (i) the ten closest oaks surrounding each trap-bearing tree, hereafter referred as a “*plot*”; and (ii) 30 oaks in the stand where the two plots were located (*i.e.,* the 20 trees in the two monitored plots and ten trees in an additional plot located between the two monitored plots), hereafter referred as a “*stand*”. Decline level was evaluated in January or February in 2019 and 2020, *i.e.,* before and after Hymenoptera sampling, and we averaged the 2019 and 2020 values for our analyses. We classified our plots and stands into three decline categories depending on the proportion of declining oak trees. When the proportion of declining trees was < 30%, we considered the plot/stand “*healthy*”; when the proportion was between 30% and 60%, the plot/stand was considered “*moderately declining*”; and when the proportion was > 60%, the plot/stand was considered “*severely declining*”.

In the early summer of 2019, we used a plant canopy analyzer LAI-2200C with a half view cap attached to the optical sensor to quantify the leaf area index (LAI) in the 12 stands in the Orléans and Vierzon forests. More than 50 measurements were carried out at each plot at twilight below the canopy along two diagonal transects; these values were then compared with above-canopy readings taken outside the plot. We applied an ellipsoidal post-processing correction to the first three zenith angles and used the FV2200 software to compute LAI by the gap fraction method. Then, in October 2020, we used a Bitterlich relascope with an opening angle corresponding to counting factor n° 1 (ratio 1/50) with a mean plot area of about 0.3 ha (Bouget et al., 2023) to characterise the woody resources in each of the 18 plots in the Orléans and Vierzon forests. For each tree, we recorded its status (*i.e.,* living tree, standing dead tree (snag) or downed dead tree (log)), decay stage, tree species and diameter at breast height (DBH). We defined the mortality rate as the basal area of standing and lying deadwood over the total basal area of all the recorded trees (*i.e.,* the sum of both living and dead trees). We calculated the overall volume of living trees and deadwood. For deadwood, we calculated the volume for each status (standing or downed deadwood), decay stage (slightly vs. highly decayed) and diameter class (deadwood DBH < 40 cm vs. ≥ 40 cm). We compiled all this information (*i.e.,* status, species, decay stage and DBH) to calculate deadwood diversity. Finally, we used the dbFD function from the FD R-package to calculate the mean status value of the woody resources and the dispersion value for each plot, based on an ordinal scale (*i.e.,* living trees = 3, snags = 2, logs = 1). Further information on the acquisition and processing of forest structure data is presented in Bouget et al. (2023).

### Insect sampling

We hung two types of traps in each selected tree: one green multi-funnel trap (ChemTica Internacional, San José, Costa Rica) with 12 fluon-coated funnels, and one cross-vane flight interception trap (Polytrap^TM^). The first type is used to sample borers like *Agrilus* (Santoiemma et al., in press); the experiment was initially designed to sample the community of borers along the decline gradient in oak stands. This type of trap is also efficient for sampling guilds of foliage-associated insects (Sallé et al., 2020), and was sometimes used to sample Hymenoptera (Barnes et al., 2014; Haavik et al., 2014; Skvarla et al., 2016). Green multi-funnel traps in particular have proven to be quite effective in collecting sawflies (Skvarla et al., 2016). Flight interception traps are commonly used in forests to sample saproxylic beetles (Bouget et al., 2008).

Both types of traps were suspended among the lower branches in the canopy (*i.e.,* approximately 10 - 15m above the ground). The collectors were filled with a 50% (v/v) monopropylene glycol - water solution with a drop of detergent. The traps were in place from the end of March to the beginning of September 2019, during the insects’activity period, and they were emptied every month.

### Identification and ecological trait list

All sampled Hymenoptera were sorted and identified to the family or superfamily level, according to the identification key by Goulet and Huber (1993). Then the specimens were sent to taxonomical experts who identified them to the genus, species or morphospecies level. Only the Cynipidae and Figitidae were not identified at a lower taxonomic level.

We arranged the taxa into ten larval trophic guilds: gall-inducers, parasitoids, oak-associated phyllophagous larvae, other phyllophagous larvae, phytophagous larvae (*i.e.,* feeding on plant parts other than leaves, for example, seeds), carnivorous larvae, social polyphagous larvae, xylophagous larvae and pollinivorous/nectarivorous larvae. Parasitoids were separated into idiobionts (*i.e.,* larvae develop on non-growing host stages or paralyzed hosts) and koinobionts (*i.e.,* the host continues feeding and growing during parasitism). This is often used as a surrogate of whether a parasitoid is a generalist (idiobiont) or a specialist (koinobiont) (Quicke, 2015). Preferred larval habitat (larval nesting guild) was also considered, when known. Taxa were divided into gall-nesters (*i.e.,* gall-inducers and their inquiline parasitoids), soil-nesters, stem-nesters (generally in hollow plant stems) and wood-nesters. For the latter three guilds, we separated the specialists (spe.) from the generalists (gen.), nesting in wood and/or soil and/or stems. Specialist wood-nesters were considered analogous to saproxylic taxa (Jonsell et al., 2023).

### Data analysis

All analyses and graphs were performed in R, version 4.1.1 (R Core Team, 2021). We used the ggplot2 R-package (Wickham, 2016) to produce the figures. Unless otherwise stated, analyses were realised with the Hymenoptera community collected at the plot level, and the proportion of declining trees at both plot and stand scales. The figures with stand decline levels are included in the body of the present article since they are more informative than the figures with plot decline levels, which are available in the Supplementary Data. We also provide the figures with ecological guilds in the main text, the figures with taxonomic families or species are provided in the supplementary data.

Some specimens of Cynipidae, Figitidae, Halictidae, Andrenidae, Pteromalidae, Encyrtidae and Platygastridae were not identified, either because of a lack of expertise or because the specimens were too damaged for identification. These unidentified specimens were removed from the community analyses.

To extrapolate the species richness for the whole community (*i.e.,* γ-diversity) and for each decline category, we first plotted species rarefaction curves (functions *iNEXT* and *ggiNEXT*, iNEXT R-package; Hsieh et al., 2016) and calculated Chao, Jackknife, Jackknife 2 and Bootstrap diversity indices (function *specpool*, vegan R-package; Oksanen et al., 2022). We used the additive diversity partitioning method (Lande, 1996; Crist et al., 2003) to evaluate the contribution of α-and β-diversity to the γ-diversity of canopy-dwelling Hymenoptera over the entire sampled area (function *adipart*, index = “richness”, 1,000 permutations, vegan R-package). We tested whether observed species richness differed from the one we expected by chance (null hypothesis). At the plot scale, we used three levels of differentiation: plot, plot decline category and the whole sampled area. α plot corresponds to the average species diversity per plot, β plot to the diversity among plots and β cat_plot to the diversity among plot decline categories. γ-diversity is the sum of α plot, β plot and β cat_plot (γ = α plot + β plot + β cat_plot). Consequently, four levels of differentiation were included: plot, stand, stand decline category and the whole sampled area. At the stand scale, four levels of differentiation were included: plot, stand, stand decline category and the whole sampled area. As an additional measure of β-diversity, we calculated the dissimilarity between plots, stands, decline categories and pairs of decline categories to separate β-diversity (Sorensen dissimilarity index) into β-turnover (Simpson dissimilarity index) and β-nestedness (nestedness-resultant fraction of Sorensen dissimilarity) (function *beta.multi*, Sorensen family, betapart R-package; Baselga, 2010; Baselga and Orme, 2012). β-turnover indicates the replacement of some species by others, while β-nestedness indicates that species assemblages are subsets of species occurring at larger spatial scales.

We assessed the influence of plot and stand decline categories (*i.e.,* healthy, moderately declining and severely declining) on the community of Hymenoptera in each plot by performing non-metric multidimensional scaling analyses (NMDS, function *metaMDS*, k = 3, Bray-Curtis index, 1,000 permutations, vegan R-package), and pairwise PERMANOVA (function *adonis2*, Bray-Curtis index, 999 permutations, vegan R-package). Species with less than ten individuals were excluded from these analyses.

We assessed the effect of oak decline on the abundance and species richness of the overall community, of each taxonomic family (when n > 30 ind.), and on the abundance of each taxa (when n > 30 ind.). We also assessed the oak-decline effect on each guild (i.e., larval trophic guilds and nesting guilds). To do this, we used generalised linear mixed-effects models (GLMMs; function glmer, glmer.nb or lmer, lme4 R-package; Bates et al., 2022) fitted for the Poisson family, the negative binomial family or the log-normal distribution (*i.e.*, log(x+1) transformed). We first selected the best suited family distribution, with the fitdistrplus R-package (function *fitdist* and *gofstat*; Delignette-Muller and Dutang, 2015). We then performed GLMMs with either abundance or species richness at both scales (plot and stand). The decline variable was used as either a linear term or a quadratic term to be better fit our dataset. We added forest and stand as nested random effects on the intercept in the mixed models to account for repeated measurements and the spatial configuration of the sampling design. Since some traps were not continuously functional, for example, because they fell from the tree, we also added an offset of the log-number of effective traps across the entrapment season. The offset was weighted by the efficacy of each type of trap (*i.e.,* multi-funnel vs. flight interception traps), estimated with the average percentage of specimens collected by each type of trap. To assess model quality, we used marginal R² (function *R2*, performance R-package; Lüdecke et al., 2021). We used the differences in AICc scores (function *AICc*, AICCmodavg R-package; Mazerolle, 2023) to compare the fit among models. When the linear and quadratic models were undifferentiated, we show only the results for the simplest one, *i.e.,* the linear model. We also used the indicspecies R-package (function *multipatt*, abundance data, func = “IndVal.g”, 1,000 permutations) to identify indicator species for healthy, moderately-declining and severely-declining stands and plots (Dufrene and Legendre, 1997; De Cáceres and Legendre, 2009; De Cáceres et al., 2010).

In a second step, we implemented Structural Equation Modelling (SEM, piecewiseSEM R-package; Lefcheck et al., 2022) to evaluate the cascading effects of tree decline (*i.e.,* the proportion of declining trees) and tree mortality (*i.e.,* the proportion of dead trees) on taxonomic families and ecological guilds through changes in canopy closure (LAI), and changes in variables related to living trees (volume, density, and DBH), and deadwood resources. SEM was used only for the plots located in the Orléans and Vierzon forests. LAI was originally measured in 12 plots in the Orléans and Vierzon forests, and correlated strongly with tree decline rate at the stand scale (Fig. S1). Consequently, in order to avoid excluding additional plots in the SEM analysis, we extrapolated LAI from the tree decline rate in the six remaining plots using the equation of the regression curve (Fig. S1). We studied Hymenoptera responses through (i) the abundance of each taxonomic family, and (ii) the abundance and species richness of each ecological guild (see above). Response variables that did not respect assumptions of residual normality and homoscedasticity for linear regressions were log-transformed. In addition, in a first set of mixed models, we added forest identity (*i.e.,* Orléans or Vierzon) as a random effect in the *lmer* function from lme4 R-package. We finally dropped this random effect since it was redundant with the fixed effects and caused model non-convergence. Finally, we used a linear model with the *lm* function. Several plots were installed close to wetlands. We added this information, as a binary variable (*i.e.,* wetland / no wetland), to the models as a fixed effect since the characteristic might affect soil-nesters and phyllophagous species associated with plants in the herbaceous and shrub layers. We also added an offset term of the log-number of effective traps across the entrapment season to our models. Finally, we accounted for the increased type I error risk due to multiple testing (16 parameters) with the Bonferroni correction (family-wise error rate = 0.05/16 = 0.0031).

## RESULTS

### Overview of the community of canopy-dwelling Hymenoptera

We collected 19,289 individuals, belonging to 54 families and 918 taxa (Fig. 2; Tab. S2). The most abundant families were the Ichneumonidae (22% of the specimens), the Cynipidae (22%) and the Perilampidae (14%), while the most diverse were the Ichneumonidae (321 taxa), the Braconidae (121) and the Tenthredinidae (88) (Fig. 2). Most of the community was composed of singletons (344 taxa) and doubletons (129), and only 198 taxa occurred with more than ten individuals (Tab. S2). Several species cited for the first time in France were sampled, especially members of the Ichneumonidae family (Tab. S2). We also collected 35 bee species registered on Worldwide or European Red Lists (Tab. S2). Among them, three species were near-threatened, 13 species were classified as “data deficient”, and 19 as “of least concern”.

**Figure 2.**
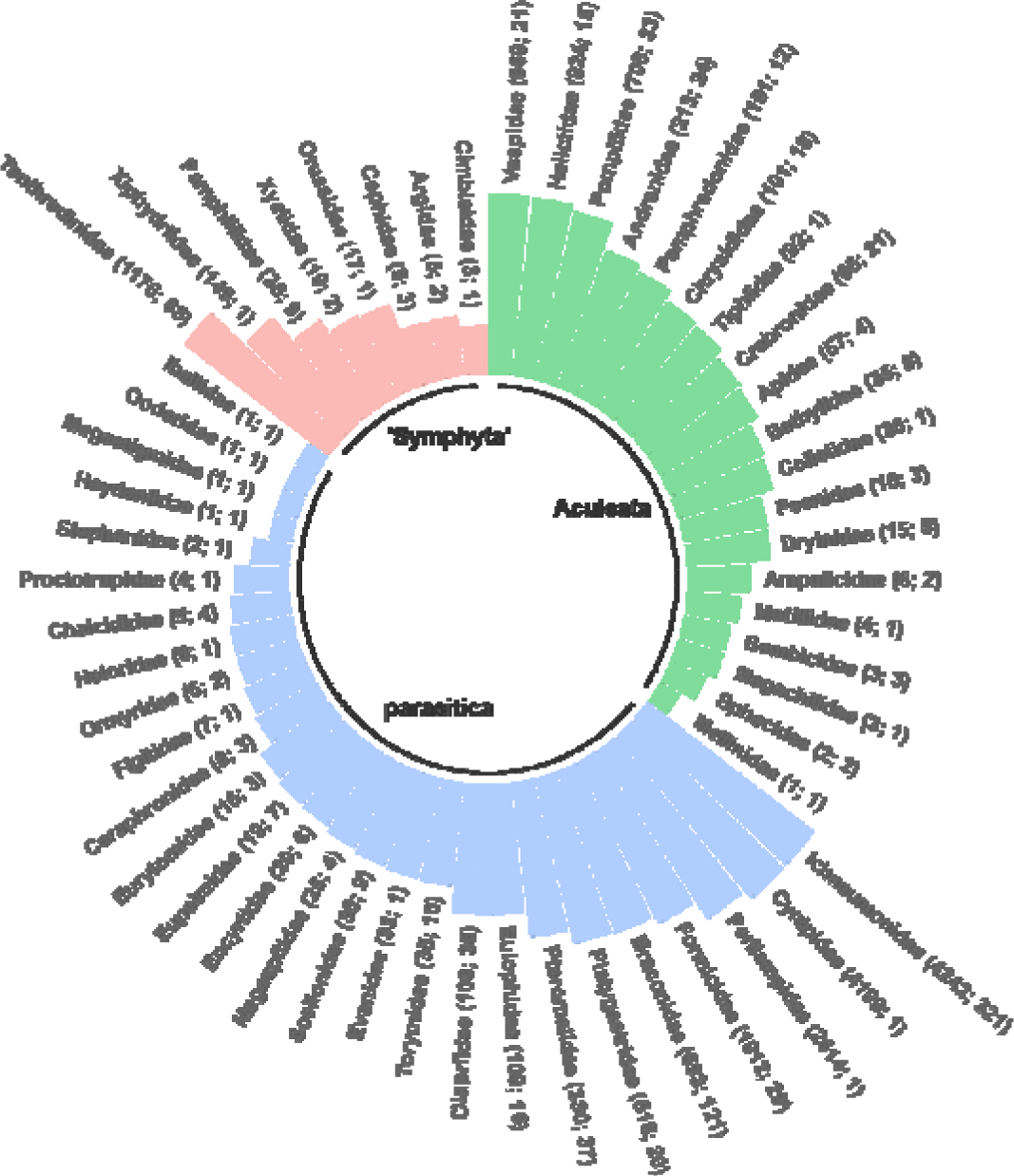
Overview of the diversity and abundance (log-transformed) of the Hymenoptera families sampled in oak canopies. Families are arranged in three taxonomic groups: “Symphyta”, Apocrita Aculeata and Apocrita parasitica. Numbers between brackets indicate the number of individuals and the number of taxa identified for each family.

The activity of most families peaked in the spring (Fig. S2), and the community was the most abundant (64% of total specimens) in April. Species richness was more evenly distributed throughout the sampling period, but also peaked in April (48% of all taxa). Overall, the green multi-funnel traps clearly outperformed the flight interception traps (93% vs. 7% of total abundance, 94% vs. 28% of all taxa) (Fig. S3); for each guild and family, both abundance and species richness were higher in the multi-funnel traps.

According to our species-diversity estimators (Tab. S3), there should have been between 1,074 and 1,505 species in the entire sampled area. We collected between 61% and 85% of these estimations, and consequently, the rarefaction curve for the whole community barely approached an asymptote (Fig. 3A). Both moderately declining and severely declining plots and stands were predicted to shelter slightly more taxa than healthy ones (Tab. S3, Figs. 3A & S4A). The communities in healthy and severely declining stands were clearly distinct, but both were partially similar to those in moderately declining stands (Fig. 3B). At the plot scale, a similar pattern was observed, although the divergence was less marked (Fig. S4B). Consequently, the differences among decline categories were significant at both the plot and stand scales, except between moderately declining and healthy plots (Tab. S4).

**Figure 3.**
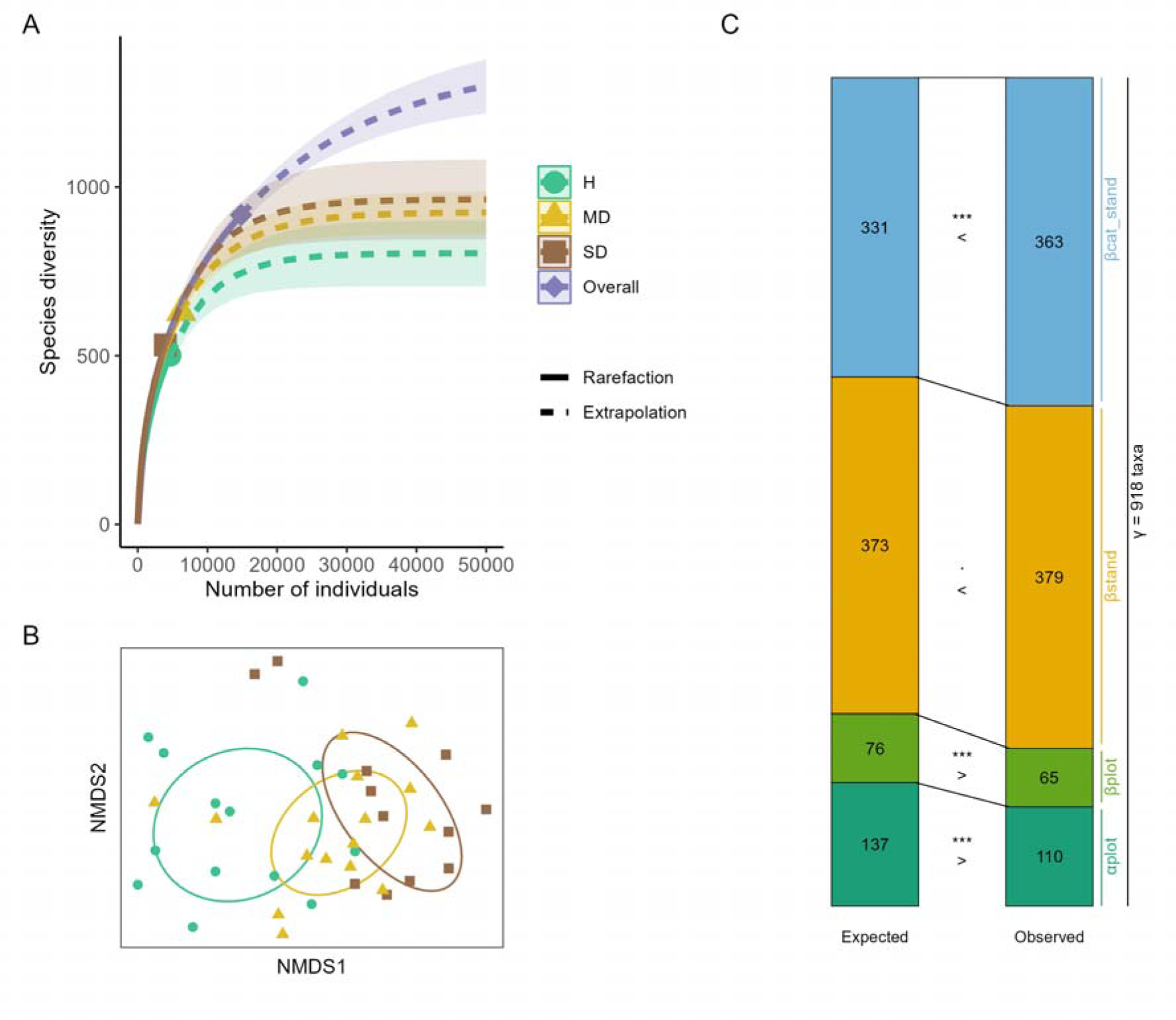
Responses of the community of canopy-dwelling Hymenoptera, collected from three oak forests, 21 stands and 42 plots, to stand decline severity. **A)** Rarefaction curves for the overall dataset and for healthy (H: < 30% of trees are declining), moderately declining (MD: 30-60% of trees are declining) and severely declining (SD: > 60% of trees are declining) stands. **B)** NMDS ordination (k=3, stress=0.18) of species composition per plot, grouped by stand decline category. **C)** Global additive partitioning of species richness of the canopy-dwelling Hymenoptera in the three oak forests, at plot and stand scales. Four levels are represented: α plot (within plot), β plot (among plots), β stand (among stands) and β cat_stand (among levels of stand decline). Significance levels correspond to the difference between expected and observed values, with ***: p < 0.001; . : p < 0.1. H = healthy stands, MD = moderately declining stands, SD = severely declining stands, Overall = all stands. See main text for details on stand decline categories.

### Contribution of α- and β-diversity to γ- diversity

The additive partitioning of species richness showed that, at the plot scale, *β plot* did not present any significant difference between observed and expected richness (Fig. S4C). At the stand scale, the observed richness of *β plot* was significantly lower than expected (Fig. 3C), while observed *β stand* was similar to predicted *β stand*. Conversely, decline categories contributed largely to γ-diversity. Observed *β cat_plot* and *β cat_stand* were significantly higher than expected and respectively represented 39% and 40% of the γ-diversity.

Overall *β*-diversity was mainly due to species turnover, among all plots and stands (Tab. 1), and between each pair of plots (Tab. S5) and stand decline categories (Tab. 1). At the plot scale, *β* nestedness was very low. At the stand scale, however, *β* nestedness was a significant part of the *β*-diversity between moderately declining stands and healthy ones (22%), as well as between moderately and severely declining stands (16%).

**Table 1.**
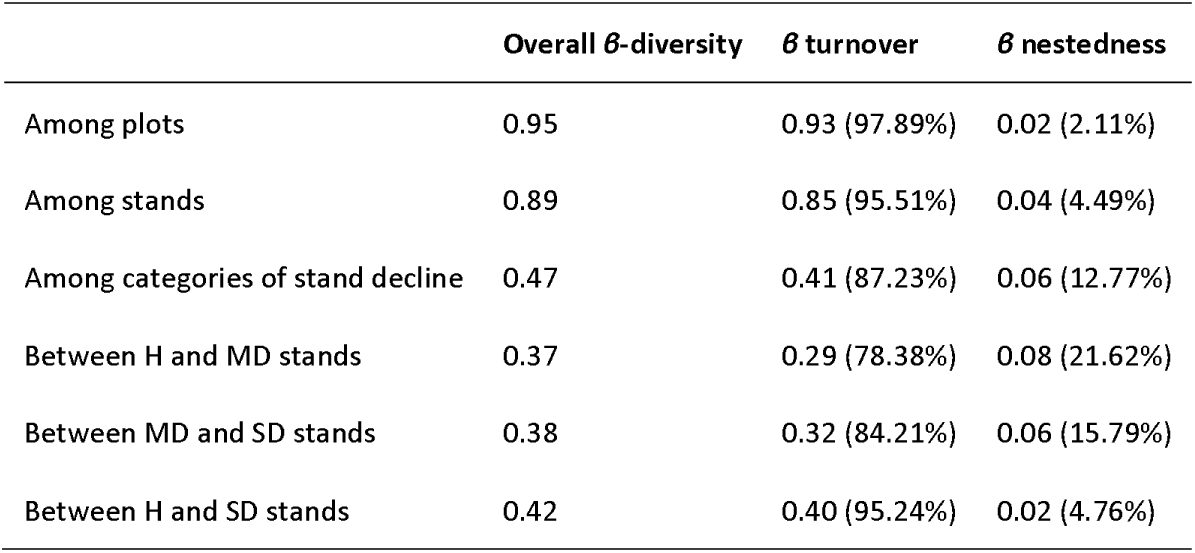
Overall β-diversity, β turnover and β nestedness between pairs of stand-decline category. Overall β-diversity corresponds to Sorensen dissimilarity, β turnover to Simpson dissimilarity and β nestedness to the difference between Sorensen and Simpson dissimilarity. H = healthy stands, MD = moderately declining stands, SD = severely declining stands.

### Indicator species

At the plot scale, one species was associated with healthy plots, 11 with moderately declining plots and 11 with severely declining plots (Tab. S6). At the stand scale, five species were associated with healthy stands, five with moderately declining stands and seventeen with severely declining stands (Tab. S7). Four species were indicators at both plot and stand scales. The majority of the indicators were parasitoids. Several sawflies that feed on non-oak host plants were also indicators of severely declining stands, and a few species with pollinivorous/nectarivorous larvae were indicators of moderately declining plots.

### Effect of oak decline on ecological guilds

Neither the abundance nor the species richness of the community of canopy-dwelling Hymenoptera was influenced by the extent of the decline. However, contrasting effects were observed among guilds, taxonomic families and taxa (Fig. 4, Tabs. S8 & S9). For larval trophic guilds, the abundance of pollinivorous/nectarivorous and polyphagous species increased with decline severity, while those of xylophagous species decreased (Fig. 4A, Tab. S8). Koinobiont parasitoids were more abundant at intermediary levels of decline (46 % of declining trees), while the abundance of idiobiont parasitoids decreased when decline severity increased (Fig. 4A, Tab. S8). Carnivorous and polyphagous taxa also became more diverse as decline severity increased (Fig. 4B, Tab. S9). For larval habitat guilds, oak decline only slightly affected the abundance of gall-nesting taxa, while the abundance of both specialist wood-nesters and specialist soil-nesters was promoted (Fig. 4A, Tab. S8). Decline also promoted the diversity of generalist soil-, wood- and stem-nesters (Fig. 4B, Tab. S9).

**Figure 4.**
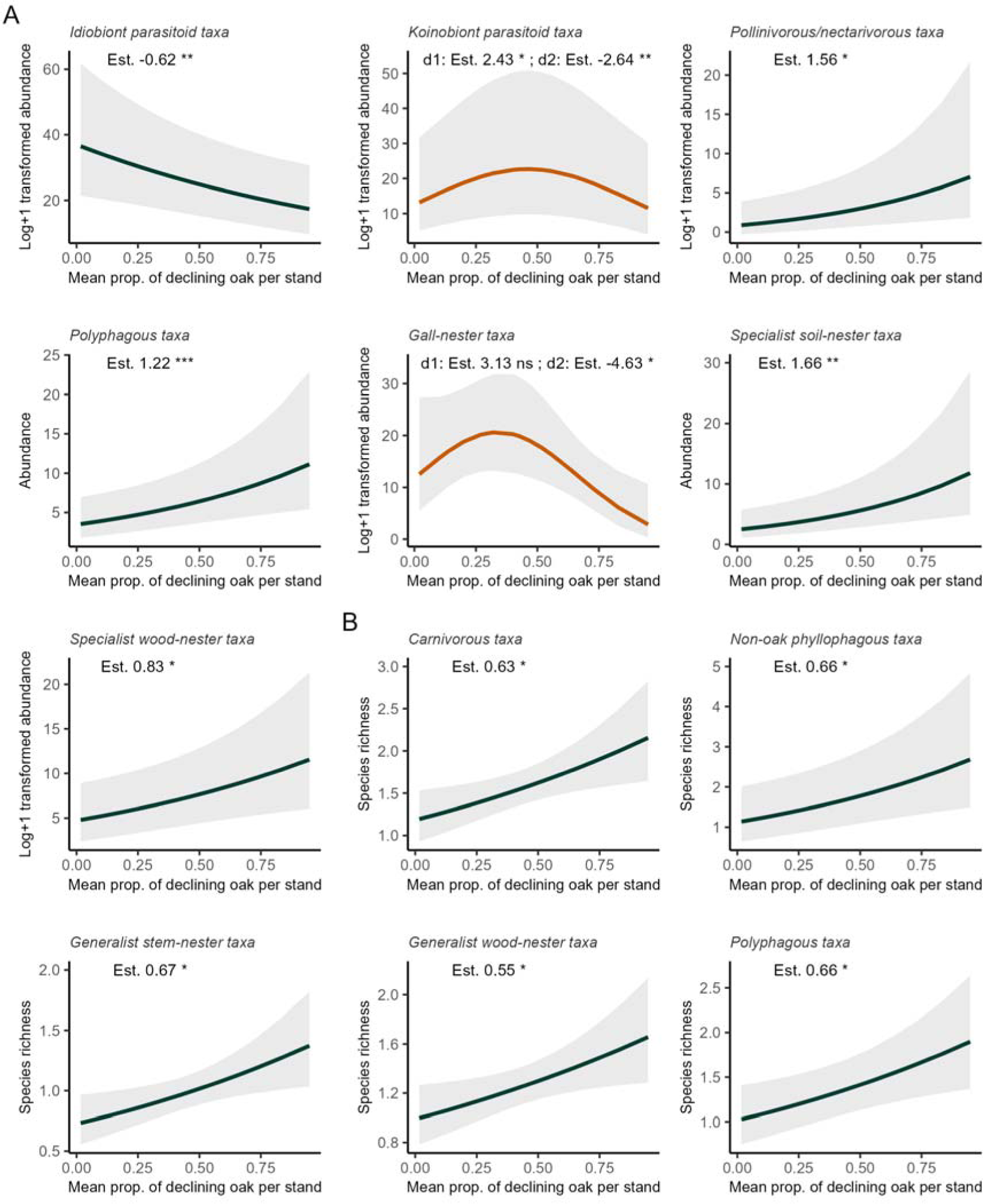
Graphical representations of the predicted linear (in dark green) or quadratic (in orange) relationships between the abundance **(A)** or species richness **(B)** of larval trophic or nesting guilds and the mean proportion (prop.) of declining oak per stand. ggeffects::ggpredict was used to predict the values. The grey area corresponds to the confidence interval for the predicted values. When the quadratic model is the best model, d1 and d2 are displayed, with d1 corresponding to the linear form of the decline and d2 to the quadratic form of the decline. Est. corresponds to the estimate and only significant relationships are shown (*** p < 0.001, ** p < 0.01, * p < 0.05). Standard error, t and z value and marginal R² are available in Tables S8 and S9.

The SEM showed that tree mortality promoted the abundance and diversity of deadwood habitats and that tree decline markedly reduced LAI (Figs. 5 & S7). Tree mortality rate negatively affected overall species richness, and both tree mortality and decline rates also had direct effects on ecological guilds (Fig. 5), and taxonomic families (Fig. S7). An increasing tree mortality rate reduced the abundance and species richness of nectarivorous/pollinivorous species, and the species richness of koinobiont parasitoids, non-oak phyllophagous, polyphagous and soil-nesting species. As mentioned above, tree decline rate negatively affected the abundance of xylophagous Hymenoptera, more specifically of the Xiphydriidae (Fig. S7), but also promoted the species richness of polyphagous species. It also promoted the abundance of Tiphiidae (Fig. S7), and a similar trend was observed for the Halictidae (*P*=0.0035). Nonetheless, the most important impact of tree decline rate was mediated by the reduction in LAI which promoted the species richness of non-oak phyllophagous species, and both the species richness and abundance of nectarivorous/pollinivorous and specialist soil-nesting taxa. LAI impacts on guilds were consistently similar to those of tree mortality, leading to opposite effects of tree decline and tree mortality on ecological guilds (Fig. 5). For the overall community, as well as for nectarivorous/pollinivorous and non-oak phyllophagous taxa, species richness consistently decreased when LAI increased, and was always low when tree mortality rate was high (Fig. S8). Finally, changes in deadwood resources, mediated by tree mortality, only marginally influenced xylophagous taxa and had no impact on the other ecological guilds (Fig. 5). Mortality or decline rates did not affect tree density, but tree density promoted the Platygastridae (Fig. S7). In line with this, it also promoted the diversity of gall-nesters (Fig. 5).

**Figure 5.**
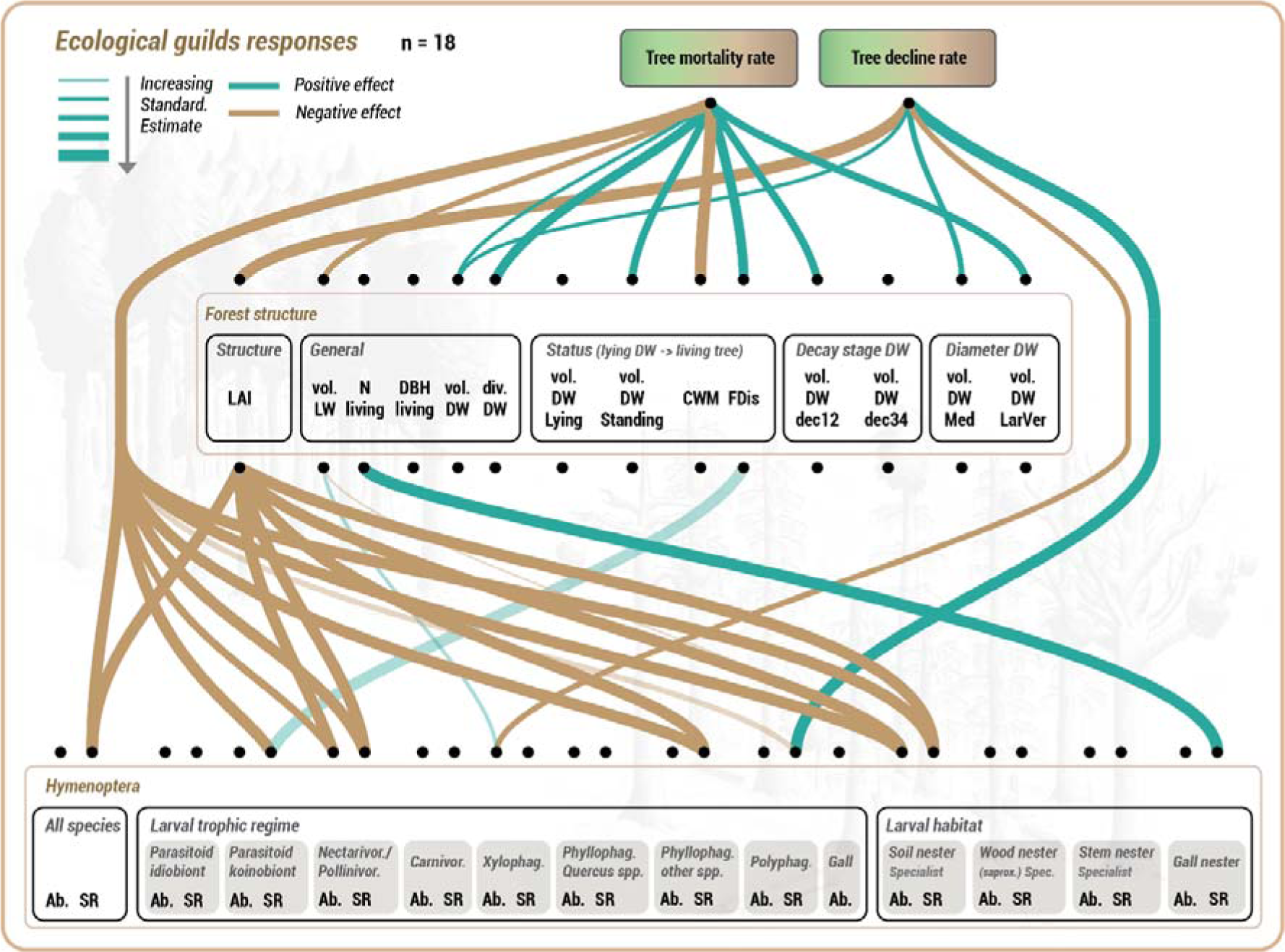
Results of the Structural Equation Modelling (SEM) for the effects of oak decline on Hymenoptera ecological guilds through changes in forest structure. Since we tested 16 predictors, we used 0.0031 as the p.value (0.05/16). The transparent lines had a p.value below 0.05 but above 0.0031 and helped to explain the response variables in the multiple regression models. LAI: leaf area index, vol.: volume, DW: deadwood, LW: living wood, N: density, DBH: diameter at breast height, CWM: community-weighted mean trait value, FDis: functional diversity, Med: medium-sized, LarVer: large and very large, dec12: low level of decay, dec34: high level of decay.

## DISCUSSION

### The community of canopy-dwelling Hymenoptera in temperate oak forests

With more than 150,000 species described worldwide, Hymenoptera are one of the most diverse orders of insects (Huber, 2017; Aberlenc, 2020). In our study, among the 19,289 specimens collected, we identified 918 taxa. As a comparison, with the same sampling design, in the same plots and at the same time, we collected approximately 97,000 Coleoptera and 10,000 Hemiptera, and identified 562 and 169 taxa respectively (unpubl. data). This indicates that the community of Hymenoptera dwelling in the canopy of the oak stands was particularly diverse. In agreement with earlier predictions (Forbes et al., 2018), these results suggest that this community could be one of the most diverse, if not the most diverse, community of arthropods in temperate oak forests.

We collected numerous phyllophagous and gall-inducing species (Tab. S2), which supports using green multi-funnel traps to collect foliage-feeding arthropods (Sallé et al., 2020). In the same traps, we also collected a wide diversity of parasitoids and predators, which might have been attracted by the green colour, indicative of potential hunting grounds for their hosts and prey. For example, the most common species in our sampling, *Perilampus ruficornis* (F.), is a hyperparasitoid that frequently parasitises leaf-chewing caterpillars and cynipids (Mitroiu and Koutsoukos, 2023). We also collected several phytophagous species that feed on oak foliage (*e.g.*, *Periclista* spp.) and oak pollen (*e.g.*, *Lasioglossum pallens* (Brullé)), and saproxylic taxa known to develop in oak branches (*i.e., Xiphydria longicollis* (Geoffroy)) or to nest in standing deadwood and/or in branches (*e.g.*, *Dolichoderus quadripunctatus* (L.)). We therefore expected to find all of these taxa in the canopy. Conversely, we found several unexpected taxa in the canopy layer. For instance, we collected species that feed on plants in the herbaceous layer (*e.g.*, *Strongylogaster multifasciata* (Geoffroy), a fern-feeder), as well as parasitoids of soil-dwelling grubs (*i.e., Tiphia femorata* F.) or spiders (*i.e., Aporus unicolor* Spinola). These may be considered as “tourist species”, without intimate relationship with oaks but which may be attracted by the microenvironments and trophic resources provided by the canopy (Moran and Southwood, 1982). We also sampled several uncommon or even rare species (*e.g.*, *Chrysis equestris* Dahlbom, *Pamphilius balteatus* (Fallén), Marhic & Noblecourt, pers. obs.), including three species of bees that are near-threatened at the European level (*i.e., Andrena fulvida* Schenck, *Lasioglossum monstrificum* (Morawitz) and *Lasioglossum sexnotatum* (Kirby); Nieto et al., 2014), and several species new for France (*e.g.*, *Spathius polonicus* Niezabitowski), or even Europe (*i.e., Plastanoxus evansi* Gorbatovsky). This highlights how little we still know about the composition and ecology of canopy-dwelling arthropods in general, and Hymenoptera in particular (Hilszczański, 2018; Sallé et al., 2021), in temperate forests.

Our sampling protocol was not initially designed to collect Hymenoptera, which are more commonly, and probably more efficiently, sampled with Malaise traps or yellow pan traps, or by fogging in the canopy (Skvarla et al., 2016; Floren et al., 2022). Although efficient in terms of diversity, our trapping design may still have underestimated the actual diversity of canopy-dwelling Hymenoptera. For instance, taxa with a low mobility might be underrepresented compared to what could be collected by fogging. It may also have biased the representativeness of our sampling in relation to the actual assemblage of canopy species. Most of our specimens were collected in the attractive green multifunnel traps. This may have led to an overrepresentation of leaf-dwelling taxa and an underestimation of floricolous or saproxylic taxa. In addition, it may have attracted species dwelling in lower strata of the ecosystem, the herbaceous and understory layers, and led to the aforementioned unexpected occurrence of these taxa in our sampling. Nonetheless, whether these species were sampled because of our trapping design or because a part of their life cycle actually takes place in the canopy is unclear at the moment since we do not have an accurate knowledge of their ecology.

### Effects of forest decline on the community, guilds and taxa of canopy-dwelling Hymenoptera

Oak decline did not directly affect the abundance and richness of the community of canopy-dwelling Hymenoptera. However, it did markedly reshape its composition, which differed significantly along the decline gradient. Species turnover was the main component of these changes, as has been commonly observed (Soininen et al., 2018), while difference among decline categories was the main contributor to global diversity. The significant contribution of nestedness to the β-diversity between moderately declining stands and both healthy and severely declining stands is congruent with the NMDS results, and indicates that the community characteristic of an intermediate level of decline is composed of taxa from the other two levels rather than of specific taxa new to the community. Similarly, in a previous study, the species richness of canopy-dwelling beetles was not influenced by oak decline, although both their abundance and their biomass increased (Sallé et al., 2020). Our result is nonetheless surprising since a recent meta-analysis indicates that forest Hymenoptera diversity (excluding ants) is generally promoted by disturbances like wildfires, windstorms and pest outbreaks (Viljur et al., 2022). However, the meta-analysis also found that the magnitude of increase in species richness was quite variable among studies (Viljur et al., 2022). In addition, these disturbances are brief and intense, while oak declines are slower and longer processes. Their impacts on the overall community of Hymenoptera might then be more progressive. In our case study, the decline effect was generally consistent between the two spatial scales considered (plot vs. stand), but the magnitude of the effects was sometimes different. This might reflect differences in foraging behaviour and habitat use among taxa and/or guilds.

Changes in community composition were driven by contrasting responses of ecological guilds to oak decline, supporting the importance of multi-taxonomic and multitrophic approaches in research on how environmental changes affect community structure (Seibold et al., 2018). As expected by our first hypothesis (H1), the abundance of pollinivorous/nectarivorous taxa increased with severity of decline. This is in line with previous studies showing that disturbances generally promote pollinators by providing more floral resources and nesting sites (Wermelinger et al., 2017; Viljur et al., 2022; Perlík et al., 2023). Interestingly, however, the dominant family of pollinivorous/nectarivorous bees, Halictidae, was mostly represented by *L. pallens*, a soil-nesting bee which mainly feeds on oak pollen and consequently, does not rely on other floral resources (Hermann et al., 2003). Since the overall abundance of specialist soil-nesters was also promoted by stand decline, this suggests that soil conditions were potentially a more important driver of community change for the guild of pollinivorous/nectarivorous taxa than were floral resources in the herbaceous layer.

Oak decline promoted the abundance, but not the diversity, of specialist wood-nesters, *i.e.,* of saproxylic taxa. Yet, the accumulation and diversification of deadwood resources and weakened hosts during forest dieback and decline events generally favour saproxylic taxa abundance and diversity (Beudert et al., 2015; Kozák et al., 2020; Cours et al., 2022, 2023), including those dwelling in the canopy (Stone et al., 2010; Sallé et al., 2020). This overall trend for the guild might be contradicted at the species level, since some saproxylic species can be negatively affected by forest dieback or decline (Vincent et al., 2020). In line with this, the only xylophagous species in our survey *i.e., X. longicollis*, was negatively affected by decline severity. The larvae of this species bore into the branches of weakened broadleaved trees and are sometimes associated with oak decline (Dominik and Starzyk, 1988). Nonetheless, severe decline conditions may have reduced the amount of living branches in the canopy, thereby reducing the amount of breeding substrates for the species, as suggested by both the direct effect of tree decline rate, and the indirect effect of tree mortality on *X. longicollis* abundance. The diversity of generalist stem-, wood- and soil-nesters was promoted by oak decline, while the species richness of the specialist guilds remained unaffected. The decline process may lead to resources pulses, which may be more readily exploited by generalist species (Devictor et al., 2008; Cours et al., 2023).

Contrary to our second hypothesis (H2), oak decline did not influence the gall-inducer taxa and phyllophagous species typically associated with oaks, and only marginally affected gall-nesters. Responses of leaf-feeding insects to tree decline can be highly variable, ranging from negative to positive effects (Martel and Mauffette, 1997; Stone et al., 2010; Sallé et al., 2020). Host specificity might modulate the outcome of tree decline on leaf-feeding species, but we did not observe any such tendency in our survey. Interestingly, oak decline had an overall negative effect on the parasitoids of defoliating or mining caterpillars. This suggests a negative impact of decline on leaf-feeding moths, with potential cascading impacts on higher trophic levels. Conversely, the diversity of leaf-feeders associated with other host plants increased with decline severity, and several of them were indicators of high levels of decline (*i.e., Strongylogaster multifasciata* (Geoffroy) and *Aneugmenus padi* (L.), both fern-feeders; *Tenthredopsis nassata* (L.) and *Dolerus gonager* (F.), bentgrass-feeders; *Eutomostethus luteiventris* (Klug), a rush-feeder; and *Pachyprotasis simulans* (Klug), a goldenrod-feeder (Lacourt, 2020)). All of the above were “Symphyta” and their increase might reflect the preference of these wasps for open forests (Lehnert et al., 2013), but it might also result from a cascading effect of an increase in plant diversity and biomass along with increased canopy openness (Lehnert et al., 2013; Lu et al., 2019; Cacciatori et al., 2022).

As expected from the literature (Gaston, 1991), the parasitoid guild was both the most diverse and the most abundant guild. Since parasitoids, and especially koinobionts, require a diversity of habitats to support the full assemblage of their hosts, they are highly sensitive to changes in habitat (Fraser et al., 2007; Hilszczański, 2018; Jonsell et al., 2023). As anticipated by our third hypothesis (H3), koinobiont parasitoids exhibited a quadratic relationship with decline levels, and were more abundant at intermediate levels of decline severity. This suggests that a moderate decline may provide a greater diversity of habitats and trophic resources for specialist parasitoids, as predicted, for instance, by the intermediate disturbance hypothesis (Grime, 1973; Horn, 1975). Idiobiont parasitoids, however, where consistently negatively affected by decline severity. This was unexpected since they are considered as generalists in terms of host range, and generalist species generally thrive in disturbed ecosystems (*e.g.*, Devictor et al., 2008). These overall trends might be further modulated though by the host type of the parasitoid. We observed that both Tiphiidae (*e.g.*, *T. femorata*) and Evanidae (*e.g.*, *Brachygaster minuta* (Olivier), which parasitize chafers and cockroaches respectively, benefited from tree decline, while the parasitoids of defoliating or mining moths (*e.g.*, *Agrypon flaveolatum* (Gravenhorst), *Bassus* sp. 1, *Earinus gloriatorius* (Panzer), *Macrocentrus nitidus* (Wesmael), *P. ruficornis* (Mitroiu and Koutsoukos, 2023)), were somewhat disadvantaged by high levels of decline. Further studies would be necessary to assess whether these effects depend on host type or are idiosyncratic species responses. The other guild belonging to a higher trophic level, the carnivorous taxa, was promoted by oak decline. Disturbances can have highly contrasted effects on predators (Cours et al., 2023). In our study, this guild included several taxonomic families such as Vespidae, Formicidae and Pemphredonidae that prey on a wide range of taxa. The positive decline effect might result either from a higher abundance or accessibility of prey (Cours et al., 2023), or from better habitat conditions. Further studies would be required to identify the factors underpinning this positive effect. The abundance and the diversity of polyphagous species were also enhanced by oak decline. This is congruent with the fact that ants, which dominated the guild of polyphagous species, are known to benefit from silvicultural practices leading to more open conditions (Tausan et al., 2017; Grevé et al., 2018; Cours et al., 2023).

### Key parameters of decline-driven changes in Hymenoptera guilds

The results of the SEMs were consistent with those of the GLMMs, indicating that the effects highlighted by the SEM approach were probably also involved in hymenopteran responses to oak decline over the whole sampling design. Changes in community composition and structure resulted either from the direct effects of tree mortality and tree decline, or from indirect effects mediated by alterations of the forest structure. As dead branches accumulated in the crown of declining trees, an increase in tree decline rate markedly reduced the LAI. Most of the tree decline effects on ecological guilds were mediated by changes in LAI, which was therefore a major driver of community change. In parallel, an increase in tree mortality rate increased the volume and diversity of deadwood resources.

An increasing tree decline rate directly promoted the abundance of Halictidae and Tiphiidae (Figs. 5, S7). Both families rely on ground-related trophic resources (floral resources for the Halictidae and soil-dwelling larvae for the Tiphiidae) and ground habitats since they are soil-nesters (Michez et al., 2019). In declining stands, soil-nesters may benefit from better or warmer micro-environmental conditions but also from better underground resources due to LAI reduction (Cours et al., 2023). In line with this, generalist leaf-feeding weevils, with root-feeding larvae, have been shown to benefit from oak decline (Sallé et al., 2020). The positive effect of a reduction in LAI on nectarivorous/pollinivorous taxa is also congruent with its positive effect on the diversity of non-oak phyllophagous taxa, supporting the above-mentioned hypothesis of a pulse in plant diversity and floral resources when canopy openness increases (Romey et al., 2007; Lu et al., 2019; Cacciatori et al., 2022). In our study, canopy opening promoted the overall diversity of the Hymenoptera taxa, which supports previous observations (Eckerter et al., 2022; Perlík et al., 2023; Rappa et al., 2023). This result also indicates that micro-environmental modifications at the ground level can have critical impacts on the communities dwelling in higher forest strata, especially when species have developmental instars that rely on ground resources or ground micro-habitats.

Surprisingly, while tree mortality rate altered a large array of deadwood resources, we did not observe any effect of these alterations on the taxonomic families and guilds of Hymenoptera, aside from the negative effect of a reduction in the number of living trees on the xylophagous Xiphydriidae. This contradicts several studies where the accumulation and/or diversification of deadwood resources promoted the abundance, species richness or community composition of different Hymenoptera guilds and taxa like bees and wasps (Bogusch and Horák, 2018; Urban-Mead et al., 2021; Rappa et al., 2023), and parasitoid wasps (Hilszczański et al., 2005; Ulyshen et al., 2011a). However, our results are congruent with recent observations where changes in deadwood amounts did not influence the species richness or community composition of either cavity- or non-cavity-nesting Hymenoptera, at least at a local scale (Perlík et al., 2023). Our study plots were located in managed oak forests, and active forest management likely restricted the magnitude of accumulation and diversification of deadwood resources (Bouget et al., 2023). This may have limited the potentially beneficial effects of tree mortality and decline on saproxylic Hymenoptera, thus only weakly promoting saproxylic taxa and wood-nesters. It may also be a matter of spatial scale; considering the amount and diversity of deadwood resources over larger spatial scales might be more relevant, especially for taxa with a high dispersal capacity (Cours et al., 2022). In addition, considering more precise parasitoid subguilds, especially those with saproxylic hosts, might also help to unravel the impacts of changes in deadwood on the forest Hymenoptera community.

In our study, the effects of tree mortality rate on the ecological guilds were consistently opposed to those of the decline rate, and were, overall, negative. The stands with a high tree mortality rate might very well have experienced more severe and frequent droughts in the years before the study, leading to pervasive cascading effects on Hymenoptera through alterations in plant-pollinator or host-parasitoid interactions, for instance (Rouault et al., 2006; Endres et al., 2021). The negative effects could also indicate that canopies with dead trees are avoided by Hymenoptera, thus leading to an overall reduction in abundance and species richness. Stands with high levels of tree mortality might also have experienced critical changes in micro-environmental conditions and trophic resources, but further studies are required to better understand the processes underpinning these deleterious effects.

## CONCLUSION

Our results highlight the extreme diversity of the forest Hymenoptera community, and the relevance of Hymenoptera in studying changes in global community structure and forest functional processes. Our results also support their value as forest bioindicators, as previously proposed by Maleque *et al*. (2009). Nonetheless, our study also underlines the relative lack of knowledge of this group in temperate forests (Hilszczański, 2018; Jonsell et al., 2023). Consequently, considering the growing threats to biodiversity in general, we call for thorough investigations on the diversity and ecology of Hymenoptera to make better use of their potential as bioindicators, understand their ecological services, and better evaluate their conservation value. This suggests that micro-environmental changes at the ground level strongly influence changes in canopy communities. Consequently, we also call for a thorough investigation of the changes in ecological processes and community structure at the ground level following forest decline and dieback.

Oak decline had mixed effects on canopy-dwelling Hymenoptera in our study. We found contrasting responses for some taxa and ecological guilds, which profoundly reshaped the species assemblage but did not affect the overall abundance or diversity of the community. Although tree mortality rate induced local changes in deadwood resources, it only had negative impacts on the guilds and taxonomic families considered in our study, which is in line with the negative or quadratic relationships frequently observed in our analyses. This suggests that late stages of the decline process, and/or severe declines or diebacks, would be detrimental for several ecological guilds of canopy-dwelling arthropods. When dead trees accumulate, only scarce fragments of the canopy remain, and the ecosystem shifts towards open habitats, where taxa dependant on ground-related resources and microhabitats (floral resources, herbaceous layer and soil conditions) can thrive, leading to a major shift in forest insect communities. We have shown that the β-diversity among decline categories at the stand level significantly contributed to the overall γ-diversity, mostly through species turnover. This suggests that, when possible, maintaining a landscape mosaic including stands at different levels of decline or dieback, and/or stands with different levels of canopy closure, may serve a conservation purpose (Socolar et al., 2016), especially since maintaining high habitat heterogeneity is generally a key goal for the conservation of insect communities (Samways, 2015).

## ACKNOWLEDGEMENTS

We thank C. Moliard (INRAE), G. Parmain (INRAE), X. Pineau (P2e) and O. Denux (INRAE) for their technical assistance. We also thank T. Wood, S. Flaminio and P. Rosa for their help in determining the Andrenidae, Halictidae and Chrysididae. We are grateful to the National Forest Office (Office National des Forêts) for their field assistance. We would like to thank Vicky Moore for her English proofreading and the two anonymous reviewers for their comments, which helped to improve this manuscript considerably. This work was supported by Région Centre-Val de Loire Project no. 2018-00124136 (CANOPEE) coordinated by A. Sallé.

## SUPPLEMENTARY INFORMATION

Representative specimens of Braconidae, Ichneumonidae, Halictidae, Andrenidae, Crabronidae and Pemphredonidae are housed at the Orleans Museum for Biodiversity and the Environment (MOBE, Orléans, France). R scripts, the list of species traits and the list of plot and stand characteristics can be found at: https://doi.org/10.57745/H8CBG1

