## supplementary tables and figures for "Responses of the hyper-diverse community of canopy-dwelling Hymenoptera to oak decline"

**SUPPLEMENTARY DATA - TABLES**

Table S1. Stand and plot locations and characteristics. *Q: Quercus, P: Pinus, F: Fagus, C: Carpinus, Po: Populus, T: Tilia, Ca: Castanea, So: Sorbus, B: Betulus, Sa: Salix, Fr: Frangula.*

| **Forest** | **Stand** | **Plot** | **Long.** | **Lat.** | **Mean plot decline rate** | **Mean stand decline rate** | **Mean stand height (m)** | **Mean stand DBH (cm) ± SE** | **Stand tree density (tree/ha)** | **Oak basal area (m²/ha)** | **Stand composition (dominant level)** | **Stand composition (understory)** |
| --- | --- | --- | --- | --- | --- | --- | --- | --- | --- | --- | --- | --- |
| Marcenat | 12 | 12_1 | 3.36082 | 46.24709 | 0.6 | 0.56 | 29 | 56 ± 6 | 154 | 19 | *Q* | *C* |
| Marcenat | 12 | 12_3 | 3.36194 | 46.24607 | 0.48 | 0.56 | 29 | 56 ± 6 | 154 | 19 | *Q* | *C* |
| Vierzon | 19 | 19_1 | 2.18129 | 47.25792 | 1.00 | 0.95 | 26 | 57 ± 3 | 87 | 18 | *Q, P* | *C* |
| Vierzon | 19 | 19_3 | 2.18125 | 47.25891 | 0.85 | 0.95 | 26 | 57 ± 3 | 87 | 18 | *Q, P* | *C* |
| Vierzon | 35 | 35_1 | 2.17526 | 47.29789 | 0.2 | 0.43 | 26 | 53 ± 3 | 105 | 21 | *Q, P, F* | *C* |
| Vierzon | 35 | 35_3 | 2.17310 | 47.29719 | 0.8 | 0.43 | 26 | 53 ± 3 | 105 | 21 | *Q, P, F* | *C* |
| Marcenat | 37 | 37_1 | 3.37381 | 46.23167 | 0.15 | 0.2 | 37 | 79 ± 4 | 76 | 18 | *Q* | *C, T* |
| Marcenat | 37 | 37_3 | 3.37339 | 46.22902 | 0.3 | 0.2 | 37 | 79 ± 4 | 76 | 18 | *Q* | *C, T* |
| Marcenat | 58 | 58_1 | 3.35888 | 46.20639 | 0.2 | 0.52 | 23 | 40 ± 4 | 389 | 33 | *Q* | *C* |
| Marcenat | 58 | 58_3 | 3.36050 | 46.20959 | 0.8 | 0.52 | 23 | 40 ± 4 | 389 | 33 | *Q* | *C* |
| Vierzon | 70 | 70_1 | 2.15505 | 47.29036 | 0.65 | 0.83 | 24 | 50 ± 6 | 58 | 9 | *Q, P, F* | *P* |
| Vierzon | 70 | 70_2 | 2.15439 | 47.28989 | 0.95 | 0.83 | 24 | 50 ± 6 | 58 | 9 | *Q, P, F* | *P* |
| Vierzon | 71 | 71_1 | 2.15458 | 47.28711 | 0.45 | 0.68 | 25 | 47 ± 3 | 105 | 15 | *Q, P, F* | *F* |
| Vierzon | 71 | 71_2 | 2.15554 | 47.28721 | 0.7 | 0.68 | 25 | 47 ± 3 | 105 | 15 | *Q, P, F* | *F* |
| Vierzon | 81 | 81_1 | 2.19792 | 47.26712 | 0.075 | 0.14 | 24 | 43 ± 2 | 197 | 15 | *Q* | *C* |
| Vierzon | 81 | 81_2 | 2.19679 | 47.26614 | 0.05 | 0.14 | 24 | 43 ± 2 | 197 | 15 | *Q* | *C* |
| Vierzon | 179 | 179_1 | 2.11877 | 47.26536 | 0.575 | 0.65 | 29 | 64 ± 6 | 81 | 21 | *Q* |  |
| Vierzon | 179 | 179_3 | 2.11811 | 47.26589 | 0.55 | 0.65 | 29 | 64 ± 6 | 81 | 21 | *Q* |  |
| Vierzon | 236 | 236_1 | 2.08074 | 47.26758 | 0.4 | 0.43 | 24 | 73 ± 4 | 88 | 23 | *Q, F* | *C, F, B* |
| Vierzon | 236 | 236_3 | 2.07986 | 47.27009 | 0.65 | 0.43 | 24 | 73 ± 4 | 88 | 23 | *Q, F* | *C, F, B* |
| Vierzon | 249 | 249_1 | 2.06739 | 47.26554 | 0.65 | 0.64 | 25 | 52 ± 2 | 127 | 22 | *Q, P, C* |  |
| Vierzon | 249 | 249_2 | 2.06770 | 47.26462 | 0.41 | 0.64 | 25 | 52 ± 2 | 127 | 22 | *Q, P, C* |  |
| Vierzon | 290 | 290_1 | 2.03273 | 47.26114 | 0.075 | 0.32 | 24 | 50 ± 2 | 160 | 22 | *Q* | *So* |
| Vierzon | 290 | 290_2 | 2.03277 | 47.26007 | 0.45 | 0.32 | 24 | 50 ± 2 | 160 | 22 | *Q* | *So* |
| Orléans | 351 | 351_1 | 2.46153 | 47.86942 | 0.05 | 0.02 | 24 | 63 ± 5 | 136 | 18 | *Q, P* | *Ca, So* |
| Orléans | 351 | 351_2 | 2.46296 | 47.86943 | 0,00 | 0.03 | 24 | 63 ± 5 | 136 | 18 | *Q, P* | *Ca, So* |
| Orléans | 751 | 751_1 | 2.29365 | 47.98251 | 0.5 | 0.40 | 27 | 64 ± 4 | 107 | 24 | *Q, F* | *P, C* |
| Orléans | 751 | 751_2 | 2.29409 | 47.98275 | 0.4 | 0.41 | 27 | 64 ± 4 | 107 | 24 | *Q, F* | *P, C* |
| Orléans | 1140 | 1140_1 | 2.18281 | 48.03878 | 0.75 | 0.73 | 25 | 72 ± 6 | 90 | 26 | *Q* | *Fr* |
| Orléans | 1140 | 1140_2 | 2.18181 | 48.039 | 0.75 | 0.74 | 25 | 72 ± 6 | 90 | 26 | *Q* | *Fr* |
| Orléans | 1343 | 1343_1 | 1.98182 | 48.00018 | 0,00 | 0.02 | 28 | 75 ± 4 | 100 | 28 | *Q* | *C* |
| Orléans | 1343 | 1343_2 | 1.98265 | 47.99997 | 0,00 | 0.03 | 28 | 75 ± 4 | 100 | 28 | *Q* | *C* |
| Orléans | 1344 | 1344_1 | 1.97403 | 47.99847 | 0.3 | 0.47 | 24 | 78 ± 5 | 74 | 27 | *Q* | *C* |
| Orléans | 1344 | 1344_2 | 1.97383 | 47.99771 | 0.4 | 0.48 | 24 | 78 ± 5 | 74 | 27 | *Q* | *C* |
| Orléans | 1427 | 1427_2 | 1.95332 | 47.94775 | 0.5 | 0.52 | 26 |  | 140 | 15 | *Q, P* | *C, Sa* |
| Orléans | 1427 | 1427_3 | 1.95451 | 47.94805 | 0.1 | 0.53 | 26 |  | 140 | 15 | *Q, P* | *C, Sa* |
| Orléans | 1490 | 1490_1 | 1.90605 | 48.01824 | 0.15 | 0.1 | 23 | 70 ± 9 | 73 | 17 | *Q, Po* | *C* |
| Orléans | 1490 | 1490_3 | 1.90521 | 48.01915 | 0.15 | 0.1 | 23 | 70 ± 9 | 73 | 17 | *Q, Po* | *C* |
| Orléans | 1491 | 1491_1 | 1.90963 | 48.01389 | 0.2 | 0.1 | 23 | 65 ± 3 | 90 | 17 | *Q, Po* | *C* |
| Orléans | 1491 | 1491_2 | 1.90888 | 48.01383 | 0.1 | 0.1 | 23 | 65 ± 3 | 90 | 17 | *Q, Po* | *C* |
| Orléans | 1502 | 1502_1 | 1.90291 | 48.01353 | 0.5 | 0.27 | 21 | 48 ± 3 | 222 | 16 | *Q, Po* | *C* |
| Orléans | 1502 | 1502_3 | 1.90158 | 48.01330 | 0.15 | 0.28 | 21 | 48 ± 3 | 222 | 16 | *Q, Po* | *C* |

Table S2. List of taxa collected in 2019 in the canopy of three oak forests in France. Larval trophic guild: carnivorous (Carn.), idiobiont parasitoid (Par. I.), koinobiont parasitoid (Par. K.), phyllophagous feeding on oaks (O. Phyll.), phyllophagous feeding on other host plants (N. O. Phyll.), phytophagous (Phyt.), pollinivorous / nectarivorous (Poll./Nec.), polyphagous (Poly.), xylophagous (Xyl.). Nesting site: soil (So.), wood (Wo.), stems (St.), galls (Ga.). IUCN status: ERL corresponds to the European Red List and WRL to the Worldwide Red List. Abund.: abundance. Species in bold are new to France.

| **Family** | **Taxa** | **Larval trophic guild** | **Nesting site** | | | | **IUCN**  **(2022)** | **Abund.** | **Identifier** |
| --- | --- | --- | --- | --- | --- | --- | --- | --- | --- |
|  |  |  | **So.** | **Wo.** | **St.** | **Ga.** |  |  |  |
| Ampulicidae | *Ampulex fasciata Jurine, 1807* | Carn. |  |  |  |  |  | 4 | P. Burguet |
| Ampulicidae | *Dolichurus corniculus*  *(Spinola, 1808)* | Carn. |  |  |  |  |  | 2 | F. Herbrecht |
| Andrenidae | *Andrena afzeliella (Kirby, 1802)* | Poll./Nec. |  |  |  |  |  | 2 | T. Wood |
| Andrenidae | *Andrena angustior (Kirby, 1802)* | Poll./Nec. |  |  |  |  | ERL (DD) | 1 | T. Wood |
| Andrenidae | *Andrena bicolor Fabricius, 1775* | Poll./Nec. |  |  |  |  | ERL (LC) | 2 | T. Wood |
| Andrenidae | *Andrena bimaculata*  *(Kirby, 1802)* | Poll./Nec. |  |  |  |  | ERL (DD) | 1 | T. Wood |
| Andrenidae | *Andrena chrysosceles*  *(Kirby, 1802)* | Poll./Nec. |  |  |  |  | ERL (DD) | 5 | T. Wood |
| Andrenidae | *Andrena cinerea Brullé, 1832* | Poll./Nec. |  |  |  |  | ERL (DD) | 8 | T. Wood |
| Andrenidae | *Andrena dorsata (Kirby, 1802)* | Poll./Nec. |  |  |  |  | ERL (DD) | 13 | T. Wood |
| Andrenidae | *Andrena ferox Smith, 1847* | Poll./Nec. |  |  |  |  | ERL (DD) | 50 | T. Wood |
| Andrenidae | *Andrena flavipes Panzer, 1799* | Poll./Nec. |  |  |  |  | ERL (LC) | 3 | T. Wood |
| Andrenidae | *Andrena fulva (Müller, 1766)* | Poll./Nec. |  |  |  |  | ERL (DD) | 8 | T. Wood |
| Andrenidae | *Andrena fulvida Schenck, 1853* | Poll./Nec. |  |  |  |  | ERL (NT) | 4 | T. Wood |
| Andrenidae | *Andrena haemorrhoa*  *(Fabricius, 1781)* | Poll./Nec. |  |  |  |  | ERL (LC) | 21 | T. Wood |
| Andrenidae | *Andrena helvola*  *(Linnaeus, 1758)* | Poll./Nec. |  |  |  |  | ERL (DD) | 4 | T. Wood |
| Andrenidae | *Andrena minutula (Kirby, 1802)* | Poll./Nec. |  |  |  |  | ERL (DD) | 8 | T. Wood |
| Andrenidae | *Andrena minutuloides*  *Perkins, 1914* | Poll./Nec. |  |  |  |  | ERL (DD) | 2 | T. Wood |
| Andrenidae | *Andrena mitis*  *Schmiedeknecht, 1883* | Poll./Nec. |  |  |  |  | ERL (DD) | 4 | T. Wood |
| Andrenidae | *Andrena nigroaenea*  *(Kirby, 1802)* | Poll./Nec. |  |  |  |  | ERL (LC) | 6 | T. Wood |
| Andrenidae | *Andrena nitida (Müller, 1776)* | Poll./Nec. |  |  |  |  | ERL (LC) | 5 | T. Wood |
| Andrenidae | *Andrena propinqua*  *Schenck, 1853* | Poll./Nec. |  |  |  |  | ERL (DD) | 1 | T. Wood |
| Andrenidae | *Andrena scotica Perkins, 1916* | Poll./Nec. |  |  |  |  |  | 18 | T. Wood |
| Andrenidae | *Andrena strohmella*  *Stoeckhert, 1928* | Poll./Nec. |  |  |  |  | ERL (LC) | 4 | T. Wood |
| Andrenidae | *Andrena subopaca*  *Nylander, 1848* | Poll./Nec. |  |  |  |  | ERL (LC) | 3 | T. Wood |
| Andrenidae | *Andrena sulcata Donovan, 1977* | Poll./Nec. |  |  |  |  |  | 10 | T. Wood |
| Andrenidae | Andrenidae sp. | Poll./Nec. |  |  |  |  |  | 30 | LBLGC team |
| Apidae | *Apis mellifera Linnaeus, 1758* | Poll./Nec. |  |  |  |  | ERL (DD) | 32 | LBLGC team |
| Apidae | *Bombus gr. terrestris* | Poll./Nec. |  |  |  |  |  | 21 | D. Michez |
| Apidae | *Bombus (Psithyrus) sp.* | Poll./Nec. |  |  |  |  |  | 1 | LBLGC team |
| Apidae | *Nomada sp.* | Poll./Nec. |  |  |  |  |  | 3 | LBLGC team |
| Argidae | *Arge rustica (Linnaeus, 1758)* | O. Phyll. |  |  |  |  |  | 3 | T. Noblecourt |
| Argidae | *Sterictiphora longicornis*  *Chevin, 1982* | N.O. Phyll. |  |  |  |  |  | 2 | T. Noblecourt |
| Bembicidae | *Argogorytes mystaceus (Linnaeus, 1760)* | Carn. |  |  |  |  |  | 1 | P. Burguet |
| Bembicidae | *Gorytes quinquecinctus (Fabricius, 1793)* | Carn. |  |  |  |  |  | 1 | P. Burguet |
| Bembicidae | *Lestiphorus bicinctus*  *(Rossi, 1794)* | Carn. |  |  |  |  |  | 1 | P. Burguet |
| Bethylidae | *Bethylus boops (Thomson, 1861)* | Par. I. |  |  |  |  |  | 4 | E. Marhic |
| Bethylidae | *Bethylus dendrophilus*  *Richards, 1939* | Par. I. |  |  |  |  |  | 5 | E. Marhic |
| Bethylidae | *Bethylus fuscicornis*  *(Jurine, 1807)* | Par. I. |  |  |  |  |  | 3 | E. Marhic |
| Bethylidae | *Epyris cf. bilineatus* | Par. I. |  |  |  |  |  | 8 | E. Marhic |
| Bethylidae | *Epyris niger  Westwood, 1832* | Par. I. |  |  |  |  |  | 7 | E. Marhic |
| Bethylidae | *Laelius femoralis*  *(Foerster, 1860)* | Par. I. |  |  |  |  |  | 6 | E. Marhic |
| Bethylidae | *Parascleroderma sulcatifrons*  *(Kieffer, 1908)* | Par. I. |  |  |  |  |  | 1 | E. Marhic |
| Bethylidae | ***Plastanoxus evansi Gorbatovsky, 1995*** | Par. I. |  |  |  |  |  | 1 | E. Marhic, J. De Rond |
| Braconidae | *Acampsis alternipes*  *(Nees, 1816)* | Par. K. |  |  |  |  |  | 4 | Y. Braet |
| Braconidae | *Ascogaster annularis*  *(Nees, 1816)* | Par. K. |  |  |  |  |  | 1 | Y. Braet |
| Braconidae | *Ascogaster armata*  *Wesmael, 1835* | Par. K. |  |  |  |  |  | 1 | Y. Braet |
| Braconidae | *Ascogaster exigua*  *Huddleston, 1984* | Par. K. |  |  |  |  |  | 2 | Y. Braet |
| Braconidae | *Ascogaster sp. 1* | Par. K. |  |  |  |  |  | 1 | Y. Braet |
| Braconidae | *Ascogaster sp. 2* | Par. K. |  |  |  |  |  | 1 | Y. Braet |
| Braconidae | *Ascogaster varipes*  *Wesmael, 1835* | Par. K. |  |  |  |  |  | 16 | Y. Braet |
| Braconidae | *Aspicolpus sp.* | Par. K. |  |  |  |  |  | 2 | Y. Braet |
| Braconidae | *Bassus sp. 1* | Par. K. |  |  |  |  |  | 38 | Y. Braet |
| Braconidae | *Bassus sp. 2* | Par. K. |  |  |  |  |  | 22 | Y. Braet |
| Braconidae | *Bassus sp. 3* | Par. K. |  |  |  |  |  | 1 | Y. Braet |
| Braconidae | *Bassus sp. 4* | Par. K. |  |  |  |  |  | 2 | Y. Braet |
| Braconidae | Blacinae sp. 1 | Par. K. |  |  |  |  |  | 1 | Y. Braet |
| Braconidae | Blacinae sp. 2 | Par. K. |  |  |  |  |  | 1 | Y. Braet |
| Braconidae | Blacinae sp. 3 | Par. K. |  |  |  |  |  | 1 | Y. Braet |
| Braconidae | Blacinae sp. 4 | Par. K. |  |  |  |  |  | 1 | Y. Braet |
| Braconidae | Blacinae sp. 5 | Par. K. |  |  |  |  |  | 1 | Y. Braet |
| Braconidae | Blacinae sp. 6 | Par. K. |  |  |  |  |  | 1 | Y. Braet |
| Braconidae | Blacinae sp. 7 | Par. K. |  |  |  |  |  | 1 | Y. Braet |
| Braconidae | Blacinae sp. 8 | Par. K. |  |  |  |  |  | 1 | Y. Braet |
| Braconidae | Blacinae sp. 9 | Par. K. |  |  |  |  |  | 1 | Y. Braet |
| Braconidae | Braconidae sp. | Par. |  |  |  |  |  | 4 | LBLGC team |
| Braconidae | Braconinae sp. 1 | Par. I. |  |  |  |  |  | 1 | Y. Braet |
| Braconidae | Braconinae sp. 2 | Par. I. |  |  |  |  |  | 1 | Y. Braet |
| Braconidae | Braconinae sp. 3 | Par. I. |  |  |  |  |  | 1 | Y. Braet |
| Braconidae | Braconinae sp. 4 | Par. I. |  |  |  |  |  | 1 | Y. Braet |
| Braconidae | Braconinae sp. 5 | Par. I. |  |  |  |  |  | 2 | Y. Braet |
| Braconidae | Braconinae sp. 6 | Par. I. |  |  |  |  |  | 2 | Y. Braet |
| Braconidae | Braconinae sp. 7 | Par. I. |  |  |  |  |  | 1 | Y. Braet |
| Braconidae | Braconinae sp. 8 | Par. I. |  |  |  |  |  | 1 | Y. Braet |
| Braconidae | *Chrysopophthorus hungaricus*  *(Zilahi-Kiss, 1927)* | Par. K. |  |  |  |  |  | 2 | Y. Braet |
| Braconidae | *Dendrosoter protuberans*  *(Nees, 1834)* | Par. I. |  |  |  |  |  | 3 | Y. Braet |
| Braconidae | *Diospilus sp. 1* | Par. K. |  |  |  |  |  | 9 | Y. Braet |
| Braconidae | *Diospilus sp. 2* | Par. K. |  |  |  |  |  | 1 | Y. Braet |
| Braconidae | *Diospilus sp. 3* | Par. K. |  |  |  |  |  | 1 | Y. Braet |
| Braconidae | *Diospilus sp. 4* | Par. K. |  |  |  |  |  | 23 | Y. Braet |
| Braconidae | *Diospilus sp. 5* | Par. K. |  |  |  |  |  | 1 | Y. Braet |
| Braconidae | *Doryctes aff. mutillator* | Par. I. |  |  |  |  |  | 3 | Y. Braet |
| Braconidae | *Doryctes leucogaster*  *(Nees, 1834)* | Par. I. |  |  |  |  |  | 7 | Y. Braet |
| Braconidae | Doryctinae sp. 1 | Par. I. |  |  |  |  |  | 1 | Y. Braet |
| Braconidae | Doryctinae sp. 2 | Par. I. |  |  |  |  |  | 1 | Y. Braet |
| Braconidae | Doryctinae sp. 3 | Par. I. |  |  |  |  |  | 1 | Y. Braet |
| Braconidae | *Earinus elator  (Fabricius, 1804)* | Par. K. |  |  |  |  |  | 180 | Y. Braet |
| Braconidae | *Earinus gloriatorius*  *(Panzer, 1809)* | Par. K. |  |  |  |  |  | 43 | Y. Braet |
| Braconidae | *Eubazus sp. 1* | Par. K. |  |  |  |  |  | 4 | Y. Braet |
| Braconidae | *Eubazus sp. 2* | Par. K. |  |  |  |  |  | 2 | Y. Braet |
| Braconidae | *Eubazus sp. 3* | Par. K. |  |  |  |  |  | 1 | Y. Braet |
| Braconidae | *Eubazus sp. 4* | Par. K. |  |  |  |  |  | 2 | Y. Braet |
| Braconidae | *Eubazus sp. 5* | Par. K. |  |  |  |  |  | 8 | Y. Braet |
| Braconidae | *Eubazus sp. 6* | Par. K. |  |  |  |  |  | 1 | Y. Braet |
| Braconidae | *Eubazus sp. 7* | Par. K. |  |  |  |  |  | 2 | Y. Braet |
| Braconidae | *Eubazus sp. 8* | Par. K. |  |  |  |  |  | 5 | Y. Braet |
| Braconidae | *Gnamptodon pilosus*  *(van Achterberg, 1983)* | Par. K. |  |  |  |  |  | 2 | Y. Braet |
| Braconidae | *Gnamptodon pumilio*  *(Nees, 1834)* | Par. K. |  |  |  |  |  | 1 | Y. Braet |
| Braconidae | *Hecabolus sulcatus Curtis, 1834* | Par. I. |  |  |  |  |  | 1 | Y. Braet |
| Braconidae | *Helcon claviventris*  *Wesmael, 1835* | Par. K. |  |  |  |  |  | 6 | Y. Braet |
| Braconidae | *Helcon tardator Nees, 1812* | Par. K. |  |  |  |  |  | 8 | Y. Braet |
| Braconidae | *Homolobus flagitator*  *(Curtis, 1837)* | Par. K. |  |  |  |  |  | 1 | Y. Braet |
| Braconidae | *Macrocentrus bicolor*  *Curtis, 1833* | Par. K. |  |  |  |  |  | 9 | Y. Braet |
| Braconidae | *Macrocentrus flavus*  *Snellen van Vollenhoven, 1878* | Par. K. |  |  |  |  |  | 3 | Y. Braet |
| Braconidae | *Macrocentrus marginator*  *(Nees, 1811)* | Par. K. |  |  |  |  |  | 3 | Y. Braet |
| Braconidae | *Macrocentrus nitidus*  *(Wesmael, 1835)* | Par. K. |  |  |  |  |  | 42 | Y. Braet |
| Braconidae | *Meteorus aff. longipilosus* | Par. K. |  |  |  |  |  | 18 | Y. Braet |
| Braconidae | *Meteorus sp.1* | Par. K. |  |  |  |  |  | 1 | Y. Braet |
| Braconidae | *Meteorus sp.2* | Par. K. |  |  |  |  |  | 2 | Y. Braet |
| Braconidae | *Meteorus sp.3* | Par. K. |  |  |  |  |  | 1 | Y. Braet |
| Braconidae | *Meteorus sp.4* | Par. K. |  |  |  |  |  | 1 | Y. Braet |
| Braconidae | *Meteorus sp.5* | Par. K. |  |  |  |  |  | 2 | Y. Braet |
| Braconidae | Metopiinae sp. 1 | Par. K. |  |  |  |  |  | 2 | Y. Braet |
| Braconidae | Metopiinae sp. 10 | Par. K. |  |  |  |  |  | 2 | Y. Braet |
| Braconidae | Metopiinae sp. 11 | Par. K. |  |  |  |  |  | 2 | Y. Braet |
| Braconidae | Metopiinae sp. 12 | Par. K. |  |  |  |  |  | 2 | Y. Braet |
| Braconidae | Metopiinae sp. 13 | Par. K. |  |  |  |  |  | 7 | Y. Braet |
| Braconidae | Metopiinae sp. 14 | Par. K. |  |  |  |  |  | 1 | Y. Braet |
| Braconidae | Metopiinae sp. 15 | Par. K. |  |  |  |  |  | 11 | Y. Braet |
| Braconidae | Metopiinae sp. 16 | Par. K. |  |  |  |  |  | 27 | Y. Braet |
| Braconidae | Metopiinae sp. 2 | Par. K. |  |  |  |  |  | 1 | Y. Braet |
| Braconidae | Metopiinae sp. 3 | Par. K. |  |  |  |  |  | 2 | Y. Braet |
| Braconidae | Metopiinae sp. 4 | Par. K. |  |  |  |  |  | 1 | Y. Braet |
| Braconidae | Metopiinae sp. 5 | Par. K. |  |  |  |  |  | 3 | Y. Braet |
| Braconidae | Metopiinae sp. 6 | Par. K. |  |  |  |  |  | 1 | Y. Braet |
| Braconidae | Metopiinae sp. 7 | Par. K. |  |  |  |  |  | 4 | Y. Braet |
| Braconidae | Metopiinae sp. 8 | Par. K. |  |  |  |  |  | 1 | Y. Braet |
| Braconidae | Metopiinae sp. 9 | Par. K. |  |  |  |  |  | 1 | Y. Braet |
| Braconidae | *Ontsira aff. imperator* | Par. I. |  |  |  |  |  | 8 | Y. Braet |
| Braconidae | *Opiinae sp. 1* | Par. K. |  |  |  |  |  | 1 | Y. Braet |
| Braconidae | *Opiinae sp. 2* | Par. K. |  |  |  |  |  | 1 | Y. Braet |
| Braconidae | *Opiinae sp. 3* | Par. K. |  |  |  |  |  | 1 | Y. Braet |
| Braconidae | *Orgilus sp. 1* | Par. K. |  |  |  |  |  | 3 | Y. Braet |
| Braconidae | *Orgilus sp. 2* | Par. K. |  |  |  |  |  | 2 | Y. Braet |
| Braconidae | *Peristenus aff. picipes* | Par. K. |  |  |  |  |  | 4 | Y. Braet |
| Braconidae | *Phanerotoma ocularis*  *Kohl, 1906* | Par. K. |  |  |  |  |  | 1 | Y. Braet |
| Braconidae | *Phanerotoma robusta*  *Zettel, 1988* | Par. K. |  |  |  |  |  | 4 | Y. Braet |
| Braconidae | *Phanerotoma sp. 1* | Par. K. |  |  |  |  |  | 1 | Y. Braet |
| Braconidae | *Phanerotoma sp. 2* | Par. K. |  |  |  |  |  | 2 | Y. Braet |
| Braconidae | *Polystenus rugosus Förster, 1862* | Par. I. |  |  |  |  |  | 3 | Y. Braet |
| Braconidae | *Rhoptrocentrus piceus*  *Marshall, 1897* | Par. I. |  |  |  |  |  | 8 | Y. Braet |
| Braconidae | Rogadinae sp. 1 | Par. K. |  |  |  |  |  | 4 | Y. Braet |
| Braconidae | Rogadinae sp. 2 | Par. K. |  |  |  |  |  | 1 | Y. Braet |
| Braconidae | Rogadinae sp. 3 | Par. K. |  |  |  |  |  | 1 | Y. Braet |
| Braconidae | Rogadinae sp. 4 | Par. K. |  |  |  |  |  | 1 | Y. Braet |
| Braconidae | Rogadinae sp. 5 | Par. K. |  |  |  |  |  | 2 | Y. Braet |
| Braconidae | *Spathius aff. brevicaudis* | Par. I. |  |  |  |  |  | 8 | Y. Braet |
| Braconidae | *Spathius aff. rubidus* | Par. I. |  |  |  |  |  | 4 | Y. Braet |
| Braconidae | ***Spathius polonicus***  ***Niezabitowski, 1910*** | Par. I. |  |  |  |  |  | 2 | Y. Braet |
| Braconidae | *Triaspis sp.1* | Par. K. |  |  |  |  |  | 1 | Y. Braet |
| Braconidae | *Triaspis sp.2* | Par. K. |  |  |  |  |  | 1 | Y. Braet |
| Braconidae | *Triaspis sp.3* | Par. K. |  |  |  |  |  | 2 | Y. Braet |
| Braconidae | *Triaspis sp.4* | Par. K. |  |  |  |  |  | 3 | Y. Braet |
| Braconidae | *Triaspis sp.5* | Par. K. |  |  |  |  |  | 4 | Y. Braet |
| Braconidae | *Triaspis sp.6* | Par. K. |  |  |  |  |  | 2 | Y. Braet |
| Braconidae | *Triaspis sp.7* | Par. K. |  |  |  |  |  | 19 | Y. Braet |
| Braconidae | *Triaspis sp.8* | Par. K. |  |  |  |  |  | 2 | Y. Braet |
| Braconidae | *Triaspis sp.9* | Par. K. |  |  |  |  |  | 1 | Y. Braet |
| Braconidae | *Wroughtonia spinator (Lepeletier de Saint Fargeau & Audinet-Serville, 1827)* | Par. K. |  |  |  |  |  | 5 | Y. Braet |
| Braconidae | *Zele albiditarsus Curtis, 1832* | Par. K. |  |  |  |  |  | 1 | Y. Braet |
| Braconidae | *Zele chlorophthalmus*  *(Spinola, 1808)* | Par. K. |  |  |  |  |  | 3 | Y. Braet |
| Braconidae | *Zele deceptor (Wesmael, 1835)* | Par. K. |  |  |  |  |  | 3 | Y. Braet |
| Cephidae | *Cephus pygmaeus*  *(Linnaeus, 1767)* | Phyto. |  |  |  |  |  | 2 | T. Noblecourt |
| Cephidae | *Janus cynosbati (Linnaeus, 1758)* | Phyto. |  |  |  |  |  | 2 | T. Noblecourt |
| Cephidae | *Janus luteipes (Le Peletier, 1823)* | Phyto. |  |  |  |  |  | 1 | T. Noblecourt |
| Ceraphronidae | *Aphanogmus sp.* | Par. K. |  |  |  |  |  | 2 | A. Staverlokk |
| Ceraphronidae | *Ceraphron sp.* | Par. K. |  |  |  |  |  | 5 | A. Staverlokk, P.N. Buhl |
| Ceraphronidae | *Conostigmus abdominalis*  *(Boheman, 1832)* | Par. K. |  |  |  |  |  | 1 | A. Staverlokk |
| Chalcididae | *Brachymeria femorata*  *(Panzer, 1801)* | Par. K. |  |  |  |  |  | 1 | E. Marhic |
| Chalcididae | *Brachymeria minuta*  *(Linnaeus, 1767)* | Par. K. |  |  |  |  |  | 1 | J-Y. Rasplus |
| Chalcididae | *Brachymeria rugulosa*  *(Förster, 1859)* | Par. K. |  |  |  |  |  | 1 | E. Marhic |
| Chalcididae | *Haltichella rufipes*  *(Olivier, 1791)* | Par. K. |  |  |  |  |  | 2 | E. Marhic, J-Y. Rasplus |
| Chrysididae | *Chrysis corusca Valkeila, 1971* | Par. I. |  |  |  |  |  | 2 | E. Marhic |
| Chrysididae | *Chrysis equestris Dahlbom, 1854* | Par. I. |  |  |  |  |  | 1 | E. Marhic, P. Rosa |
| Chrysididae | *Chrysis fasciata Olivier, 1790* | Par. I. |  |  |  |  |  | 3 | E. Marhic |
| Chrysididae | *Chrysis fulgida Linnaeus, 1761* | Par. I. |  |  |  |  |  | 6 | E. Marhic |
| Chrysididae | *Chrysis gr. ignita* | Par. I. |  |  |  |  |  | 12 | E. Marhic |
| Chrysididae | *Chrysis longula*  *Abeille de Perrin, 1879* | Par. I. |  |  |  |  |  | 2 | E. Marhic |
| Chrysididae | *Chrysis pseudobrevitarsis*  *Linsenmaier, 1951* | Par. I. |  |  |  |  |  | 4 | E. Marhic |
| Chrysididae | *Chrysis ragusae*  *De-Stefani, 1888* | Par. I. |  |  |  |  |  | 1 | E. Marhic |
| Chrysididae | *Chrysis solida  Haupt, 1956* | Par. I. |  |  |  |  |  | 1 | E. Marhic |
| Chrysididae | *Chrysis terminata*  *Dahlbom, 1854* | Par. I. |  |  |  |  |  | 34 | E. Marhic |
| Chrysididae | *Chrysura radians (Harris, 1778)* | Par. I. |  |  |  |  |  | 2 | E. Marhic |
| Chrysididae | *Hedychrum nobile*  *(Scopoli, 1763)* | Par. I. |  |  |  |  |  | 2 | E. Marhic |
| Chrysididae | *Philoctetes bidentulus (Lepeletier, 1806)* | Par. I. |  |  |  |  |  | 1 | E. Marhic, P. Rosa |
| Chrysididae | *Pseudomalus auratus*  *(Linnaeus, 1758)* | Par. I. |  |  |  |  |  | 2 | E. Marhic |
| Chrysididae | *Pseudomalus cupratus*  *(Mocsáry, 1889)* | Par. I. |  |  |  |  |  | 1 | E. Marhic |
| Chrysididae | *Pseudomalus triangulifer (Abeille de Perrin, 1877)* | Par. I. |  |  |  |  |  | 2 | E. Marhic |
| Chrysididae | *Pseudomalus violaceus*  *(Scopoli, 1763)* | Par. I. |  |  |  |  |  | 5 | E. Marhic |
| Chrysididae | *Trichrysis cyanea*  *(Linnaeus, 1758)* | Par. I. |  |  |  |  |  | 20 | E. Marhic |
| Cimbicidae | *Abia lonicerae (Linnaeus, 1758)* | N.O. Phyll. |  |  |  |  |  | 3 | T. Noblecourt |
| Colletidae | *Hylaeus sp.* | Poll./Nec. |  |  |  |  |  | 30 | LBLGC team |
| Crabronidae | *Crossocerus annulipes (Lepeletier & Brullé, 1835)* | Carn. |  |  |  |  |  | 1 | P. Burguet |
| Crabronidae | *Crossocerus cetratus*  *(Shuckard, 1837)* | Carn. |  |  |  |  |  | 3 | P. Burguet |
| Crabronidae | *Crossocerus guichardi*  *Leclercq, 1972* | Carn. |  |  |  |  |  | 1 | P. Burguet |
| Crabronidae | *Crossocerus megacephalus*  *(Rossi, 1790)* | Carn. |  |  |  |  |  | 12 | P. Burguet |
| Crabronidae | *Crossocerus nigritus*  *(Lepeletier & Brullé, 1835)* | Carn. |  |  |  |  |  | 1 | P. Burguet |
| Crabronidae | *Crossocerus ovalis*  *Lepeletier & Brullé, 1835* | Carn. |  |  |  |  |  | 1 | P. Burguet |
| Crabronidae | *Crossocerus quadrimaculatus (Fabricius, 1793)* | Carn. |  |  |  |  |  | 3 | P. Burguet |
| Crabronidae | *Crossocerus varus*  *Lepeletier & Brullé, 1835* | Carn. |  |  |  |  |  | 2 | P. Burguet |
| Crabronidae | *Ectemnius cavifrons*  *(Thomson, 1870)* | Carn. |  |  |  |  |  | 1 | P. Burguet |
| Crabronidae | *Ectemnius continuus*  *(Fabricius, 1804)* | Carn. |  |  |  |  |  | 2 | P. Burguet |
| Crabronidae | *Ectemnius lituratus*  *(Panzer, 1804)* | Carn. |  |  |  |  |  | 3 | P. Burguet |
| Crabronidae | *Ectemnius ruficornis (Zetterstedt, 1838)* | Carn. |  |  |  |  |  | 1 | P. Burguet |
| Crabronidae | *Miscophus ater*  *Lepeletier, 1845* | Carn. |  |  |  |  |  | 1 | P. Burguet |
| Crabronidae | *Nitela fallax*  *Kohl, 1884* | Carn. |  |  |  |  |  | 1 | P. Burguet |
| Crabronidae | *Nitela lucens*  *Gayubo & Felton, 2000* | Carn. |  |  |  |  |  | 12 | P. Burguet |
| Crabronidae | *Nitela spinolae*  *Latreille, 1809* | Carn. |  |  |  |  |  | 5 | P. Burguet |
| Crabronidae | *Rhopalum clavipes*  *(Linnaeus, 1758)* | Carn. |  |  |  |  |  | 2 | P. Burguet |
| Crabronidae | *Rhopalum coarctatum*  *(Scopoli, 1763)* | Carn. |  |  |  |  |  | 1 | P. Burguet |
| Crabronidae | *Tachytes panzeri*  *(Dufour, 1841)* | Carn. |  |  |  |  |  | 1 | P. Burguet |
| Crabronidae | *Trypoxylon clavicerum*  *Lepeletier de Saint Fargeau & Audinet-Serville, 1828* | Carn. |  |  |  |  |  | 7 | P. Burguet |
| Crabronidae | *Trypoxylon minus*  *Beaumont, 1945* | Carn. |  |  |  |  |  | 4 | P. Burguet |
| Cynipidae | Cynipidae sp. | Gall |  |  |  |  |  | 4199 | LBLGC team |
| Diapriidae | *Aclista sp.* | Par. K. |  |  |  |  |  | 1 | D. Notton |
| Diapriidae | *Aclista prolongata*  *(Kieffer, 1907)* | Par. K. |  |  |  |  |  | 19 | D. Notton |
| Diapriidae | *Aclista rufopetiolata*  *(Nees, 1834)* | Par. K. |  |  |  |  |  | 1 | D. Notton |
| Diapriidae | *Basalys sp.* | Par. K. |  |  |  |  |  | 1 | D. Notton |
| Diapriidae | *Belyta depressa Thomson, 1859* | Par. K. |  |  |  |  |  | 2 | D. Notton |
| Diapriidae | *Belyta sp.* | Par. K. |  |  |  |  |  | 1 | D. Notton |
| Diapriidae | *Belyta validicornis*  *Thomson, 1859* | Par. K. |  |  |  |  |  | 1 | D. Notton |
| Diapriidae | *Coptera sp. 1* | Par. K. |  |  |  |  |  | 5 | D. Notton |
| Diapriidae | *Coptera sp. 2* | Par. K. |  |  |  |  |  | 1 | D. Notton |
| Diapriidae | *Diapria cf. conica* | Par. K. |  |  |  |  |  | 5 | D. Notton |
| Diapriidae | *Ismarus sp.* | Par. K. |  |  |  |  |  | 2 | D. Notton |
| Diapriidae | *Pantoclis sp. 1* | Par. K. |  |  |  |  |  | 2 | D. Notton |
| Diapriidae | *Pantoclis sp. 2* | Par. K. |  |  |  |  |  | 1 | D. Notton |
| Diapriidae | *Pantoclis sp. 3* | Par. K. |  |  |  |  |  | 7 | D. Notton |
| Diapriidae | *Pantolyta pseudosciarivora*  *(Macek, 1998)* | Par. K. |  |  |  |  |  | 1 | D. Notton |
| Diapriidae | *Paramesius rufipes (Fonscolombe, 1832)* | Par. K. |  |  |  |  |  | 4 | D. Notton |
| Diapriidae | *Spilomicrus hemipterus Marshall, 1868* | Par. K. |  |  |  |  |  | 1 | D. Notton |
| Diapriidae | *Spilomicrus integer*  *Thomson, 1859* | Par. K. |  |  |  |  |  | 1 | D. Notton |
| Diapriidae | *Spilomicrus sp.* | Par. K. |  |  |  |  |  | 2 | D. Notton |
| Diapriidae | *Spilomicrus stigmaticalis Westwood, 1832* | Par. K. |  |  |  |  |  | 2 | D. Notton |
| Diapriidae | *Trichopria aequata*  *(Thomson, 1859)* | Par. K. |  |  |  |  |  | 13 | D. Notton |
| Diapriidae | *Trichopria cameroni*  *(Kieffer, 1909)* | Par. K. |  |  |  |  |  | 7 | D. Notton |
| Diapriidae | *Trichopria cf. morio* | Par. K. |  |  |  |  |  | 1 | D. Notton |
| Diapriidae | *Trichopria conotoma*  *(Kieffer, 1911)* | Par. K. |  |  |  |  |  | 1 | D. Notton |
| Diapriidae | *Trichopria gr. nigra* | Par. K. |  |  |  |  |  | 1 | D. Notton |
| Diapriidae | *Trichopria gr. verticillata* | Par. K. |  |  |  |  |  | 2 | D. Notton |
| Diapriidae | *Trichopria hyalinipennis (Thomson, 1859)* | Par. K. |  |  |  |  |  | 2 | D. Notton |
| Diapriidae | *Trichopria modesta*  *(Ratzeburg, 1848)* | Par. K. |  |  |  |  |  | 2 | D. Notton |
| Diapriidae | *Trichopria nigra (Nees, 1834)* | Par. K. |  |  |  |  |  | 1 | D. Notton |
| Diapriidae | *Trichopria nixoni*  *Notton, 1995* | Par. K. |  |  |  |  |  | 12 | D. Notton |
| Diapriidae | *Trichopria sociabilis*  *Masner, 1965* | Par. K. |  |  |  |  |  | 1 | D. Notton |
| Diapriidae | *Trichopria striata Notton, 1993* | Par. K. |  |  |  |  |  | 1 | D. Notton |
| Diapriidae | *Trichopria suspecta*  *(Nees, 1834)* | Par. K. |  |  |  |  |  | 1 | D. Notton |
| Diapriidae | *Trichopria verticillata*  *(Latreille, 1805)* | Par. K. |  |  |  |  |  | 2 | D. Notton |
| Diapriidae | *Zygota sp.* | Par. K. |  |  |  |  |  | 1 | D. Notton |
| Dryinidae | *Anteon brachycerum*  *(Dalman, 1823)* | Par. I. |  |  |  |  |  | 2 | E. Marhic |
| Dryinidae | *Anteon infectum (Haliday, 1837)* | Par. I. |  |  |  |  |  | 5 | E. Marhic |
| Dryinidae | *Anteon jurineanum*  *Latreille, 1809* | Par. I. |  |  |  |  |  | 6 | E. Marhic |
| Dryinidae | *Anteon scapulare*  *(Haliday, 1837)* | Par. I. |  |  |  |  |  | 1 | E. Marhic,  J. De Rond |
| Dryinidae | *Aphelopus camus Richards, 1939* | Par. I. |  |  |  |  |  | 1 | E. Marhic |
| Encyrtidae | *Anagyrus sp.* | Par. K. |  |  |  |  |  | 2 | J-Y. Rasplus |
| Encyrtidae | *Blastothrix longipennis*  *Howard, 1881* | Par. K. |  |  |  |  |  | 11 | J-Y. Rasplus |
| Encyrtidae | *Blastothrix sp.* | Par. K. |  |  |  |  |  | 2 | J-Y. Rasplus |
| Encyrtidae | *Copidosoma sp.* | Par. K. |  |  |  |  |  | 2 | J-Y. Rasplus |
| Encyrtidae | Encyrtidae sp. | Par. K. |  |  |  |  |  | 2 | J-Y. Rasplus |
| Encyrtidae | *Microterys chalcostomus (Dalman, 1820)* | Par. K. |  |  |  |  |  | 1 | J-Y. Rasplus |
| Eulophidae | *Aprostocetus sp.* | Par. K. |  |  |  |  |  | 3 | J-Y. Rasplus |
| Eulophidae | *Aulogymnus skianeuros (Ratzeburg, 1844)* | Par. K. |  |  |  |  |  | 36 | J-Y. Rasplus |
| Eulophidae | *Aulogymnus sp.* | Par. K. |  |  |  |  |  | 42 | J-Y. Rasplus |
| Eulophidae | *Aulogymnus trilineatus*  *(Mayr, 1877)* | Par. K. |  |  |  |  |  | 13 | J-Y. Rasplus |
| Eulophidae | *Chrysocharis pubicornis (Zetterstedt, 1838)* | Par. K. |  |  |  |  |  | 1 | J-Y. Rasplus |
| Eulophidae | *Chrysocharis sp.* | Par. K. |  |  |  |  |  | 1 | J-Y. Rasplus |
| Eulophidae | *Elachertus sp.* | Par. K. |  |  |  |  |  | 1 | J-Y. Rasplus |
| Eulophidae | *Entedon armigerae*  *Graham, 1971* | Par. K. |  |  |  |  |  | 1 | J-Y. Rasplus |
| Eulophidae | *Entedon zanara Walker, 1839* | Par. K. |  |  |  |  |  | 3 | J-Y. Rasplus |
| Eulophidae | *Ionympha carne (Walker, 1839)* | Par. K. |  |  |  |  |  | 1 | J-Y. Rasplus |
| Eulophidae | *Omphale lugubris Askew, 2003* | Par. K. |  |  |  |  |  | 1 | J-Y. Rasplus |
| Eulophidae | *Omphale sp.* | Par. K. |  |  |  |  |  | 1 | J-Y. Rasplus |
| Eulophidae | *Pediobius foliorum*  *(Geoffroy, 1785)* | Par. K. |  |  |  |  |  | 1 | J-Y. Rasplus |
| Eulophidae | *Pediobius lysis (Walker, 1839)* | Par. K. |  |  |  |  |  | 1 | J-Y. Rasplus |
| Eulophidae | *Pediobius sp.* | Par. K. |  |  |  |  |  | 1 | J-Y. Rasplus |
| Eulophidae | *Pnigalio sp.* | Par. K. |  |  |  |  |  | 2 | J-Y. Rasplus |
| Eupelmidae | *Anastatus bifasciatus*  *(Geoffroy, 1785)* | Par. I. |  |  |  |  |  | 1 | J-Y. Rasplus |
| Eupelmidae | *Calosota aestivalis*  *Curtis, 1836* | Par. I. |  |  |  |  |  | 4 | J-Y. Rasplus |
| Eupelmidae | *Eupelmus matranus*  *Erdös, 1947* | Par. I. |  |  |  |  |  | 1 | J-Y. Rasplus |
| Eupelmidae | *Eupelmus pini*  *Taylor, 1927* | Par. I. |  |  |  |  |  | 1 | J-Y. Rasplus |
| Eupelmidae | *Eupelmus sp.* | Par. I. |  |  |  |  |  | 1 | J-Y. Rasplus |
| Eupelmidae | *Eupelmus urozonus*  *Dalman, 1820* | Par. I. |  |  |  |  |  | 5 | J-Y. Rasplus |
| Eupelmidae | *Metapelma nobile*  *(Förster, 1860)* | Par. I. |  |  |  |  |  | 6 | J-Y. Rasplus |
| Eurytomidae | *Eurytoma sp.* | Par. I. |  |  |  |  |  | 12 | J-Y. Rasplus |
| Eurytomidae | *Sycophila sp.* | Par. K. |  |  |  |  |  | 1 | J-Y. Rasplus |
| Eurytomidae | *Sycophila submutica*  *(Thomson, 1876)* | Par. K. |  |  |  |  |  | 3 | J-Y. Rasplus |
| Evanidae | *Brachygaster minuta*  *(Olivier, 1791)* | Par. I. |  |  |  |  |  | 33 | E. Marhic |
| Figitidae | Figitidae sp. | Par. |  |  |  |  |  | 7 | LBLGC team |
| Formicidae | *Aphaenogaster subterranea*  *(Latreille, 1798)* | Carn. |  |  |  |  |  | 10 | C. Galkowski |
| Formicidae | *Camponotus fallax*  *(Nylander, 1856)* | Poly. |  |  |  |  |  | 6 | C. Galkowski |
| Formicidae | *Camponotus tergestinus*  *Mueller, 1921* | Poly. |  |  |  |  |  | 3 | C. Galkowski |
| Formicidae | *Colobopsis truncata*  *(Spinola, 1808)* | Poly. |  |  |  |  |  | 21 | C. Galkowski |
| Formicidae | *Dolichoderus quadripunctatus (Linnaeus, 1771)* | Poly. |  |  |  |  |  | 718 | C. Galkowski |
| Formicidae | *Formica cunicularia*  *Latreille, 1798* | Poly. |  |  |  |  |  | 12 | C. Galkowski |
| Formicidae | *Formica fusca Linnaeus, 1758* | Poly. |  |  |  |  |  | 18 | C. Galkowski |
| Formicidae | *Formica polyctena*  *Foerster, 1850* | Poly. |  |  |  |  |  | 6 | C. Galkowski |
| Formicidae | Formicidae sp. | Poly. |  |  |  |  |  | 155 | C. Galkowski |
| Formicidae | *Lasius alienus (Foerster, 1850)* | Poly. |  |  |  |  |  | 23 | C. Galkowski |
| Formicidae | *Lasius bicornis (Foerster, 1850)* | Poly. |  |  |  |  |  | 8 | C. Galkowski |
| Formicidae | *Lasius brunneus (Latreille, 1798)* | Poly. |  |  |  |  |  | 77 | C. Galkowski |
| Formicidae | *Lasius distinguendus*  *Emery, 1916* | Poly. |  |  |  |  |  | 5 | C. Galkowski |
| Formicidae | *Lasius emarginatus*  *(Olivier, 1792)* | Poly. |  |  |  |  |  | 3 | C. Galkowski |
| Formicidae | *Lasius flavus (Fabricius, 1782)* | Poly. |  |  |  |  |  | 13 | C. Galkowski |
| Formicidae | *Lasius fuliginosus*  *(Latreille, 1798)* | Poly. |  |  |  |  |  | 3 | C. Galkowski |
| Formicidae | *Lasius platythorax*  *Seifert, 1992* | Poly. |  |  |  |  |  | 63 | C. Galkowski |
| Formicidae | *Lasius sabularum*  *(Bondroit, 1918)* | Poly. |  |  |  |  |  | 1 | C. Galkowski |
| Formicidae | *Lasius sp.* | Poly. |  |  |  |  |  | 51 | C. Galkowski |
| Formicidae | *Lasius umbratus*  *(Nylander, 1846)* | Poly. |  |  |  |  |  | 23 | C. Galkowski |
| Formicidae | *Myrmecina graminicola (Latreille, 1802)* | Poly. |  |  |  |  |  | 39 | C. Galkowski |
| Formicidae | *Myrmica ruginodis*  *Nylander, 1846* | Poly. |  |  |  |  |  | 26 | C. Galkowski |
| Formicidae | *Myrmica sabuleti Meinert, 1861* | Poly. |  |  |  |  |  | 1 | C. Galkowski |
| Formicidae | *Myrmica scabrinodis*  *Nylander, 1846* | Carn. |  |  |  |  |  | 11 | C. Galkowski |
| Formicidae | *Myrmica specioides*  *Bondroit, 1918* | Poly. |  |  |  |  |  | 1 | C. Galkowski |
| Formicidae | *Ponera coarctata*  *(Latreille, 1802)* | Carn. |  |  |  |  |  | 1 | C. Galkowski |
| Formicidae | *Solenopsis fugax*  *(Latreille, 1798)* | Poly. |  |  |  |  |  | 1 | C. Galkowski |
| Formicidae | *Temnothorax affinis*  *(Mayr, 1855)* | Carn. |  |  |  |  |  | 9 | C. Galkowski |
| Formicidae | *Temnothorax parvulus*  *(Schenck, 1852)* | Carn. |  |  |  |  |  | 29 | C. Galkowski |
| Formicidae | *Tetramorium impurum (Foerster, 1850)* | Poly. |  |  |  |  |  | 2 | C. Galkowski |
| Halictidae | Halictidae sp. | Poll./Nec. |  |  |  |  |  | 227 | LBLGC team |
| Halictidae | *Lasioglossum bluethgeni*  *Ebmer, 1971* | Poll./Nec. |  |  |  |  | ERL (LC) | 5 | S. Flaminio |
| Halictidae | *Lasioglossum fulvicorne*  *(Kirby, 1802)* | Poll./Nec. |  |  |  |  | ERL (LC) | 3 | S. Flaminio |
| Halictidae | *Lasioglossum laticeps*  *(Schenck, 1868)* | Poll./Nec. |  |  |  |  | ERL (LC) | 1 | S. Flaminio |
| Halictidae | *Lasioglossum lativentre (Schenck, 1853)* | Poll./Nec. |  |  |  |  | ERL (LC) | 20 | S. Flaminio |
| Halictidae | *Lasioglossum leucozonium*  *(Schrank, 1781)* | Poll./Nec. |  |  |  |  | ERL (LC) | 1 | S. Flaminio |
| Halictidae | *Lasioglossum marginatum*  *(Brullé, 1832)* | Poll./Nec. |  |  |  |  | ERL (LC) | 3 | S. Flaminio |
| Halictidae | *Lasioglossum monstrificum*  *(Morawitz, 1891)* | Poll./Nec. |  |  |  |  | ERL (NT), WRL (NT) | 1 | S. Flaminio |
| Halictidae | *Lasioglossum pallens*  *(Brullé, 1832)* | Poll./Nec. |  |  |  |  | ERL (LC) | 605 | S. Flaminio |
| Halictidae | *Lasioglossum pauperatum*  *(Brullé, 1832)* | Poll./Nec. |  |  |  |  | ERL (LC) | 2 | S. Flaminio |
| Halictidae | *Lasioglossum pauxillum (Schenck, 1853)* | Poll./Nec. |  |  |  |  | ERL (LC) | 3 | S. Flaminio |
| Halictidae | *Lasioglossum punctatissimum*  *(Schenck, 1853)* | Poll./Nec. |  |  |  |  | ERL (LC) | 2 | S. Flaminio |
| Halictidae | *Lasioglossum sexnotatum*  *(Kirby, 1802)* | Poll./Nec. |  |  |  |  | ERL (NT) | 1 | S. Flaminio |
| Halictidae | *Lasioglossum subhirtum (Lepeletier, 1841)* | Poll./Nec. |  |  |  |  | ERL (LC) | 5 | S. Flaminio |
| Halictidae | *Lasioglossum transitorium*  *(Schenck, 1868)* | Poll./Nec. |  |  |  |  | ERL (LC) | 1 | S. Flaminio |
| Halictidae | *Sphecodes sp.* | Poll./Nec. |  |  |  |  |  | 44 | LBLGC team |
| Heloridae | *Helorus ruficornis*  *Förster, 1856* | Par. K. |  |  |  |  |  | 6 | E. Marhic |
| Heydeniidae | *Heydenia pretiosa*  *Förster, 1856* | Par. I. |  |  |  |  |  | 1 | J-Y. Rasplus |
| Ibaliidae | *Ibalia leucospoides (Hochenwarth, 1785)* | Par. K. |  |  |  |  |  | 1 | LBLGC team |
| Ichneumonidae | *Acrodactyla carinator*  *(Aubert, 1965)* | Par. I. |  |  |  |  |  | 1 | T. Robert |
| Ichneumonidae | *Adelognathinae sp.* | Par. I. |  |  |  |  |  | 2 | LBLGC team |
| Ichneumonidae | *Agrypon flaveolatum (Gravenhorst, 1807)* | Par. K. |  |  |  |  |  | 32 | W. Penigot |
| Ichneumonidae | *Agrypon flexorium*  *(Thunberg, 1824)* | Par. K. |  |  |  |  |  | 1 | W. Penigot |
| Ichneumonidae | *Alloplasta tomentosa (Gravenhorst, 1829)* | Par. K. |  |  |  |  |  | 44 | T. Robert |
| Ichneumonidae | *Amblyteles armatorius*  *(Forster, 1771)* | Par. K. |  |  |  |  |  | 1 | W. Penigot |
| Ichneumonidae | *Apechthis rufata*  *(Gmelin, 1790)* | Par. I. |  |  |  |  |  | 24 | T. Robert |
| Ichneumonidae | *Aphanistes gliscens*  *(Hartig, 1838)* | Par. K. |  |  |  |  |  | 17 | W. Penigot, T. Robert |
| Ichneumonidae | *Apophua bipunctoria*  *(Thunberg, 1824)* | Par. K. |  |  |  |  |  | 1 | T. Robert |
| Ichneumonidae | *Arotes albicinctus*  *Gravenhorst, 1829* | Par. K. |  |  |  |  |  | 2 | T. Robert |
| Ichneumonidae | *Astiphromma pictum*  *(Brischke, 1880)* | Par. K. |  |  |  |  |  | 164 | T. Robert |
| Ichneumonidae | ***Atractogaster semisculptus***  ***Kriechbaumer, 1872*** | Par. I. |  |  |  |  |  | 4 | T. Robert |
| Ichneumonidae | *Baeosemus mitigosus (Gravenhorst, 1829)* | Par. K. |  |  |  |  |  | 3 | W. Penigot |
| Ichneumonidae | *Barichneumon derogator (Wesmael, 1845)* | Par. K. |  |  |  |  |  | 18 | W. Penigot |
| Ichneumonidae | Campopleginae sp. 1 | Par. K. |  |  |  |  |  | 40 | LBLGC team |
| Ichneumonidae | Campopleginae sp. 10 | Par. K. |  |  |  |  |  | 2 | LBLGC team |
| Ichneumonidae | Campopleginae sp. 11 | Par. K. |  |  |  |  |  | 1 | LBLGC team |
| Ichneumonidae | Campopleginae sp. 12 | Par. K. |  |  |  |  |  | 12 | LBLGC team |
| Ichneumonidae | Campopleginae sp. 13 | Par. K. |  |  |  |  |  | 3 | LBLGC team |
| Ichneumonidae | Campopleginae sp. 14 | Par. K. |  |  |  |  |  | 1 | LBLGC team |
| Ichneumonidae | Campopleginae sp. 15 | Par. K. |  |  |  |  |  | 1 | LBLGC team |
| Ichneumonidae | Campopleginae sp. 16 | Par. K. |  |  |  |  |  | 1 | LBLGC team |
| Ichneumonidae | Campopleginae sp. 17 | Par. K. |  |  |  |  |  | 8 | LBLGC team |
| Ichneumonidae | Campopleginae sp. 18 | Par. K. |  |  |  |  |  | 18 | LBLGC team |
| Ichneumonidae | Campopleginae sp. 19 | Par. K. |  |  |  |  |  | 1 | LBLGC team |
| Ichneumonidae | Campopleginae sp. 2 | Par. K. |  |  |  |  |  | 4 | LBLGC team |
| Ichneumonidae | Campopleginae sp. 20 | Par. K. |  |  |  |  |  | 12 | LBLGC team |
| Ichneumonidae | Campopleginae sp. 21 | Par. K. |  |  |  |  |  | 3 | LBLGC team |
| Ichneumonidae | Campopleginae sp. 22 | Par. K. |  |  |  |  |  | 27 | LBLGC team |
| Ichneumonidae | Campopleginae sp. 23 | Par. K. |  |  |  |  |  | 28 | LBLGC team |
| Ichneumonidae | Campopleginae sp. 24 | Par. K. |  |  |  |  |  | 3 | LBLGC team |
| Ichneumonidae | Campopleginae sp. 25 | Par. K. |  |  |  |  |  | 3 | LBLGC team |
| Ichneumonidae | Campopleginae sp. 26 | Par. K. |  |  |  |  |  | 4 | LBLGC team |
| Ichneumonidae | Campopleginae sp. 3 | Par. K. |  |  |  |  |  | 2 | LBLGC team |
| Ichneumonidae | Campopleginae sp. 4 | Par. K. |  |  |  |  |  | 13 | LBLGC team |
| Ichneumonidae | Campopleginae sp. 5 | Par. K. |  |  |  |  |  | 6 | LBLGC team |
| Ichneumonidae | Campopleginae sp. 6 | Par. K. |  |  |  |  |  | 3 | LBLGC team |
| Ichneumonidae | Campopleginae sp. 7 | Par. K. |  |  |  |  |  | 38 | LBLGC team |
| Ichneumonidae | Campopleginae sp. 8 | Par. K. |  |  |  |  |  | 3 | LBLGC team |
| Ichneumonidae | Campopleginae sp. 9 | Par. K. |  |  |  |  |  | 2 | LBLGC team |
| Ichneumonidae | *Chasmias motatorius*  *(Fabricius, 1775)* | Par. K. |  |  |  |  |  | 1 | W. Penigot |
| Ichneumonidae | *Cidaphus alarius*  *(Gravenhorst, 1829)* | Par. K. |  |  |  |  |  | 7 | T. Robert |
| Ichneumonidae | *Coelichneumon desinatorius*  *(Thunberg, 1824)* | Par. K. |  |  |  |  |  | 3 | W. Penigot |
| Ichneumonidae | *Coelichneumon dubius (Tischbein, 1876)* | Par. K. |  |  |  |  |  | 2 | W. Penigot |
| Ichneumonidae | *Coelichneumon leucocerus*  *(Gravenhorst, 1820)* | Par. K. |  |  |  |  |  | 7 | W. Penigot |
| Ichneumonidae | *Coelichneumon probator Horstmann, 2000* | Par. K. |  |  |  |  |  | 1 | W. Penigot |
| Ichneumonidae | *Coelichneumon sinister (Wesmael, 1848)* | Par. K. |  |  |  |  |  | 1 | W. Penigot |
| Ichneumonidae | *Coleocentrus croceicornis (Gravenhorst, 1829)* | Par. K. |  |  |  |  |  | 2 | T. Robert |
| Ichneumonidae | ***Coleocentrus exareolatus***  ***Kriechbaumer, 1894*** | Par. K. |  |  |  |  |  | 1 | W. Penigot |
| Ichneumonidae | *Coleocentrus excitator*  *(Poda, 1761)* | Par. K. |  |  |  |  |  | 1 | T. Robert |
| Ichneumonidae | *Coleocentrus soleatus*  *(Gravenhorst, 1829)* | Par. K. |  |  |  |  |  | 4 | T. Robert |
| Ichneumonidae | *Colpognathus celerator (Gravenhorst, 1807)* | Par. K. |  |  |  |  |  | 1 | W. Penigot |
| Ichneumonidae | *Colpognathus divisus*  *Thomson, 1891* | Par. K. |  |  |  |  |  | 1 | W. Penigot |
| Ichneumonidae | *Cratichneumon albifrons (Stephens, 1835)* | Par. K. |  |  |  |  |  | 1 | W. Penigot |
| Ichneumonidae | *Cratichneumon coruscator*  *(Linnaeus, 1758)* | Par. K. |  |  |  |  |  | 12 | W. Penigot |
| Ichneumonidae | *Cratichneumon culex*  *(Müller, 1776)* | Par. K. |  |  |  |  |  | 30 | W. Penigot |
| Ichneumonidae | *Cratichneumon flavifrons (Schrank, 1781)* | Par. K. |  |  |  |  |  | 2 | W. Penigot |
| Ichneumonidae | *Cratichneumon rufifrons (Gravenhorst, 1829)* | Par. K. |  |  |  |  |  | 2 | W. Penigot |
| Ichneumonidae | *Cratichneumon viator*  *(Scopoli, 1763)* | Par. K. |  |  |  |  |  | 2 | W. Penigot |
| Ichneumonidae | *Crypteffigies lanius (Gravenhorst, 1829)* | Par. K. |  |  |  |  |  | 158 | W. Penigot |
| Ichneumonidae | Cryptinae sp. 1 | Par. I. |  |  |  |  |  | 208 | LBLGC team |
| Ichneumonidae | Cryptinae sp. 10 | Par. I. |  |  |  |  |  | 4 | LBLGC team |
| Ichneumonidae | Cryptinae sp. 11 | Par. I. |  |  |  |  |  | 17 | LBLGC team |
| Ichneumonidae | Cryptinae sp. 12 | Par. I. |  |  |  |  |  | 80 | LBLGC team |
| Ichneumonidae | Cryptinae sp. 13 | Par. I. |  |  |  |  |  | 9 | LBLGC team |
| Ichneumonidae | Cryptinae sp. 14 | Par. I. |  |  |  |  |  | 12 | LBLGC team |
| Ichneumonidae | Cryptinae sp. 15 | Par. I. |  |  |  |  |  | 12 | LBLGC team |
| Ichneumonidae | Cryptinae sp. 16 | Par. I. |  |  |  |  |  | 7 | LBLGC team |
| Ichneumonidae | Cryptinae sp. 17 | Par. I. |  |  |  |  |  | 2 | LBLGC team |
| Ichneumonidae | Cryptinae sp. 18 | Par. I. |  |  |  |  |  | 41 | LBLGC team |
| Ichneumonidae | Cryptinae sp. 19 | Par. I. |  |  |  |  |  | 10 | LBLGC team |
| Ichneumonidae | Cryptinae sp. 2 | Par. I. |  |  |  |  |  | 2 | LBLGC team |
| Ichneumonidae | Cryptinae sp. 20 | Par. I. |  |  |  |  |  | 2 | LBLGC team |
| Ichneumonidae | Cryptinae sp. 21 | Par. I. |  |  |  |  |  | 1 | LBLGC team |
| Ichneumonidae | Cryptinae sp. 22 | Par. I. |  |  |  |  |  | 1 | LBLGC team |
| Ichneumonidae | Cryptinae sp. 23 | Par. I. |  |  |  |  |  | 33 | LBLGC team |
| Ichneumonidae | Cryptinae sp. 24 | Par. I. |  |  |  |  |  | 5 | LBLGC team |
| Ichneumonidae | Cryptinae sp. 25 | Par. I. |  |  |  |  |  | 1 | LBLGC team |
| Ichneumonidae | Cryptinae sp. 26 | Par. I. |  |  |  |  |  | 2 | LBLGC team |
| Ichneumonidae | Cryptinae sp. 27 | Par. I. |  |  |  |  |  | 35 | LBLGC team |
| Ichneumonidae | Cryptinae sp. 28 | Par. I. |  |  |  |  |  | 3 | LBLGC team |
| Ichneumonidae | Cryptinae sp. 29 | Par. I. |  |  |  |  |  | 3 | LBLGC team |
| Ichneumonidae | Cryptinae sp. 3 | Par. I. |  |  |  |  |  | 50 | LBLGC team |
| Ichneumonidae | Cryptinae sp. 30 | Par. I. |  |  |  |  |  | 6 | LBLGC team |
| Ichneumonidae | Cryptinae sp. 31 | Par. I. |  |  |  |  |  | 1 | LBLGC team |
| Ichneumonidae | Cryptinae sp. 32 | Par. I. |  |  |  |  |  | 7 | LBLGC team |
| Ichneumonidae | Cryptinae sp. 33 | Par. I. |  |  |  |  |  | 3 | LBLGC team |
| Ichneumonidae | Cryptinae sp. 34 | Par. I. |  |  |  |  |  | 11 | LBLGC team |
| Ichneumonidae | Cryptinae sp. 35 | Par. I. |  |  |  |  |  | 1 | LBLGC team |
| Ichneumonidae | Cryptinae sp. 36 | Par. I. |  |  |  |  |  | 8 | LBLGC team |
| Ichneumonidae | Cryptinae sp. 37 | Par. I. |  |  |  |  |  | 2 | LBLGC team |
| Ichneumonidae | Cryptinae sp. 38 | Par. I. |  |  |  |  |  | 2 | LBLGC team |
| Ichneumonidae | Cryptinae sp. 39 | Par. I. |  |  |  |  |  | 6 | LBLGC team |
| Ichneumonidae | Cryptinae sp. 4 | Par. I. |  |  |  |  |  | 14 | LBLGC team |
| Ichneumonidae | Cryptinae sp. 40 | Par. I. |  |  |  |  |  | 1 | LBLGC team |
| Ichneumonidae | Cryptinae sp. 41 | Par. I. |  |  |  |  |  | 3 | LBLGC team |
| Ichneumonidae | Cryptinae sp. 42 | Par. I. |  |  |  |  |  | 46 | LBLGC team |
| Ichneumonidae | Cryptinae sp. 43 | Par. I. |  |  |  |  |  | 14 | LBLGC team |
| Ichneumonidae | Cryptinae sp. 44 | Par. I. |  |  |  |  |  | 1 | LBLGC team |
| Ichneumonidae | Cryptinae sp. 45 | Par. I. |  |  |  |  |  | 40 | LBLGC team |
| Ichneumonidae | Cryptinae sp. 46 | Par. I. |  |  |  |  |  | 1 | LBLGC team |
| Ichneumonidae | Cryptinae sp. 47 | Par. I. |  |  |  |  |  | 2 | LBLGC team |
| Ichneumonidae | Cryptinae sp. 48 | Par. I. |  |  |  |  |  | 1 | LBLGC team |
| Ichneumonidae | Cryptinae sp. 49 | Par. I. |  |  |  |  |  | 1 | LBLGC team |
| Ichneumonidae | Cryptinae sp. 5 | Par. I. |  |  |  |  |  | 13 | LBLGC team |
| Ichneumonidae | Cryptinae sp. 50 | Par. I. |  |  |  |  |  | 4 | LBLGC team |
| Ichneumonidae | Cryptinae sp. 51 | Par. I. |  |  |  |  |  | 4 | LBLGC team |
| Ichneumonidae | Cryptinae sp. 52 | Par. I. |  |  |  |  |  | 1 | LBLGC team |
| Ichneumonidae | Cryptinae sp. 53 | Par. I. |  |  |  |  |  | 1 | LBLGC team |
| Ichneumonidae | Cryptinae sp. 54 | Par. I. |  |  |  |  |  | 2 | LBLGC team |
| Ichneumonidae | Cryptinae sp. 55 | Par. I. |  |  |  |  |  | 1 | LBLGC team |
| Ichneumonidae | Cryptinae sp. 56 | Par. I. |  |  |  |  |  | 132 | LBLGC team |
| Ichneumonidae | Cryptinae sp. 57 | Par. I. |  |  |  |  |  | 3 | LBLGC team |
| Ichneumonidae | Cryptinae sp. 58 | Par. I. |  |  |  |  |  | 4 | LBLGC team |
| Ichneumonidae | Cryptinae sp. 59 | Par. I. |  |  |  |  |  | 32 | LBLGC team |
| Ichneumonidae | Cryptinae sp. 6 | Par. I. |  |  |  |  |  | 6 | LBLGC team |
| Ichneumonidae | Cryptinae sp. 60 | Par. I. |  |  |  |  |  | 44 | LBLGC team |
| Ichneumonidae | Cryptinae sp. 61 | Par. I. |  |  |  |  |  | 101 | LBLGC team |
| Ichneumonidae | Cryptinae sp. 7 | Par. I. |  |  |  |  |  | 16 | LBLGC team |
| Ichneumonidae | Cryptinae sp. 8 | Par. I. |  |  |  |  |  | 11 | LBLGC team |
| Ichneumonidae | Cryptinae sp. 9 | Par. I. |  |  |  |  |  | 6 | LBLGC team |
| Ichneumonidae | *Crytea sanguinator*  *(Rossi, 1794)* | Par. K. |  |  |  |  |  | 2 | W. Penigot |
| Ichneumonidae | Ctenopelmatinae sp. 1 | Par. K. |  |  |  |  |  | 88 | LBLGC team |
| Ichneumonidae | Ctenopelmatinae sp. 10 | Par. K. |  |  |  |  |  | 2 | LBLGC team |
| Ichneumonidae | Ctenopelmatinae sp. 11 | Par. K. |  |  |  |  |  | 1 | LBLGC team |
| Ichneumonidae | Ctenopelmatinae sp. 12 | Par. K. |  |  |  |  |  | 59 | LBLGC team |
| Ichneumonidae | Ctenopelmatinae sp. 13 | Par. K. |  |  |  |  |  | 2 | LBLGC team |
| Ichneumonidae | Ctenopelmatinae sp. 14 | Par. K. |  |  |  |  |  | 1 | LBLGC team |
| Ichneumonidae | Ctenopelmatinae sp. 15 | Par. K. |  |  |  |  |  | 2 | LBLGC team |
| Ichneumonidae | Ctenopelmatinae sp. 16 | Par. K. |  |  |  |  |  | 1 | LBLGC team |
| Ichneumonidae | Ctenopelmatinae sp. 17 | Par. K. |  |  |  |  |  | 2 | LBLGC team |
| Ichneumonidae | Ctenopelmatinae sp. 18 | Par. K. |  |  |  |  |  | 12 | LBLGC team |
| Ichneumonidae | Ctenopelmatinae sp. 19 | Par. K. |  |  |  |  |  | 1 | LBLGC team |
| Ichneumonidae | Ctenopelmatinae sp. 2 | Par. K. |  |  |  |  |  | 4 | LBLGC team |
| Ichneumonidae | Ctenopelmatinae sp. 20 | Par. K. |  |  |  |  |  | 1 | LBLGC team |
| Ichneumonidae | Ctenopelmatinae sp. 21 | Par. K. |  |  |  |  |  | 1 | LBLGC team |
| Ichneumonidae | Ctenopelmatinae sp. 22 | Par. K. |  |  |  |  |  | 33 | LBLGC team |
| Ichneumonidae | Ctenopelmatinae sp. 23 | Par. K. |  |  |  |  |  | 24 | LBLGC team |
| Ichneumonidae | Ctenopelmatinae sp. 24 | Par. K. |  |  |  |  |  | 134 | LBLGC team |
| Ichneumonidae | Ctenopelmatinae sp. 25 | Par. K. |  |  |  |  |  | 39 | LBLGC team |
| Ichneumonidae | Ctenopelmatinae sp. 26 | Par. K. |  |  |  |  |  | 73 | LBLGC team |
| Ichneumonidae | Ctenopelmatinae sp. 27 | Par. K. |  |  |  |  |  | 1 | LBLGC team |
| Ichneumonidae | Ctenopelmatinae sp. 28 | Par. K. |  |  |  |  |  | 3 | LBLGC team |
| Ichneumonidae | Ctenopelmatinae sp. 3 | Par. K. |  |  |  |  |  | 6 | LBLGC team |
| Ichneumonidae | Ctenopelmatinae sp. 4 | Par. K. |  |  |  |  |  | 6 | LBLGC team |
| Ichneumonidae | Ctenopelmatinae sp. 5 | Par. K. |  |  |  |  |  | 2 | LBLGC team |
| Ichneumonidae | Ctenopelmatinae sp. 6 | Par. K. |  |  |  |  |  | 2 | LBLGC team |
| Ichneumonidae | Ctenopelmatinae sp. 7 | Par. K. |  |  |  |  |  | 1 | LBLGC team |
| Ichneumonidae | Ctenopelmatinae sp. 8 | Par. K. |  |  |  |  |  | 8 | LBLGC team |
| Ichneumonidae | Ctenopelmatinae sp. 9 | Par. K. |  |  |  |  |  | 1 | LBLGC team |
| Ichneumonidae | *Deuteroxorides elevator*  *(Panzer, 1799)* | Par. I. |  |  |  |  |  | 15 | T. Robert |
| Ichneumonidae | *Diadromus arrisor*  *Wesmael, 1845* | Par. K. |  |  |  |  |  | 1 | W. Penigot |
| Ichneumonidae | *Diadromus troglodytes (Gravenhorst, 1829)* | Par. K. |  |  |  |  |  | 6 | W. Penigot |
| Ichneumonidae | *Dicaelotus erythrogaster (Holmgren, 1890)* | Par. K. |  |  |  |  |  | 7 | W. Penigot |
| Ichneumonidae | *Dicaelotus pictus (Schmiedeknecht, 1903)* | Par. K. |  |  |  |  |  | 12 | W. Penigot |
| Ichneumonidae | *Dicaelotus punctiventris (Thomson, 1891)* | Par. K. |  |  |  |  |  | 15 | W. Penigot |
| Ichneumonidae | *Dicaelotus resplendens Holmgren, 1890* | Par. K. |  |  |  |  |  | 1 | W. Penigot |
| Ichneumonidae | *Diphyus restitutor*  *(Wesmael, 1859)* | Par. K. |  |  |  |  |  | 1 | W. Penigot |
| Ichneumonidae | *Diplazon sp.* | Par. K. |  |  |  |  |  | 1 | LBLGC team |
| Ichneumonidae | *Dirophanes callopus*  *(Wesmael, 1845)* | Par. K. |  |  |  |  |  | 3 | W. Penigot |
| Ichneumonidae | *Dirophanes fulvitarsis (Wesmael, 1845)* | Par. K. |  |  |  |  |  | 1 | W. Penigot |
| Ichneumonidae | *Dirophanes maculicornis (Stephens, 1835)* | Par. K. |  |  |  |  |  | 64 | W. Penigot |
| Ichneumonidae | *Dolichomitus curticornis (Perkins, 1943)* | Par. I. |  |  |  |  |  | 3 | T. Robert |
| Ichneumonidae | *Dolichomitus mesocentrus (Gravenhorst, 1829)* | Par. I. |  |  |  |  |  | 1 | T. Robert |
| Ichneumonidae | ***Dolichomitus milleri***  ***Zwakhals, 2010*** | Par. I. |  |  |  |  |  | 2 | T. Robert |
| Ichneumonidae | *Dolichomitus pterelas*  *(Say, 1829)* | Par. I. |  |  |  |  |  | 4 | T. Robert |
| Ichneumonidae | *Dolichomitus sp.* | Par. I. |  |  |  |  |  | 2 | T. Robert |
| Ichneumonidae | *Dolichomitus terebrans (Ratzeburg, 1844)* | Par. I. |  |  |  |  |  | 7 | T. Robert |
| Ichneumonidae | *Drepanoctonus tibialis*  *Pfankuch, 1911* | Par. K. |  |  |  |  |  | 1 | T. Robert |
| Ichneumonidae | *Dusona sp.* | Par. K. |  |  |  |  |  | 13 | LBLGC team |
| Ichneumonidae | *Enizemum ornatum (Gravenhorst, 1829)* | Par. K. |  |  |  |  |  | 1 | T. Robert |
| Ichneumonidae | *Ephialtes manifestator (Linnaeus, 1758)* | Par. I. |  |  |  |  |  | 4 | T. Robert |
| Ichneumonidae | *Eristicus cf. clericus* | Par. K. |  |  |  |  |  | 1 | W. Penigot |
| Ichneumonidae | *Eristicus clarigator*  *(Wesmael, 1845)* | Par. K. |  |  |  |  |  | 1 | W. Penigot |
| Ichneumonidae | *Eristicus clericus*  *(Gravenhorst, 1829)* | Par. K. |  |  |  |  |  | 3 | W. Penigot |
| Ichneumonidae | *Euceros albitarsus*  *Curtis, 1837* | Par. K. |  |  |  |  |  | 9 | W. Penigot |
| Ichneumonidae | ***Euceros kiushensis***  ***Uchida*** | Par. K. |  |  |  |  |  | 4 | W. Penigot, T. Robert |
| Ichneumonidae | *Euceros pruinosus*  *(Gravenhorst, 1829)* | Par. K. |  |  |  |  |  | 1 | T. Robert |
| Ichneumonidae | *Eupalamus lacteator (Gravenhorst, 1829)* | Par. K. |  |  |  |  |  | 7 | W. Penigot |
| Ichneumonidae | *Eupalamus wesmaeli*  *(Thomson, 1886)* | Par. K. |  |  |  |  |  | 10 | W. Penigot |
| Ichneumonidae | *Fredegunda diluta*  *(Ratzeburg, 1852)* | Par. I. |  |  |  |  |  | 1 | W. Penigot |
| Ichneumonidae | *Glypta extincta*  *Ratzeburg, 1852* | Par. K. |  |  |  |  |  | 1 | T. Robert |
| Ichneumonidae | *Glypta nigrina*  *Desvignes, 1856* | Par. K. |  |  |  |  |  | 2 | T. Robert |
| Ichneumonidae | *Glypta pictipes*  *Taschenberg, 1863* | Par. K. |  |  |  |  |  | 1 | T. Robert |
| Ichneumonidae | *Glypta trochanterata*  *Bridgman, 1886* | Par. K. |  |  |  |  |  | 7 | T. Robert |
| Ichneumonidae | *Herpestomus brunnicornis (Gravenhorst, 1829)* | Par. K. |  |  |  |  |  | 1 | W. Penigot |
| Ichneumonidae | *Homotherus locutor*  *(Thunberg, 1824)* | Par. K. |  |  |  |  |  | 1 | W. Penigot |
| Ichneumonidae | *Homotherus varipes (Gravenhorst, 1829)* | Par. K. |  |  |  |  |  | 1 | W. Penigot |
| Ichneumonidae | *Hoplismenus terrificus*  *Wesmael, 1848* | Par. K. |  |  |  |  |  | 1 | W. Penigot |
| Ichneumonidae | *Hyperacmus crassicornis (Gravenhorst, 1829)* | Par. K. |  |  |  |  |  | 1 | W. Penigot, T. Robert |
| Ichneumonidae | *Ichneumon bucculentus Wesmael, 1845* | Par. K. |  |  |  |  |  | 2 | W. Penigot |
| Ichneumonidae | *Ichneumon cf. exilicornis* | Par. K. |  |  |  |  |  | 2 | W. Penigot |
| Ichneumonidae | *Ichneumon gracilicornis Gravenhorst, 1829* | Par. K. |  |  |  |  |  | 1 | W. Penigot |
| Ichneumonidae | *Ichneumon inquinatus*  *Wesmael, 1845* | Par. K. |  |  |  |  |  | 7 | W. Penigot |
| Ichneumonidae | *Ichneumon molitorius*  *Linnaeus, 1760* | Par. K. |  |  |  |  |  | 1 | W. Penigot |
| Ichneumonidae | *Ichneumon simulans*  *Tischbein, 1873* | Par. K. |  |  |  |  |  | 8 | W. Penigot |
| Ichneumonidae | *Ichneumon stenocerus*  *Thomson, 1887* | Par. K. |  |  |  |  |  | 1 | W. Penigot |
| Ichneumonidae | *Ichneumon suspiciosus Wesmael, 1845* | Par. K. |  |  |  |  |  | 3 | W. Penigot |
| Ichneumonidae | *Ischnoceros caligatus (Gravenhorst, 1829)* | Par. I. |  |  |  |  |  | 4 | W. Penigot, T. Robert |
| Ichneumonidae | *Ischnoceros rusticus*  *(Geoffroy, 1785)* | Par. I. |  |  |  |  |  | 13 | W. Penigot, T. Robert |
| Ichneumonidae | *Itoplectis alternans (Gravenhorst, 1829)* | Par. I. |  |  |  |  |  | 7 | W. Penigot, T. Robert |
| Ichneumonidae | *Itoplectis clavicornis*  *(Thomson, 1889)* | Par. I. |  |  |  |  |  | 1 | T. Robert |
| Ichneumonidae | *Itoplectis maculator*  *(Fabricius, 1775)* | Par. I. |  |  |  |  |  | 19 | T. Robert |
| Ichneumonidae | *Liotryphon caudatus*  *(Ratzeburg, 1848)* | Par. I. |  |  |  |  |  | 3 | T. Robert |
| Ichneumonidae | *Liotryphon punctulatus (Ratzeburg, 1848)* | Par. I. |  |  |  |  |  | 1 | T. Robert |
| Ichneumonidae | *Lissonota biguttata*  *Holmgren, 1860* | Par. K. |  |  |  |  |  | 19 | T. Robert |
| Ichneumonidae | *Lissonota complicator*  *Aubert, 1967* | Par. K. |  |  |  |  |  | 1 | T. Robert |
| Ichneumonidae | *Lissonota cruentator*  *(Panzer, 1809)* | Par. K. |  |  |  |  |  | 1 | T. Robert |
| Ichneumonidae | *Lissonota deversor*  *Gravenhorst, 1829* | Par. K. |  |  |  |  |  | 2 | T. Robert |
| Ichneumonidae | *Lissonota folii*  *Thomson, 1877* | Par. K. |  |  |  |  |  | 35 | T. Robert |
| Ichneumonidae | *Lissonota luffiator*  *Aubert, 1969* | Par. K. |  |  |  |  |  | 29 | T. Robert |
| Ichneumonidae | *Lissonota mutator*  *Aubert, 1969* | Par. K. |  |  |  |  |  | 1 | T. Robert |
| Ichneumonidae | *Lissonota palpalis*  *Thomson, 1889* | Par. K. |  |  |  |  |  | 1 | T. Robert |
| Ichneumonidae | *Lissonota variabilis*  *Holmgren, 1860* | Par. K. |  |  |  |  |  | 1 | T. Robert |
| Ichneumonidae | *Lissonota versicolor*  *Holmgren, 1860* | Par. K. |  |  |  |  |  | 1 | T. Robert |
| Ichneumonidae | *Lycorina triangulifera*  *Holmgren, 1859* | Par. K. |  |  |  |  |  | 1 | T. Robert |
| Ichneumonidae | *Lymantrichneumon disparis (Poda, 1761)* | Par. K. |  |  |  |  |  | 19 | W. Penigot |
| Ichneumonidae | Mesochorinae sp.1 | Par. K. |  |  |  |  |  | 32 | LBLGC team |
| Ichneumonidae | Mesochorinae sp.2 | Par. K. |  |  |  |  |  | 2 | LBLGC team |
| Ichneumonidae | Mesochorinae sp.3 | Par. K. |  |  |  |  |  | 3 | LBLGC team |
| Ichneumonidae | Mesochorinae sp.4 | Par. K. |  |  |  |  |  | 1 | LBLGC team |
| Ichneumonidae | Mesochorinae sp.5 | Par. K. |  |  |  |  |  | 3 | LBLGC team |
| Ichneumonidae | Mesochorinae sp.6 | Par. K. |  |  |  |  |  | 2 | LBLGC team |
| Ichneumonidae | Metopiinae sp. 1 | Par. K. |  |  |  |  |  | 90 | LBLGC team |
| Ichneumonidae | Metopiinae sp. 2 | Par. K. |  |  |  |  |  | 7 | LBLGC team |
| Ichneumonidae | Metopiinae sp. 3 | Par. K. |  |  |  |  |  | 8 | LBLGC team |
| Ichneumonidae | Metopiinae sp. 4 | Par. K. |  |  |  |  |  | 1 | LBLGC team |
| Ichneumonidae | Metopiinae sp. 5 | Par. K. |  |  |  |  |  | 6 | LBLGC team |
| Ichneumonidae | Metopiinae sp. 6 | Par. K. |  |  |  |  |  | 1 | LBLGC team |
| Ichneumonidae | *Misetus oculatus*  *Wesmael, 1845* | Par. K. |  |  |  |  |  | 1 | W. Penigot |
| Ichneumonidae | *Netelia cristata*  *(Thomson, 1888)* | Par. K. |  |  |  |  |  | 3 | T. Robert |
| Ichneumonidae | *Netelia latungula*  *(Thomson, 1888)* | Par. K. |  |  |  |  |  | 2 | T. Robert |
| Ichneumonidae | *Netelia testacea*  *(Gravenhorst, 1829)* | Par. K. |  |  |  |  |  | 1 | T. Robert |
| Ichneumonidae | *Odontocolon quercinum (Thomson, 1877)* | Par. I. |  |  |  |  |  | 2 | W. Penigot, T. Robert |
| Ichneumonidae | ***Ophion brocki***  ***Johansson, 2019*** | Par. K. |  |  |  |  |  | 2 | W. Penigot |
| Ichneumonidae | ***Ophion confusus***  ***Johansson, 2019*** | Par. K. |  |  |  |  |  | 1 | W. Penigot |
| Ichneumonidae | *Ophion gr. mocsaryi* | Par. K. |  |  |  |  |  | 4 | W. Penigot |
| Ichneumonidae | *Ophion minutus*  *Kriechbaumer, 1879* | Par. K. |  |  |  |  |  | 83 | W. Penigot, T. Robert |
| Ichneumonidae | *Ophion ocellaris*  *Ulbricht, 1926* | Par. K. |  |  |  |  |  | 2 | W. Penigot |
| Ichneumonidae | ***Ophion splendens***  ***Johansson, 2019*** | Par. K. |  |  |  |  |  | 1 | W. Penigot |
| Ichneumonidae | *Ophion variegatus*  *Rudow, 1883* | Par. K. |  |  |  |  |  | 1 | W. Penigot |
| Ichneumonidae | *Ophion ventricosus*  *Gravenhorst, 1829* | Par. K. |  |  |  |  |  | 3 | W. Penigot |
| Ichneumonidae | *Orgichneumon calcatorius*  *(Thunberg, 1822)* | Par. K. |  |  |  |  |  | 1 | W. Penigot |
| Ichneumonidae | Orthocentrinae sp. 1 | Par. K. |  |  |  |  |  | 49 | LBLGC team |
| Ichneumonidae | Orthocentrinae sp. 2 | Par. K. |  |  |  |  |  | 4 | LBLGC team |
| Ichneumonidae | Orthocentrinae sp. 3 | Par. K. |  |  |  |  |  | 7 | LBLGC team |
| Ichneumonidae | Orthocentrinae sp. 4 | Par. K. |  |  |  |  |  | 2 | LBLGC team |
| Ichneumonidae | Orthocentrinae sp. 5 | Par. K. |  |  |  |  |  | 1 | LBLGC team |
| Ichneumonidae | Orthocentrinae sp. 6 | Par. K. |  |  |  |  |  | 1 | LBLGC team |
| Ichneumonidae | Orthocentrinae sp. 7 | Par. K. |  |  |  |  |  | 1 | LBLGC team |
| Ichneumonidae | *Oxytorus armatus*  *Thomson, 1883* | Par. K. |  |  |  |  |  | 1 | W. Penigot |
| Ichneumonidae | *Paracoelichneumon rubens (Boyer de Fonscolombe, 1847)* | Par. K. |  |  |  |  |  | 2 | W. Penigot |
| Ichneumonidae | *Phaeogenes cf. semivulpinus* | Par. K. |  |  |  |  |  | 2 | W. Penigot |
| Ichneumonidae | *Phaeogenes planifrons Wesmael, 1845* | Par. K. |  |  |  |  |  | 1 | W. Penigot |
| Ichneumonidae | *Phaeogenes semivulpinus (Gravenhorst, 1829)* | Par. K. |  |  |  |  |  | 38 | W. Penigot |
| Ichneumonidae | *Phaeogenes spiniger*  *(Gravenhorst, 1829)* | Par. K. |  |  |  |  |  | 1 | E. Diller |
| Ichneumonidae | *Phytodietus cf. albipes* | Par. K. |  |  |  |  |  | 1 | W. Penigot |
| Ichneumonidae | *Pimpla contemplator*  *(Müller, 1776)* | Par. I. |  |  |  |  |  | 21 | T. Robert |
| Ichneumonidae | *Pimpla insignatoria (Gravenhorst, 1807)* | Par. I. |  |  |  |  |  | 1 | T. Robert |
| Ichneumonidae | *Pimpla rufipes*  *(Miller, 1759)* | Par. I. |  |  |  |  |  | 3 | T. Robert |
| Ichneumonidae | *Pimpla turionellae*  *(Linnaeus, 1758)* | Par. I. |  |  |  |  |  | 24 | T. Robert |
| Ichneumonidae | Pimplinae sp. | Par. I. |  |  |  |  |  | 1 | LBLGC team |
| Ichneumonidae | *Podoschistus scutellaris (Desvignes, 1856)* | Par. I. |  |  |  |  |  | 30 | T. Robert |
| Ichneumonidae | Poemeniinae sp. | Par. I. |  |  |  |  |  | 1 | LBLGC team |
| Ichneumonidae | *Polysphincta boops*  *Tschek, 1869* | Par. I. |  |  |  |  |  | 4 | Y. Braet, T. Robert |
| Ichneumonidae | *Pristomerus vulnerator*  *(Panzer, 1799)* | Par. K. |  |  |  |  |  | 1 | T. Robert |
| Ichneumonidae | *Pseudorhyssa alpestris (Holmgren, 1860)* | Par. I. |  |  |  |  |  | 1 | T. Robert |
| Ichneumonidae | *Rhyssella approximator (Fabricius, 1793)* | Par. I. |  |  |  |  |  | 1 | T. Robert |
| Ichneumonidae | *Rhyssella obliterata (Gravenhorst, 1829)* | Par. I. |  |  |  |  |  | 27 | W. Penigot, T. Robert |
| Ichneumonidae | *Rynchobanchus flavopictus*  *Heinrich, 1937* | Par. K. |  |  |  |  |  | 1 | T. Robert |
| Ichneumonidae | *Scambus calobatus (Gravenhorst, 1829)* | Par. I. |  |  |  |  |  | 24 | T. Robert |
| Ichneumonidae | *Scambus nigricans*  *(Thomson, 1877)* | Par. I. |  |  |  |  |  | 1 | T. Robert |
| Ichneumonidae | *Scambus planatus*  *(Hartig, 1838)* | Par. I. |  |  |  |  |  | 95 | T. Robert |
| Ichneumonidae | *Schizopyga flavifrons*  *Holmgren, 1856* | Par. I. |  |  |  |  |  | 1 | T. Robert |
| Ichneumonidae | *Stenichneumon militarius (Thunberg, 1824)* | Par. K. |  |  |  |  |  | 1 | W. Penigot |
| Ichneumonidae | ***Stenobarichneumon protervus  (Holmgren, 1864)*** | Par. K. |  |  |  |  |  | 1 | W. Penigot |
| Ichneumonidae | *Stilbopinae sp.* | Par. K. |  |  |  |  |  | 1 | LBLGC team |
| Ichneumonidae | *Stilbops vetulus*  *(Gravenhorst, 1829)* | Par. K. |  |  |  |  |  | 444 | T. Robert |
| Ichneumonidae | *Syrphophilus tricinctorius (Thunberg, 1824)* | Par. K. |  |  |  |  |  | 1 | T. Robert |
| Ichneumonidae | *Syspasis albiguttata (Gravenhorst, 1820)* | Par. K. |  |  |  |  |  | 3 | W. Penigot |
| Ichneumonidae | *Syspasis rufina*  *(Gravenhorst, 1820)* | Par. K. |  |  |  |  |  | 3 | W. Penigot |
| Ichneumonidae | ***Syspasis tauma***  ***(Heinrich, 1951)*** | Par. K. |  |  |  |  |  | 1 | W. Penigot |
| Ichneumonidae | Tersilochinae sp.1 | Par. K. |  |  |  |  |  | 35 | LBLGC team |
| Ichneumonidae | Tersilochinae sp.2 | Par. K. |  |  |  |  |  | 1 | LBLGC team |
| Ichneumonidae | Tersilochinae sp.3 | Par. K. |  |  |  |  |  | 1 | LBLGC team |
| Ichneumonidae | *Theronia atalantae*  *(Poda, 1761)* | Par. I. |  |  |  |  |  | 2 | T. Robert |
| Ichneumonidae | *Theronia laevigata*  *(Tschek, 1869)* | Par. I. |  |  |  |  |  | 1 | T. Robert |
| Ichneumonidae | *Tromatobia lineatoria*  *(Villers, 1789)* | Par. I. |  |  |  |  |  | 3 | T. Robert |
| Ichneumonidae | *Tromatobia ovivora*  *(Boheman, 1821)* | Par. I. |  |  |  |  |  | 1 | T. Robert |
| Ichneumonidae | Tryphoninae sp. 1 | Par. K. |  |  |  |  |  | 16 | LBLGC team |
| Ichneumonidae | Tryphoninae sp. 2 | Par. K. |  |  |  |  |  | 22 | LBLGC team |
| Ichneumonidae | Tryphoninae sp. 3 | Par. K. |  |  |  |  |  | 1 | LBLGC team |
| Ichneumonidae | Tryphoninae sp. 4 | Par. K. |  |  |  |  |  | 16 | LBLGC team |
| Ichneumonidae | Tryphoninae sp. 5 | Par. K. |  |  |  |  |  | 1 | LBLGC team |
| Ichneumonidae | Tryphoninae sp. 6 | Par. K. |  |  |  |  |  | 21 | LBLGC team |
| Ichneumonidae | Tryphoninae sp. 7 | Par. K. |  |  |  |  |  | 1 | LBLGC team |
| Ichneumonidae | Tryphoninae sp. 8 | Par. K. |  |  |  |  |  | 1 | LBLGC team |
| Ichneumonidae | *Tycherus cephalotes*  *(Wesmael, 1845)* | Par. K. |  |  |  |  |  | 3 | W. Penigot |
| Ichneumonidae | *Tycherus flavidens*  *(Wesmael, 1845)* | Par. K. |  |  |  |  |  | 5 | W. Penigot |
| Ichneumonidae | *Tycherus infimus*  *(Wesmael, 1845)* | Par. K. |  |  |  |  |  | 1 | W. Penigot |
| Ichneumonidae | *Tycherus ischiomelinus (Gravenhorst, 1829)* | Par. K. |  |  |  |  |  | 7 | W. Penigot |
| Ichneumonidae | *Tycherus stipator*  *(Wesmael, 1855)* | Par. K. |  |  |  |  |  | 1 | W. Penigot |
| Ichneumonidae | *Tycherus suspicax*  *(Wesmael, 1845)* | Par. K. |  |  |  |  |  | 9 | W. Penigot |
| Ichneumonidae | *Ulesta perspicua*  *(Wesmael, 1857)* | Par. K. |  |  |  |  |  | 1 | W. Penigot |
| Ichneumonidae | *Virgichneumon dumeticola*  *(Gravenhorst, 1829)* | Par. K. |  |  |  |  |  | 3 | W. Penigot |
| Ichneumonidae | *Virgichneumon tergenus (Gravenhorst, 1820)* | Par. K. |  |  |  |  |  | 8 | W. Penigot |
| Ichneumonidae | *Vulgichneumon bimaculatus*  *(Schrank, 1776)* | Par. K. |  |  |  |  |  | 2 | W. Penigot |
| Ichneumonidae | *Vulgichneumon deceptor (Scopoli, 1763)* | Par. K. |  |  |  |  |  | 1 | W. Penigot |
| Ichneumonidae | *Vulgichneumon suavis (Gravenhorst, 1820)* | Par. K. |  |  |  |  |  | 1 | W. Penigot |
| Ichneumonidae | *Woldstedtius citropectoralis (Schmiedeknecht, 1926)* | Par. K. |  |  |  |  |  | 1 | T. Robert |
| Ichneumonidae | *Xorides berlandi*  *Clément, 1938* | Par. I. |  |  |  |  |  | 4 | W. Penigot, T. Robert |
| Ichneumonidae | ***Xorides corcyrensis***  ***(Kriechbaumer, 1894)*** | Par. I. |  |  |  |  |  | 1 | T. Robert |
| Ichneumonidae | *Xorides csikii*  *Clément, 1938* | Par. I. |  |  |  |  |  | 137 | W. Penigot, T. Robert |
| Ichneumonidae | *Xorides filiformis*  *(Gravenhorst, 1829)* | Par. I. |  |  |  |  |  | 6 | W. Penigot, T. Robert |
| Ichneumonidae | *Xorides fuligator*  *(Thunberg, 1824)* | Par. I. |  |  |  |  |  | 4 | W. Penigot |
| Ichneumonidae | *Xorides gravenhorstii*  *(Curtis, 1831)* | Par. I. |  |  |  |  |  | 5 | W. Penigot, T. Robert |
| Ichneumonidae | *Xorides indicatorius*  *(Latreille, 1806)* | Par. I. |  |  |  |  |  | 1 | T. Robert |
| Ichneumonidae | *Xorides minutus*  *Clément, 1938* | Par. I. |  |  |  |  |  | 14 | W. Penigot |
| Ichneumonidae | *Xorides praecatorius*  *(Fabricius, 1793)* | Par. I. |  |  |  |  |  | 30 | W. Penigot, T. Robert |
| Ichneumonidae | *Xorides rufipes*  *(Gravenhorst, 1829)* | Par. I. |  |  |  |  |  | 12 | W. Penigot, T. Robert |
| Ichneumonidae | *Xorides sepulchralis*  *(Holmgren, 1860)* | Par. I. |  |  |  |  |  | 1 | T. Robert |
| Megachilidae | *Megachile sp.* | Poll./Nec. |  |  |  |  |  | 3 | LBLGC team |
| Megaspilidae | *Conostigmus sp.* | Par. I. |  |  |  |  |  | 14 | A. Staverlokk |
| Megaspilidae | *Dendrocerus sp.* | Par. I. |  |  |  |  |  | 9 | A. Staverlokk |
| Megaspilidae | *Platyceraphron muscidarum*  *Kieffer, 1906* | Par. I. |  |  |  |  |  | 1 | A. Staverlokk |
| Megaspilidae | *Trichosteresis glabra*  *(Boheman, 1832)* | Par. I. |  |  |  |  |  | 1 | A. Staverlokk |
| Megastigmidae | *Megastigmus dorsalis*  *(Fabricius, 1798)* | Par. I. |  |  |  |  |  | 1 | J-Y. Rasplus |
| Mellinidae | *Mellinus arvensis*  *(Linnaeus, 1758)* | Carn. |  |  |  |  |  | 1 | P. Burguet |
| Mutillidae | *Myrmosa atra*  *Panzer, 1801* | Par. I. |  |  |  |  |  | 4 | F. Herbrecht, E. Marhic |
| Ooderidae | *Oodera formosa*  *(Giraud, 1863)* | Par. I. |  |  |  |  |  | 1 | J-Y. Rasplus |
| Ormyridae | *Ormyrus pomaceus*  *(Geoffroy, 1785)* | Par. I. |  |  |  |  |  | 1 | J-Y. Rasplus |
| Ormyridae | *Ormyrus wachtli*  *Mayr, 1904* | Par. I. |  |  |  |  |  | 5 | J-Y. Rasplus |
| Orussidae | *Orussus unicolor*  *Latreille, 1812* | Par. I. |  |  |  |  |  | 17 | T. Noblecourt |
| Pamphiliidae | *Acantholyda posticalis pinivora Enslin, 1918* | N.O. Phyll. |  |  |  |  |  | 1 | T. Noblecourt |
| Pamphiliidae | *Pamphilius balteatus*  *(Fallén, 1808)* | N.O. Phyll. |  |  |  |  |  | 1 | T. Noblecourt |
| Pamphiliidae | *Pamphilius hortorum*  *(Klug, 1808)* | N.O. Phyll. |  |  |  |  |  | 3 | T. Noblecourt |
| Pamphiliidae | *Pamphilius ignymontiensis*  *Lacourt, 1973* | N.O. Phyll. |  |  |  |  |  | 1 | T. Noblecourt |
| Pamphiliidae | *Pamphilius marginatus (Audinet-Serville, 1823)* | N.O. Phyll. |  |  |  |  |  | 18 | T. Noblecourt |
| Pamphiliidae | *Pamphilius pallipes*  *(Zetterstedt, 1838)* | N.O. Phyll. |  |  |  |  |  | 6 | T. Noblecourt |
| Pamphiliidae | *Pamphilius sylvarum*  *(Stephens, 1835)* | O. Phyll. |  |  |  |  |  | 1 | T. Noblecourt |
| Pamphiliidae | *Pamphilius sylvaticus*  *(Linnaeus, 1758)* | N.O. Phyll. |  |  |  |  |  | 5 | T. Noblecourt |
| Pamphiliidae | *Pamphilius varius*  *(Audinet-Serville, 1823)* | N.O. Phyll. |  |  |  |  |  | 2 | T. Noblecourt |
| Pemphredonidae | *Passaloecus corniger*  *Shuckard, 1837* | Carn. |  |  |  |  |  | 3 | P. Burguet |
| Pemphredonidae | *Passaloecus gracilis*  *Curtis, 1834* | Carn. |  |  |  |  |  | 1 | P. Burguet |
| Pemphredonidae | *Passaloecus insignis*  *(Vander Linden, 1829)* | Carn. |  |  |  |  |  | 24 | P. Burguet, F. Herbrecht |
| Pemphredonidae | *Passaloecus vandeli*  *Ribaut, 1952* | Carn. |  |  |  |  |  | 6 | P. Burguet |
| Pemphredonidae | *Pemphredon austriaca*  *(Kohl, 1888)* | Carn. |  |  |  |  |  | 2 | P. Burguet |
| Pemphredonidae | *Pemphredon inornata*  *Say, 1824* | Carn. |  |  |  |  |  | 3 | P. Burguet |
| Pemphredonidae | *Pemphredon lethifer*  *(Shuckard, 1837)* | Carn. |  |  |  |  |  | 54 | P. Burguet |
| Pemphredonidae | *Pemphredon lugens*  *Dahlbom, 1842* | Carn. |  |  |  |  |  | 7 | P. Burguet |
| Pemphredonidae | *Pemphredon lugubris*  *(Fabricius, 1793)* | Carn. |  |  |  |  |  | 56 | P. Burguet |
| Pemphredonidae | *Spilomena troglodytes*  *(Vander Linden, 1829)* | Carn. |  |  |  |  |  | 3 | P. Burguet |
| Pemphredonidae | *Stigmus pendulus*  *Panzer, 1804* | Carn. |  |  |  |  |  | 1 | P. Burguet |
| Pemphredonidae | *Stigmus solskyi*  *A. Morawitz, 1864* | Carn. |  |  |  |  |  | 1 | P. Burguet |
| Perilampidae | *Perilampus ruficornis*  *(Fabricius, 1793)* | Par. I. |  |  |  |  |  | 2614 | J-Y. Rasplus |
| Platygastridae | *Inostemma curtum*  *Szelényi, 1938* | Par. K. |  |  |  |  |  | 7 | P.N. Buhl |
| Platygastridae | *Leptacis laodice*  *(Walker, 1836)* | Par. K. |  |  |  |  |  | 10 | P.N. Buhl |
| Platygastridae | *Leptacis nydia*  *(Walker, 1836)* | Par. K. |  |  |  |  |  | 1 | P.N. Buhl |
| Platygastridae | *Metaclisis sp.* | Par. K. |  |  |  |  |  | 1 | P.N. Buhl |
| Platygastridae | *Platygaster betulae*  *(Kieffer, 1916)* | Par. K. |  |  |  |  |  | 3 | P.N. Buhl |
| Platygastridae | *Platygaster betularia*  *Kieffer, 1916* | Par. K. |  |  |  |  |  | 1 | P.N. Buhl |
| Platygastridae | *Platygaster marchali*  *Kieffer, 1906* | Par. K. |  |  |  |  |  | 1 | P.N. Buhl |
| Platygastridae | *Platygaster munita*  *Walker, 1836* | Par. K. |  |  |  |  |  | 1 | P.N. Buhl |
| Platygastridae | *Platygaster sp. 1* | Par. K. |  |  |  |  |  | 1 | P.N. Buhl |
| Platygastridae | *Platygaster sp. 2* | Par. K. |  |  |  |  |  | 1 | P.N. Buhl |
| Platygastridae | *Platygaster sp. 3* | Par. K. |  |  |  |  |  | 1 | P.N. Buhl |
| Platygastridae | *Platygaster striatithorax*  *Buhl, 1994* | Par. K. |  |  |  |  |  | 548 | P.N. Buhl |
| Platygastridae | Platygastridae sp. | Par. K. |  |  |  |  |  | 3 | J-Y. Rasplus |
| Platygastridae | *Synopeas larides (Walker, 1836)* | Par. K. |  |  |  |  |  | 6 | P.N. Buhl |
| Platygastridae | *Synopeas lugubre*  *Thomson, 1859* | Par. K. |  |  |  |  |  | 20 | P.N. Buhl |
| Platygastridae | *Synopeas myles  (Walker, 1836)* | Par. K. |  |  |  |  |  | 3 | P.N. Buhl |
| Platygastridae | *Synopeas sosis (Walker, 1836)* | Par. K. |  |  |  |  |  | 7 | P.N. Buhl |
| Platygastridae | *Synopeas sp. 1* | Par. K. |  |  |  |  |  | 2 | P.N. Buhl |
| Platygastridae | *Synopeas sp. 2* | Par. K. |  |  |  |  |  | 1 | P.N. Buhl |
| Pompilidae | *Agenioideus cinctellus*  *(Spinola, 1808)* | Par. I. |  |  |  |  |  | 8 | F. Herbrecht |
| Pompilidae | *Anoplius nigerrimus*  *(Scopoli, 1763)* | Par. I. |  |  |  |  |  | 1 | F. Herbrecht |
| Pompilidae | *Anoplius sp.* | Par. I. |  |  |  |  |  | 1 | F. Herbrecht |
| Pompilidae | *Anoplius viaticus*  *(Linnaeus, 1758)* | Par. I. |  |  |  |  |  | 1 | F. Herbrecht |
| Pompilidae | *Aporus unicolor Spinola, 1808* | Par. I. |  |  |  |  |  | 82 | F. Herbrecht |
| Pompilidae | *Arachnospila anceps*  *(Wesmael, 1851)* | Par. I. |  |  |  |  |  | 1 | F. Herbrecht |
| Pompilidae | *Arachnospila spissa*  *(Schioedte, 1837)* | Par. I. |  |  |  |  |  | 22 | F. Herbrecht |
| Pompilidae | *Arachnospila trivialis*  *(Dahlbom, 1843)* | Par. I. |  |  |  |  |  | 2 | F. Herbrecht |
| Pompilidae | *Auplopus carbonarius*  *(Scopoli, 1763)* | Par. I. |  |  |  |  |  | 128 | F. Herbrecht |
| Pompilidae | *Caliadurgus fasciatellus (Spinola, 1808)* | Par. I. |  |  |  |  |  | 13 | F. Herbrecht |
| Pompilidae | *Cryptocheilus notatus*  *(Rossius, 1792)* | Par. I. |  |  |  |  |  | 10 | F. Herbrecht |
| Pompilidae | *Cryptocheilus versicolor*  *(Scopoli, 1763)* | Par. I. |  |  |  |  |  | 2 | F. Herbrecht |
| Pompilidae | *Deuteragenia bifasciata*  *(Geoffroy, 1785)* | Par. I. |  |  |  |  |  | 25 | F. Herbrecht |
| Pompilidae | *Deuteragenia monticola*  *(Wahis, 1972)* | Par. I. |  |  |  |  |  | 77 | F. Herbrecht |
| Pompilidae | *Deuteragenia subintermedia*  *(Magretti, 1886)* | Par. I. |  |  |  |  |  | 134 | F. Herbrecht |
| Pompilidae | *Deuteragenia variegata*  *(Linnaeus, 1758)* | Par. I. |  |  |  |  |  | 4 | F. Herbrecht |
| Pompilidae | *Deuteragenia vechti*  *(Day, 1979)* | Par. I. |  |  |  |  |  | 44 | F. Herbrecht |
| Pompilidae | *Evagetes siculus*  *(Lepeletier, 1845)* | Par. I. |  |  |  |  |  | 2 | F. Herbrecht |
| Pompilidae | *Poecilagenia rubricans (Lepeletier, 1845)* | Par. I. |  |  |  |  |  | 1 | F. Herbrecht |
| Pompilidae | *Priocnemis agilis*  *(Shuckard, 1837)* | Par. I. |  |  |  |  |  | 1 | F. Herbrecht |
| Pompilidae | *Priocnemis cordivalvata*  *Haupt, 1927* | Par. I. |  |  |  |  |  | 1 | F. Herbrecht |
| Pompilidae | *Priocnemis coriacea*  *Dahlbom, 1843* | Par. I. |  |  |  |  |  | 6 | F. Herbrecht |
| Pompilidae | *Priocnemis enslini*  *Haupt, 1927* | Par. I. |  |  |  |  |  | 1 | F. Herbrecht |
| Pompilidae | *Priocnemis exaltata*  *(Fabricius, 1775)* | Par. I. |  |  |  |  |  | 1 | F. Herbrecht |
| Pompilidae | *Priocnemis fallax*  *Verhoeff, 1922* | Par. I. |  |  |  |  |  | 1 | F. Herbrecht |
| Pompilidae | *Priocnemis fennica*  *Haupt, 1927* | Par. I. |  |  |  |  |  | 1 | F. Herbrecht |
| Pompilidae | *Priocnemis hyalinata*  *(Fabricius, 1793)* | Par. I. |  |  |  |  |  | 60 | F. Herbrecht |
| Pompilidae | *Priocnemis parvula*  *Dahlbom, 1845* | Par. I. |  |  |  |  |  | 1 | F. Herbrecht |
| Pompilidae | *Priocnemis perturbator*  *(Harris, 1780)* | Par. I. |  |  |  |  |  | 38 | F. Herbrecht |
| Pompilidae | *Priocnemis sp.* | Par. I. |  |  |  |  |  | 4 | F. Herbrecht |
| Pompilidae | *Priocnemis schioedtei*  *Haupt, 1927* | Par. I. |  |  |  |  |  | 20 | F. Herbrecht |
| Pompilidae | *Priocnemis susterai*  *Haupt, 1927* | Par. I. |  |  |  |  |  | 7 | F. Herbrecht |
| Pompilidae | *Priocnemis vulgaris*  *(Dufour, 1841)* | Par. I. |  |  |  |  |  | 8 | F. Herbrecht |
| Proctotrupidae | *Exallonyx longicornis*  *(Nees, 1834)* | Par. K. |  |  |  |  |  | 4 | E. Marhic |
| Psenidae | *Pseneo exaratus*  *(Eversmann, 1849)* | Carn. |  |  |  |  |  | 1 | P. Burguet |
| Psenidae | *Psenulus fuscipennis*  *(Dahlbom, 1843)* | Carn. |  |  |  |  |  | 2 | P. Burguet |
| Psenidae | *Psenulus pallipes*  *(Panzer, 1798)* | Carn. |  |  |  |  |  | 13 | P. Burguet, F. Herbrecht |
| Pteromalidae | *Arthrolytus ocellus*  *(Walker, 1834)* | Par. K. |  |  |  |  |  | 1 | J-Y. Rasplus |
| Pteromalidae | *Callitula bicolor*  *Spinola, 1811* | Par. K. |  |  |  |  |  | 1 | J-Y. Rasplus |
| Pteromalidae | *Cheiropachus quadrum (Fabricius, 1787)* | Par. K. |  |  |  |  |  | 1 | J-Y. Rasplus |
| Pteromalidae | *Cleonymus laticornis*  *Walker, 1837* | Par. K. |  |  |  |  |  | 5 | J-Y. Rasplus |
| Pteromalidae | *Coelopisthia pachycera*  *Masi, 1924* | Par. K. |  |  |  |  |  | 4 | J-Y. Rasplus |
| Pteromalidae | *Conomorium amplum*  *(Walker, 1835)* | Par. K. |  |  |  |  |  | 6 | J-Y. Rasplus |
| Pteromalidae | *Cyclogastrella simplex*  *(Walker, 1834)* | Par. K. |  |  |  |  |  | 2 | J-Y. Rasplus |
| Pteromalidae | *Cyrtogaster vulgaris*  *Walker, 1833* | Par. K. |  |  |  |  |  | 13 | J-Y. Rasplus |
| Pteromalidae | *Dibrachys microgastri*  *(Bouché, 1834)* | Par. K. |  |  |  |  |  | 1 | J-Y. Rasplus |
| Pteromalidae | *Gastrancistrus compressus*  *Walker, 1834* | Par. I. |  |  |  |  |  | 1 | J-Y. Rasplus |
| Pteromalidae | ***Gastrancistrus laticeps***  ***Graham, 1969*** | Par. I. |  |  |  |  |  | 3 | J-Y. Rasplus |
| Pteromalidae | *Gastrancistrus puncticollis*  *(Thomson, 1876)* | Par. I. |  |  |  |  |  | 1 | J-Y. Rasplus |
| Pteromalidae | *Gastrancistrus sp. 1* | Par. I. |  |  |  |  |  | 9 | J-Y. Rasplus |
| Pteromalidae | *Gastrancistrus sp. 2* | Par. I. |  |  |  |  |  | 1 | J-Y. Rasplus |
| Pteromalidae | *Meraporus graminicola*  *Walker, 1834* | Par. I. |  |  |  |  |  | 24 | J-Y. Rasplus |
| Pteromalidae | *Mesopolobus diffinis*  *(Walker, 1834)* | Par. I. |  |  |  |  |  | 23 | J-Y. Rasplus |
| Pteromalidae | *Mesopolobus dubius*  *(Walker, 1834)* | Par. I. |  |  |  |  |  | 1 | J-Y. Rasplus |
| Pteromalidae | *Mesopolobus fuscipes*  *(Walker, 1834)* | Par. I. |  |  |  |  |  | 4 | J-Y. Rasplus |
| Pteromalidae | *Mesopolobus sp. 1* | Par. I. |  |  |  |  |  | 17 | J-Y. Rasplus |
| Pteromalidae | *Mesopolobus sp. 2* | Par. I. |  |  |  |  |  | 3 | J-Y. Rasplus |
| Pteromalidae | *Mesopolobus sp. 3* | Par. I. |  |  |  |  |  | 1 | J-Y. Rasplus |
| Pteromalidae | *Mesopolobus sp. 4* | Par. I. |  |  |  |  |  | 1 | J-Y. Rasplus |
| Pteromalidae | *Mesopolobus tibialis (Westwood, 1833)* | Par. I. |  |  |  |  |  | 42 | J-Y. Rasplus |
| Pteromalidae | *Mesopolobus xanthocerus*  *(Thomson, 1878)* | Par. I. |  |  |  |  |  | 1 | J-Y. Rasplus |
| Pteromalidae | *Miscogastrinae sp.* | Par. K. |  |  |  |  |  | 1 | J-Y. Rasplus |
| Pteromalidae | *Ormocerus latus Walker, 1834* | Par. |  |  |  |  |  | 12 | J-Y. Rasplus |
| Pteromalidae | *Ormocerus vernalis*  *Walker, 1834* | Par. |  |  |  |  |  | 105 | J-Y. Rasplus |
| Pteromalidae | *Pachyneuron aphidis*  *(Bouché, 1834)* | Par. I. |  |  |  |  |  | 1 | J-Y. Rasplus |
| Pteromalidae | *Pachyneuron sp.* | Par. I. |  |  |  |  |  | 1 | J-Y. Rasplus |
| Pteromalidae | Pteromalidae sp. | Par. |  |  |  |  |  | 11 | J-Y. Rasplus |
| Pteromalidae | *Pteromalus sp. 1* | Par. K. |  |  |  |  |  | 10 | J-Y. Rasplus |
| Pteromalidae | *Pteromalus sp. 2* | Par. K. |  |  |  |  |  | 4 | J-Y. Rasplus |
| Pteromalidae | *Pteromalus sp. 3* | Par. K. |  |  |  |  |  | 1 | J-Y. Rasplus |
| Pteromalidae | *Sceptrothelys parviclava*  *Graham, 1969* | Par. I. |  |  |  |  |  | 6 | J-Y. Rasplus |
| Pteromalidae | *Sphegigaster brevicornis*  *(Walker, 1833)* | Par. I. |  |  |  |  |  | 2 | J-Y. Rasplus |
| Pteromalidae | *Trigonoderus cyanescens (Förster, 1841)* | Par. I. |  |  |  |  |  | 3 | J-Y. Rasplus |
| Pteromalidae | *Trigonoderus princeps Westwood, 1832* | Par. I. |  |  |  |  |  | 7 | J-Y. Rasplus |
| Scelionidae | *Idris sp.* | Par. I. |  |  |  |  |  | 1 | P.N. Buhl |
| Scelionidae | *Scelio sp.* | Par. I. |  |  |  |  |  | 7 | P.N. Buhl |
| Scelionidae | *Telenomus chloropus*  *(Thomson, 1861)* | Par. I. |  |  |  |  |  | 8 | P.N. Buhl |
| Scelionidae | *Telenomus sp.* | Par. I. |  |  |  |  |  | 1 | P.N. Buhl |
| Scelionidae | *Trimorus flavipes*  *(Walker, 1836)* | Par. I. |  |  |  |  |  | 1 | P.N. Buhl |
| Scelionidae | *Trimorus puncticollis*  *(Thomson, 1859)* | Par. I. |  |  |  |  |  | 1 | P.N. Buhl |
| Scelionidae | *Trimorus sp.* | Par. I. |  |  |  |  |  | 1 | P.N. Buhl |
| Scelionidae | *Trissolcus cultratus*  *(Mayr, 1879)* | Par. I. |  |  |  |  |  | 5 | P.N. Buhl |
| Scelionidae | *Triteleia peyerimhoffi*  *(Kieffer, 1908)* | Par. I. |  |  |  |  |  | 5 | P.N. Buhl |
| Sphecidae | *Ammophila sabulosa*  *(Linnaeus, 1758)* | Carn. |  |  |  |  |  | 1 | F. Herbrecht |
| Sphecidae | *Isodontia mexicana*  *(Saussure, 1867)* | Carn. |  |  |  |  |  | 1 | P. Burguet |
| Stephanidae | *Stephanus serrator*  *(Fabricius, 1798)* | Par. I. |  |  |  |  |  | 2 | LBLGC team |
| Tenthredinidae | *Aglaostigma aucupariae*  *(Klug, 1817)* | N.O. Phyll. |  |  |  |  |  | 6 | T. Noblecourt |
| Tenthredinidae | *Allantus togatus*  *(Panzer, 1801)* | O. Phyll. |  |  |  |  |  | 1 | T. Noblecourt |
| Tenthredinidae | *Allantus viennensis*  *(Schrank, 1781)* | N.O. Phyll. |  |  |  |  |  | 1 | T. Noblecourt |
| Tenthredinidae | *Aneugmenus padi*  *(Linnaeus, 1760)* | N.O. Phyll. |  |  |  |  |  | 13 | T. Noblecourt |
| Tenthredinidae | *Athalia cordata*  *Audinet-Serville, 1823* | N.O. Phyll. |  |  |  |  |  | 2 | T. Noblecourt |
| Tenthredinidae | *Athalia rosae*  *(Linnaeus, 1758)* | N.O. Phyll. |  |  |  |  |  | 4 | T. Noblecourt |
| Tenthredinidae | *Caliroa cinxia (Klug, 1816)* | O. Phyll. |  |  |  |  |  | 8 | T. Noblecourt |
| Tenthredinidae | *Claremontia alternipes*  *(Klug, 1816)* | N.O. Phyll. |  |  |  |  |  | 2 | T. Noblecourt |
| Tenthredinidae | *Claremontia uncta*  *(Klug, 1816)* | N.O. Phyll. |  |  |  |  |  | 1 | T. Noblecourt |
| Tenthredinidae | *Cytisogaster chambersi*  *(Benson, 1947)* | N.O. Phyll. |  |  |  |  |  | 2 | T. Noblecourt |
| Tenthredinidae | *Cytisogaster genistae*  *(Benson, 1947)* | N.O. Phyll. |  |  |  |  |  | 16 | T. Noblecourt |
| Tenthredinidae | *Cytisogaster picta*  *(Klug, 1817)* | N.O. Phyll. |  |  |  |  |  | 8 | T. Noblecourt |
| Tenthredinidae | *Dineura stilata*  *(Klug, 1816)* | N.O. Phyll. |  |  |  |  |  | 1 | T. Noblecourt |
| Tenthredinidae | *Dolerus aeneus*  *Hartig, 1837* | N.O. Phyll. |  |  |  |  |  | 6 | T. Noblecourt |
| Tenthredinidae | *Dolerus gonager*  *(Fabricius, 1781)* | N.O. Phyll. |  |  |  |  |  | 11 | T. Noblecourt |
| Tenthredinidae | *Dolerus madidus*  *(Klug, 1818)* | N.O. Phyll. |  |  |  |  |  | 3 | T. Noblecourt |
| Tenthredinidae | *Dolerus niger*  *(Linnaeus, 1767)* | N.O. Phyll. |  |  |  |  |  | 1 | T. Noblecourt |
| Tenthredinidae | *Dolerus nigratus*  *(O.F. Müller, 1776)* | N.O. Phyll. |  |  |  |  |  | 1 | T. Noblecourt |
| Tenthredinidae | *Dolerus sanguinicollis*  *(Klug, 1818)* | N.O. Phyll. |  |  |  |  |  | 8 | T. Noblecourt |
| Tenthredinidae | *Emphytus cingulatus*  *(Scopoli, 1763)* | N.O. Phyll. |  |  |  |  |  | 1 | T. Noblecourt |
| Tenthredinidae | *Empria candidata*  *(Fallén, 1808)* | N.O. Phyll. |  |  |  |  |  | 9 | T. Noblecourt |
| Tenthredinidae | *Empria excisa*  *(C.G. Thomson, 1871)* | N.O. Phyll. |  |  |  |  |  | 1 | T. Noblecourt |
| Tenthredinidae | *Empria granatensis*  *Lacourt, 1988* | N.O. Phyll. |  |  |  |  |  | 2 | T. Noblecourt |
| Tenthredinidae | *Empria immersa*  *(Klug, 1818)* | N.O. Phyll. |  |  |  |  |  | 2 | T. Noblecourt |
| Tenthredinidae | *Empria liturata*  *(Gmelin, 1790)* | N.O. Phyll. |  |  |  |  |  | 6 | T. Noblecourt |
| Tenthredinidae | *Empria tridens*  *(Konow, 1896)* | N.O. Phyll. |  |  |  |  |  | 30 | T. Noblecourt |
| Tenthredinidae | *Eutomostethus luteiventris (Klug, 1816)* | N.O. Phyll. |  |  |  |  |  | 11 | T. Noblecourt |
| Tenthredinidae | *Euura fagi*  *(Zaddach, 1876)* | N.O. Phyll. |  |  |  |  |  | 4 | T. Noblecourt |
| Tenthredinidae | *Euura krausi*  *(Taeger & Blank, 1998)* | N.O. Phyll. |  |  |  |  |  | 2 | T. Noblecourt |
| Tenthredinidae | *Euura lateralis*  *(Konow, 1895)* | N.O. Phyll. |  |  |  |  |  | 2 | T. Noblecourt |
| Tenthredinidae | *Euura myosotidis*  *(Fabricius, 1804)* | N.O. Phyll. |  |  |  |  |  | 1 | T. Noblecourt |
| Tenthredinidae | *Euura papillosa*  *(Retzius, 1783)* | N.O. Phyll. |  |  |  |  |  | 3 | T. Noblecourt |
| Tenthredinidae | *Euura poecilonota*  *(Zaddach, 1876)* | N.O. Phyll. |  |  |  |  |  | 4 | T. Noblecourt |
| Tenthredinidae | *Halidamia affinis*  *(Fallén, 1807)* | N.O. Phyll. |  |  |  |  |  | 1 | T. Noblecourt |
| Tenthredinidae | *Harpiphorus lepidus*  *(Klug, 1818)* | O. Phyll. |  |  |  |  |  | 59 | T. Noblecourt |
| Tenthredinidae | *Hoplocampa cantoti*  *Chevin, 1986* | N.O. Phyll. |  |  |  |  |  | 23 | T. Noblecourt |
| Tenthredinidae | *Hoplocampa chrysorrhoea*  *(Klug, 1816)* | N.O. Phyll. |  |  |  |  |  | 12 | T. Noblecourt |
| Tenthredinidae | *Hoplocampa testudinea*  *(Klug, 1816)* | N.O. Phyll. |  |  |  |  |  | 2 | T. Noblecourt |
| Tenthredinidae | *Macrophya albicincta*  *(Schrank, 1776)* | N.O. Phyll. |  |  |  |  |  | 11 | T. Noblecourt |
| Tenthredinidae | *Macrophya alboannulata*  *A. Costa, 1859* | N.O. Phyll. |  |  |  |  |  | 7 | T. Noblecourt |
| Tenthredinidae | *Macrophya annulata*  *(Geoffroy, 1785)* | N.O. Phyll. |  |  |  |  |  | 1 | T. Noblecourt |
| Tenthredinidae | *Macrophya blanda*  *(Fabricius, 1775)* | N.O. Phyll. |  |  |  |  |  | 17 | T. Noblecourt |
| Tenthredinidae | *Macrophya duodecimpunctata*  *(Linnaeus, 1758)* | N.O. Phyll. |  |  |  |  |  | 4 | T. Noblecourt |
| Tenthredinidae | *Macrophya militaris (Klug, 1817)* | N.O. Phyll. |  |  |  |  |  | 16 | T. Noblecourt |
| Tenthredinidae | *Macrophya montana*  *(Scopoli, 1763)* | N.O. Phyll. |  |  |  |  |  | 1 | T. Noblecourt |
| Tenthredinidae | *Macrophya sanguinolenta (Gmelin, 1790)* | N.O. Phyll. |  |  |  |  |  | 1 | T. Noblecourt |
| Tenthredinidae | *Mesoneura opaca*  *(Fabricius, 1775)* | O. Phyll. |  |  |  |  |  | 60 | T. Noblecourt |
| Tenthredinidae | *Monophadnoides ruficruris (Brullé, 1832)* | N.O. Phyll. |  |  |  |  |  | 1 | T. Noblecourt |
| Tenthredinidae | *Monophadnus pallescens (Gmelin, 1790)* | N.O. Phyll. |  |  |  |  |  | 6 | T. Noblecourt |
| Tenthredinidae | *Monsoma pulveratum*  *(Retzius, 1783)* | N.O. Phyll. |  |  |  |  |  | 1 | T. Noblecourt |
| Tenthredinidae | *Nematus alniastri (Scharfenberg, 1805)* | N.O. Phyll. |  |  |  |  |  | 1 | T. Noblecourt |
| Tenthredinidae | *Nematus brischkei*  *Zaddach, 1876* | N.O. Phyll. |  |  |  |  |  | 1 | T. Noblecourt |
| Tenthredinidae | *Nematus umbratus*  *C.G. Thomson, 1871* | N.O. Phyll. |  |  |  |  |  | 10 | T. Noblecourt |
| Tenthredinidae | *Pachyprotasis antennata*  *(Klug, 1817)* | N.O. Phyll. |  |  |  |  |  | 1 | T. Noblecourt |
| Tenthredinidae | *Pachyprotasis rapae*  *(Linnaeus, 1767)* | N.O. Phyll. |  |  |  |  |  | 40 | T. Noblecourt |
| Tenthredinidae | *Pachyprotasis simulans*  *(Klug, 1817)* | N.O. Phyll. |  |  |  |  |  | 6 | T. Noblecourt |
| Tenthredinidae | *Periclista albida*  *(Klug, 1816)* | O. Phyll. |  |  |  |  |  | 113 | T. Noblecourt |
| Tenthredinidae | *Periclista albipennis*  *(Zaddach, 1859)* | O. Phyll. |  |  |  |  |  | 21 | T. Noblecourt |
| Tenthredinidae | *Periclista lineolata*  *(Klug, 1816)* | O. Phyll. |  |  |  |  |  | 7 | T. Noblecourt |
| Tenthredinidae | *Periclista pilosa*  *Chevin, 1971* | O. Phyll. |  |  |  |  |  | 30 | T. Noblecourt |
| Tenthredinidae | *Periclista pubescens*  *(Zaddach, 1859)* | O. Phyll. |  |  |  |  |  | 1 | T. Noblecourt |
| Tenthredinidae | *Pristiphora armata*  *(C.G. Thomson, 1863)* | N.O. Phyll. |  |  |  |  |  | 7 | T. Noblecourt |
| Tenthredinidae | *Pristiphora fausta*  *(Hartig, 1837)* | O. Phyll. |  |  |  |  |  | 23 | T. Noblecourt |
| Tenthredinidae | *Pristiphora insularis*  *Rohwer, 1910* | N.O. Phyll. |  |  |  |  |  | 2 | T. Noblecourt |
| Tenthredinidae | *Pristiphora leucopus*  *(Hellén, 1948)* | N.O. Phyll. |  |  |  |  |  | 1 | T. Noblecourt |
| Tenthredinidae | *Pristiphora maesta*  *(Zaddach, 1876)* | N.O. Phyll. |  |  |  |  |  | 10 | T. Noblecourt |
| Tenthredinidae | *Pristiphora monogyniae*  *(Hartig, 1840)* | N.O. Phyll. |  |  |  |  |  | 2 | T. Noblecourt |
| Tenthredinidae | *Pristiphora pallidiventris*  *(Fallén, 1808)* | N.O. Phyll. |  |  |  |  |  | 2 | T. Noblecourt |
| Tenthredinidae | *Pristiphora punctifrons (Thomson, 1871)* | N.O. Phyll. |  |  |  |  |  | 1 | T. Noblecourt |
| Tenthredinidae | *Pristiphora subbifida*  *(C.G. Thomson, 1871)* | N.O. Phyll. |  |  |  |  |  | 5 | T. Noblecourt |
| Tenthredinidae | *Pristiphora tetrica*  *(Zaddach, 1883)* | N.O. Phyll. |  |  |  |  |  | 1 | T. Noblecourt |
| Tenthredinidae | *Profenusa pygmaea*  *(Klug, 1816)* | O. Phyll. |  |  |  |  |  | 5 | T. Noblecourt |
| Tenthredinidae | *Rhogogaster chlorosoma (Benson, 1943)* | N.O. Phyll. |  |  |  |  |  | 1 | T. Noblecourt |
| Tenthredinidae | *Rhogogaster scalaris Klug, 1817* | O. Phyll. |  |  |  |  |  | 14 | T. Noblecourt |
| Tenthredinidae | *Scolioneura vicina*  *Konow, 1894* | N.O. Phyll. |  |  |  |  |  | 1 | T. Noblecourt |
| Tenthredinidae | *Strongylogaster multifasciata*  *(Geoffroy, 1785)* | N.O. Phyll. |  |  |  |  |  | 52 | T. Noblecourt |
| Tenthredinidae | *Strongylogaster xanthocera*  *(Stephens, 1835)* | N.O. Phyll. |  |  |  |  |  | 19 | T. Noblecourt |
| Tenthredinidae | *Tenthredo atra*  *Linnaeus, 1758* | N.O. Phyll. |  |  |  |  |  | 48 | T. Noblecourt |
| Tenthredinidae | *Tenthredo ferruginea*  *Schrank, 1776* | N.O. Phyll. |  |  |  |  |  | 44 | T. Noblecourt |
| Tenthredinidae | *Tenthredo livida*  *Linnaeus, 1758* | N.O. Phyll. |  |  |  |  |  | 2 | T. Noblecourt |
| Tenthredinidae | *Tenthredo solitaria Scopoli, 1763* | N.O. Phyll. |  |  |  |  |  | 1 | T. Noblecourt |
| Tenthredinidae | *Tenthredo temula*  *Scopoli, 1763* | N.O. Phyll. |  |  |  |  |  | 232 | T. Noblecourt |
| Tenthredinidae | *Tenthredo zona*  *Klug, 1817* | N.O. Phyll. |  |  |  |  |  | 2 | T. Noblecourt |
| Tenthredinidae | *Tenthredopsis coquebertii*  *(Klug, 1817)* | N.O. Phyll. |  |  |  |  |  | 1 | T. Noblecourt |
| Tenthredinidae | *Tenthredopsis litterata (Geoffroy, 1785)* | N.O. Phyll. |  |  |  |  |  | 29 | T. Noblecourt |
| Tenthredinidae | *Tenthredopsis nassata (Linnaeus, 1767)* | N.O. Phyll. |  |  |  |  |  | 16 | T. Noblecourt |
| Tenthredinidae | *Tenthredopsis scutellaris (Fabricius, 1804)* | N.O. Phyll. |  |  |  |  |  | 27 | T. Noblecourt |
| Tenthredinidae | *Tomostethus nigritus*  *(Fabricius, 1804)* | N.O. Phyll. |  |  |  |  |  | 1 | T. Noblecourt |
| Tiphiidae | *Tiphia femorata Fabricius, 1775* | Par. |  |  |  |  |  | 92 | F. Herbrecht, E. Marhic |
| Torymidae | *Diomorus calcaratus*  *(Nees, 1834)* | Par. I. |  |  |  |  |  | 2 | J-Y. Rasplus |
| Torymidae | *Monodontomerus aereus Walker, 1834* | Par. I. |  |  |  |  |  | 3 | J-Y. Rasplus |
| Torymidae | *Podagrion pachymerum (Walker, 1833)* | Par. I. |  |  |  |  |  | 1 | J-Y. Rasplus |
| Torymidae | *Torymus aucupariae*  *(Rodzianko, 1908)* | Par. I. |  |  |  |  |  | 3 | J-Y. Rasplus |
| Torymidae | *Torymus auratus*  *(O.F. Müller, 1764)* | Par. I. |  |  |  |  |  | 5 | J-Y. Rasplus |
| Torymidae | *Torymus chlorocopes*  *Boheman, 1834* | Par. I. |  |  |  |  |  | 3 | J-Y. Rasplus |
| Torymidae | ***Torymus druparum***  ***Boheman, 1834*** | Par. I. |  |  |  |  |  | 2 | J-Y. Rasplus |
| Torymidae | *Torymus flavipes*  *(Walker, 1833)* | Par. I. |  |  |  |  |  | 1 | J-Y. Rasplus |
| Torymidae | *Torymus sp. 1* | Par. I. |  |  |  |  |  | 11 | J-Y. Rasplus |
| Torymidae | *Torymus sp. 2* | Par. I. |  |  |  |  |  | 4 | J-Y. Rasplus |
| Vespidae | *Allodynerus koenigi*  *(Dusmet, 1917)* | Carn. |  |  |  |  |  | 2 | B. Gereys |
| Vespidae | *Allodynerus rossii*  *(Lepeletier, 1841)* | Carn. |  |  |  |  |  | 5 | B. Gereys |
| Vespidae | *Ancistrocerus antilope*  *(Panzer, 1798)* | Carn. |  |  |  |  |  | 3 | B. Gereys |
| Vespidae | *Ancistrocerus auctus*  *(Fabricius, 1793)* | Carn. |  |  |  |  |  | 1 | B. Gereys |
| Vespidae | *Ancistrocerus dusmetiolus (Strand, 1914)* | Carn. |  |  |  |  |  | 4 | B. Gereys |
| Vespidae | *Ancistrocerus longispinosus longispinosus (Saussure, 1855)* | Carn. |  |  |  |  |  | 1 | B. Gereys |
| Vespidae | *Ancistrocerus nigricornis*  *(Curtis, 1826)* | Carn. |  |  |  |  |  | 15 | B. Gereys |
| Vespidae | *Ancistrocerus oviventris oviventris (Wesmael, 1836)* | Carn. |  |  |  |  |  | 1 | B. Gereys |
| Vespidae | *Ancistrocerus parietinus (Linnaeus, 1760)* | Carn. |  |  |  |  |  | 8 | B. Gereys |
| Vespidae | *Ancistrocerus trifasciatus subsp. trifasciatus (Müller, 1776)* | Carn. |  |  |  |  |  | 9 | B. Gereys |
| Vespidae | *Discoelius dufourii subsp. dufourii Lepeletier, 1841* | Carn. |  |  |  |  |  | 1 | B. Gereys |
| Vespidae | *Discoelius zonalis (Panzer, 1801)* | Carn. |  |  |  |  |  | 4 | B. Gereys |
| Vespidae | *Dolichovespula media (Retzius, 1783)* | Carn. |  |  |  |  |  | 6 | LBLGC team |
| Vespidae | *Euodynerus (Pareuodynerus) quadrifasciatus subsp. quadrifasciatus  (Fabricius, 1793)* | Carn. |  |  |  |  |  | 1 | B. Gereys |
| Vespidae | *Polistes nimpha*  *(Christ, 1791)* | Carn. |  |  |  |  |  | 7 | B. Gereys |
| Vespidae | *Stenodynerus chevrieranus*  *(Saussure, 1855)* | Carn. |  |  |  |  |  | 2 | B. Gereys |
| Vespidae | *Symmorphus bifasciatus (Linnaeus, 1760)* | Carn. |  |  |  |  |  | 2 | B. Gereys |
| Vespidae | *Symmorphus murarius (Linnaeus, 1758)* | Carn. |  |  |  |  |  | 3 | B. Gereys |
| Vespidae | *Vespa crabro*  *Linnaeus, 1758* | Carn. |  |  |  |  |  | 240 | LBLGC team |
| Vespidae | *Vespula germanica (Fabricius, 1793)* | Carn. |  |  |  |  |  | 7 | LBLGC team |
| Vespidae | *Vespula vulgaris (Linnaeus, 1758)* | Carn. |  |  |  |  |  | 647 | LBLGC team |
| Xiphydriidae | *Xiphydria longicollis (Geoffroy, 1785)* | Xyl. |  |  |  |  |  | 149 | T. Noblecourt |
| Xyelidae | *Xyela curva Benson, 1938* | Poll./Nec. |  |  |  |  |  | 8 | T. Noblecourt |
| Xyelidae | *Xyela julii*  *(Brébisson, 1818)* | Poll./Nec. |  |  |  |  |  | 11 | T. Noblecourt |

Table S3. Species diversity estimators (Chao, Jackknife 1 and 2, bootstrap) for the Hymenoptera sampled with green multi-funnel traps and flight interception traps, for the overall sampling in three forests (21 stands and 42 plots), and according to the sanitary condition of plots and stands where the insects have been sampled. Unidentified Cynipidae, Figitidae, Halictidae, Andrenidae, Pteromalidae, Encyrtidae and Platygastridae have been removed. Nb.: number of plots used to calculate the estimators.

| **Scale** | **Health status** | **Observed number of taxa** | **Nb.** | **Chao** | **Jackknife 1** | **Jackknife 2** | **Bootstrap** | **Range** |
| --- | --- | --- | --- | --- | --- | --- | --- | --- |
| All |  | 918 | 42 | 1,407 ± 70 | 1,279 ± 65 | 1,505 | 1,074 ± 32 | 1,074 - 1,505 |
| Plot | Healthy | 548 | 16 | 898 +/- 59 | 800 +/- 76 | 956 | 657 +/- 37 | 657 – 956 |
|  | Moderately declining | 583 | 14 | 990 +/- 69 | 839 +/- 77 | 1,007 | 694 +/- 36 | 694 - 1,007 |
|  | Severely declining | 555 | 12 | 911 +/- 58 | 812 +/- 90 | 968 | 667 +/- 45 | 667 – 968 |
| Stand | Healthy | 501 | 14 | 943 ± 79 | 742 ± 76 | 909 | 604 ± 37 | 604 – 943 |
|  | Moderately declining | 629 | 16 | 943 ± 51 | 891 ± 76 | 1,041 | 745 ± 39 | 745 - 1,041 |
|  | Severely  declining | 532 | 12 | 898 ± 61 | 783 ± 78 | 940 | 641 ± 35 | 641 - 940 |

Table S4. Pairwise PERMANOVA based on the communities of canopy-dwelling Hymenoptera sampled in healthy (H: < 30% of trees were in decline), moderately declining (MD: 30-60% of trees were in decline), and severely declining (SD: > 60% of trees were in decline) plots and stands. Scale indicate at which spatial scale (plot or stand) the decline severity has been estimated.

| **Scale** | **Group 1** | **Group 2** | **F value** | ***P*** | **R²** |
| --- | --- | --- | --- | --- | --- |
| Plot | SD | MD | 1.81 | 0.03 | 0.07 |
|  | SD | H | 4.37 | < 0.001 | 0.14 |
|  | MD | H | 1.32 | 0.16 | 0.05 |
| Stand | SD | H | 5.83 | < 0.001 | 0.20 |
|  | SD | MD | 3.88 | < 0.001 | 0.13 |
|  | H | MD | 1.82 | 0.02 | 0.06 |

Table S5. Overall α diversity, β turnover and β nestedness between pairs of plot decline category. Overall β diversity corresponds to Sorensen dissimilarity, β turnover to Simpson dissimilarity and β nestedness to the difference between Sorensen and Simpson dissimilarity.

|  | **Overall** *α* **diversity** | *β* **turnover** | *β* **nestedness** |
| --- | --- | --- | --- |
| Among plots | 0.95 | 0.93 (97.89%) | 0.02 (2.11%) |
| Among categories of plot decline | 0.45 | 0.44 (97.78%) | 0.01 (2.22%) |
| Between H and MD plots | 0.37 | 0.35 (94.59%) | 0.02 (5.41%) |
| Between MD and SD plots | 0.38 | 0.37 (97.37%) | 0.01 (2.63%) |
| Between H and SD plots | 0.378 | 0.374 (98.94%) | 0.004 (1.06%) |

Table S6. Indicator species for decline categories estimated at the plot scale (i.e. on 10 trees, with H: healthy plots (< 30% of trees were in decline), MD: moderately declining plots (30-60% of trees were in decline), and SD: severely declining plots (> 60% of trees were in decline)), as estimated with the multipatt function (indicspecies R-package) and the IndVal index (IndVal.g) with 1,000 permutations. A corresponds to the probability that a site belongs to a particular category of decline because the species has been sampled at that site, and B is the probability of sampling the species if the site corresponds to the target decline category. Species in bold are indicators at both plot and stand scales (see Table S7).

| **Plot decline category** | **Indicator species** | **Family** | **Larval trophic guild** | **A** | **B** | **IndVal index** |
| --- | --- | --- | --- | --- | --- | --- |
| H | ***Coleocentrus soleatus*** | Ichneumonidae | parasitoid | 1.00 | 0.25 | 0.5* |
| MD | *Hylaeus sp.* | Colletidae | pollinivorous/nectarivorous | 0.67 | 0.71 | 0.69 * |
| MD | *Andrena fulva* | Andrenidae | pollinivorous/nectarivorous | 0.89 | 0.36 | 0.56 ** |
| MD | *Pseudomalus violaceus* | Chrysididae | parasitoid | 1.00 | 0.29 | 0.53 ** |
| MD | *Rhogogaster scalaris* | Tenthredinidae | oak-associated phyllophagous | 0.94 | 0.29 | 0.52 * |
| MD | *Cryptinae sp.38* | Ichneumonidae | parasitoid | 0.87 | 0.29 | 0.50 * |
| MD | *Eupalamus lacteator* | Ichneumonidae | parasitoid | 0.87 | 0.29 | 0.50 * |
| MD | *Diapria cf. conica* | Diapriidae | parasitoid | 0.82 | 0.29 | 0.48 * |
| MD | *Lasioglossum bluethgeni* | Halictidae | pollinivorous/nectarivorous | 0.82 | 0.29 | 0.48 * |
| MD | *Athalia rosae* | Tenthredinidae | other phyllophagous | 1.00 | 0.21 | 0.46 * |
| MD | *Cryptinae sp.39* | Ichneumonidae | parasitoid | 1.00 | 0.21 | 0.46 * |
| MD | *Torymus auratus* | Torymidae | parasitoid | 1.00 | 0.21 | 0.46 * |
| SD | ***Eutomostethus luteiventris*** | Tenthredinidae | other phyllophagous | 0.76 | 0.50 | 0.62 ** |
| SD | *Orthocentrinae sp.3* | Ichneumonidae | parasitoid | 0.89 | 0.47 | 0.61 ** |
| SD | *Formica fusca* | Formicidae | polyphagous | 0.81 | 0.42 | 0.58 * |
| SD | *Campopleginae sp.12* | Ichneumonidae | parasitoid | 0.67 | 0.50 | 0.58 * |
| SD | ***Trigonoderus princeps*** | Pteromalidae | parasitoid | 0.74 | 0.42 | 0.56 * |
| SD | ***Passaloecus vandeli*** | Pemphredonidae | carnivorous | 0.87 | 0.33 | 0.54 * |
| SD | *Dolichomitus terebrans* | Ichneumonidae | parasitoid | 0.77 | 0.33 | 0.51 * |
| SD | *Arge rustica* | Argidae | oak-associated phyllophagous | 1.00 | 0.25 | 0.50 * |
| SD | *Camponotus tergestinus* | Formicidae | polyphagous | 1.00 | 0.25 | 0.50 * |
| SD | *Cryptinae sp.46* | Ichneumonidae | parasitoid | 1.00 | 0.25 | 0.50 * |
| SD | *Tromatobia lineatoria* | Ichneumonidae | parasitoid | 1.00 | 0.25 | 0.50 * |

Table S7. Indicator species for decline categories estimated at the stand scale (i.e. on 30 trees, with H: healthy plots (< 30% of trees were in decline), MD: moderately declining plots (30-60% of trees were in decline), and SD: severely declining plots (> 60% of trees were in decline)), as estimated with the multipatt function (indicspecies R-package) and the IndVal index (IndVal.g) with 1,000 permutations. H: healthy, MD: moderately declining, SD: severely declining. A corresponds to the probability that a site belongs to a particular category of decline because the species has been sampled at that site, and B is the probability of sampling the species if the site corresponds to the target decline category. Species in bold are indicators at both plot and stand scales (see Table S6).

| **Stand decline category** | **Indicator species** | **Family** | **Larval trophic guild** | **A** | **B** | **IndVal index** |
| --- | --- | --- | --- | --- | --- | --- |
| H | *Chrysis fulgida* | Chrysididae | parasitoid | 1.00 | 0.36 | 0.60 ** |
| H | ***Coleocentrus soleatus*** | Ichneumonidae | parasitoid | 1.00 | 0.29 | 0.54 ** |
| H | *Andrena chrysosceles* | Andrenidae | pollinivorous/nectarivorous | 0.82 | 0.29 | 0.48 * |
| H | *Bethylus dendrophilus* | Bethylidae | parasitoid | 0.82 | 0.29 | 0.48 * |
| H | *Ectemnius lituratus* | Crabronidae | carnivorous | 1.00 | 0.21 | 0.46 * |
| MD | *Ophion minutus* | Ichneumonidae | parasitoid | 0.78 | 0.75 | 0.76 * |
| MD | *Tryphoninae sp.6* | Ichneumonidae | parasitoid | 0.72 | 0.50 | 0.60 * |
| MD | *Microgastrinae sp.6* | Braconidae | parasitoid | 0.84 | 0.31 | 0.51 * |
| MD | *Anteon infectum* | Dryinidae | parasitoid | 1.00 | 0.25 | 0.50 * |
| MD | *Aphaenogaster subterranea* | Formicidae | carnivorous | 1.00 | 0.25 | 0.50 * |
| SD | *Brachygaster minuta* | Evanidae | parasitoid | 0.88 | 0.92 | 0.90 *** |
| SD | *Strongylogaster multifasciata* | Tenthredinidae | other phyllophagous | 0.66 | 0.83 | 0.74 ** |
| SD | *Tiphia femorata* | Tiphiidae | parasitoid | 0.93 | 0.58 | 0.74 ** |
| SD | *Sphecodes sp.* | Halictidae | pollinivorous/nectarivorous | 0.67 | 0.75 | 0.71 * |
| SD | ***Eutomostethus luteiventris*** | Tenthredinidae | other phyllophagous | 0.85 | 0.50 | 0.65 ** |
| SD | *Pristiphora fausta* | Tenthredinidae | oak-associated phyllophagous | 0.71 | 0.58 | 0.65 ** |
| SD | *Agenioideus cinctellus* | Pompilidae | parasitoid | 0.89 | 0.42 | 0.61 ** |
| SD | *Dolerus gonager* | Tenthredinidae | other phyllophagous | 0.69 | 0.50 | 0.59 * |
| SD | *Dendrocerus sp.* | Megaspilidae | parasitoid | 0.81 | 0.42 | 0.58 ** |
| SD | *Pachyprotasis simulans* | Tenthredinidae | other phyllophagous | 1.00 | 0.33 | 0.58 ** |
| SD | ***Trigonoderus princeps*** | Pteromalidae | parasitoid | 0.77 | 0.42 | 0.57 ** |
| SD | *Triaspis sp.7* | Braconidae | parasitoid | 0.96 | 0.33 | 0.57 ** |
| SD | *Aneugmenus padi* | Tenthredinidae | other phyllophagous | 0.88 | 0.33 | 0.54 * |
| SD | ***Passaloecus vandeli*** | Pemphredonidae | carnivorous | 0.85 | 0.33 | 0.53 * |
| SD | *Tenthredopsis nassata* | Tenthredinidae | other phyllophagous | 0.85 | 0.33 | 0.53 * |
| SD | *Discoelius zonalis* | Vespidae | carnivorous | 1.00 | 0.25 | 0.50 * |
| SD | *Polistes nimpha* | Vespidae | social polyphagous | 1.00 | 0.25 | 0.50 * |

Table S8. Effect of the proportion of declining oaks at plot (left) and stand (right) scales on the abundance of oak-dwelling Hymenoptera, and on the abundance of guilds (larval trophic guilds, larval nesting guild), families (abundance > 30 ind.) and taxa (abundance > 30 ind.). GLMMs were fitted for the negative binomial family (NB), the Poisson family (P) or the log-normal family (Log), with forest and stand as random effects. Models with either linear or quadratic decline variables were tested. When the quadratic model is the best model, d1 and d2 are displayed, with d1 corresponding to the linear form of the decline and d2 to the quadratic form of the decline (Y= d1*X+d2*X²). Max. is the proportion of decline corresponding to the maximum abundance observed at peak of the quadratic curve. ΔAICc = AICc (decline model) – AICc (null model). The best model between plot and stand scales is highlighted in bold. Only significant relationships are shown (*** *p* < 0.001, ** *p* < 0.01, * *p* < 0.05).

| **Guild / taxa** | **Abund.** | **Dist. fam.** | **Plot decline** | | | | | |  | **Stand decline** | | | | | |
| --- | --- | --- | --- | --- | --- | --- | --- | --- | --- | --- | --- | --- | --- | --- | --- |
|  |  |  | **Delta AICc** | **Estimate** | **S. E.** | **z or t value** | **Marginal R²** | **Max.** |  | **Delta AICc** | **Estimate** | **S. E.** | **z or t value** | **Marginal R²** | **Max.** |
| **Larval trophic guilds** |  |  |  |  |  |  |  |  |  |  |  |  |  |  |  |
| Idiobiont parasitoid | 5,641 | Log | -2.04 | -0.62 ** | 0.22 | -2.77 | 0.10 |  |  | **-3.31** | **-0.77 *** | **0.28** | **-2.79** | **0.13** |  |
| Koinobiont parasitoid | 4,113 | Log | 3.58 |  |  |  |  |  |  | **-3.36** | **d1. 2.43 *** | **0.84** | **2.91** | **0.06** | **0.46** |
|  |  |  |  |  |  |  |  |  |  |  | **d2. -2.64 *** | **0.90** | **-2.93** |  |  |
| Pollinivorous/Nectarivorous | 1,246 | Log | -3.05 | d1. 3.19 * | 1.5 | 2.13 | 0.07 | **0.58** |  | **-4.30** | **1.56 *** | **0.6** | **2.63** | **0.15** |  |
|  |  |  |  | d2. - 2.74 . | 1.6 | -1.72 |  |  |  |  |  |  |  |  |  |
| Polyphagous | 1,281 | NB | **-9.35** | **1.29 ***** | **0.1** | **113.57** | **0.19** |  |  | -3.99 | 1.22 *** | 0.1 | 112.82 | 0.14 |  |
| Xylophagous | 149 | NB | **-3.51** | **-3.47 *** | **1.5** | **-2.33** | **0.35** |  |  | 2.57 | - | - | - | - |  |
| **Larval nesting guilds** |  |  |  |  |  |  |  |  |  |  |  |  |  |  |  |
| Gall-nesters | 5,170 | Log | 0.70 | - | - | - | - |  |  | **-5.24** | **d1. 3.13 ns** | **1.98** | **1.58** | **0.25** | **0.34** |
|  |  |  |  |  |  |  |  |  |  |  | **d2. -4.63 *** | **2.17** | **-2.13** |  |  |
| Specialist soil-nesters | 1,376 | NB | 0.10 | **-** | **-** | **-** | **-** |  |  | **-4.72** | **1.66 **** | **0.55** | **3.00** | **0.18** |  |
| Specialist wood-nesters | 1,708 | Log | -2.15 | 0.64 * | 0.26 | 2.41 | 0.09 |  |  | **-2.91** | **0.83 *** | **0.32** | **2.58** | **0.13** |  |

Tab. S8 continued

| **Guild / taxa** | **Abund.** | **Dist. fam.** | **Plot decline** | | | | | |  | **Stand decline** | | | | | |
| --- | --- | --- | --- | --- | --- | --- | --- | --- | --- | --- | --- | --- | --- | --- | --- |
|  |  |  | **Delta AICc** | **Estimate** | **S. E.** | **z or t value** | **Marginal R²** | **Max.** |  | **Delta AICc** | **Estimate** | **S. E.** | **z or t value** | **Marginal R²** | **Max.** |
| **Families** |  |  |  |  |  |  |  |  |  |  |  |  |  |  |  |
| Evanidae | 33 | P | -3.13 | 2.87 ** | 1.1 | 2.74 | 0.15 |  |  | **-11.88** | **5.03 ***** | **1.3** | **3.97** | **0.35** |  |
| Formicidae | 1,339 | NB | **-8.98** | **1.23 ***** | **0.1** | **113.00** | **0.18** |  |  | -3.35 | 1.14 ** | 0.1 | 109.721 | 0.13 |  |
| Halictidae | 924 | Log | -3.94 | d1. 3.81 * | 0.6 | 2.17 | 0.06 | 0.59 |  | **-4.90** | **1.82 *** | **0.7** | **2.70** | **0.12** |  |
|  |  |  |  | d2. -3.25 . | 0.6 | -1.75 |  |  |  |  |  |  |  |  |  |
| Ichneumonidae | 4,242 | Log | 3.89 | - | - | - | - |  |  | **-4.46** | **d1. 2.24 **** | **0.7** | **3.40** | **0.11** | **0.47** |
|  |  |  |  | - | - | - | - |  |  |  | **d2. -2.37 **** | **0.7** | **-3.33** |  |  |
| Pamphiliidae | 38 | NB | -1.55 | - | - | - | - |  |  | **-4.77** | **3 *** | **1.1** | **2.66** | **0.21** |  |
| Perilampidae | 2,614 | NB | -6.30 | -1.72 ** | 0.5 | -3.24 | 0.38 |  |  | **-9.87** | **-2.67 *** | **0.8** | **-3.45** | **0.42** |  |
| Tiphiidae | 92 | P | **-2.44** | **2.39 *** | **1.1** | **2.21** | **0.05** |  |  | -2.17 | 4.24 * | 1.9 | 2.29 | 0.12 |  |
| Xiphydriidae | 149 | NB | **-3.51** | **-3.47 *** | **1.5** | **-2.33** | **0.35** |  |  | 2.57 | - | - | - | - |  |
| **Taxa** |  |  |  |  |  |  |  |  |  |  |  |  |  |  |  |
| *Agrypon flaveolatum* | 32 | P | **-2.63** | **-2.18 *** | **1.1** | **-2.00** | **0.11** |  |  | 2.32 | - | - | - | - |  |
| *Bassus sp.1* | 38 | NB | **-4.54** | **d1. 9.06 *** | **3.5** | **2.58** | **0.36** | **0.36** |  | 0.07 | - | - | - | - |  |
|  |  |  |  | **d2. -12.59 **** | **4.7** | **-2.67** |  |  |  |  |  |  |  |  |  |
| *Brachygaster minuta* | 33 | P | -3.13 | 2.87 * | 1.1 | 2.74 | 0.15 |  |  | **-11.88** | **5.03 ***** | **1.3** | **3.97** | **0.35** |  |
| *Ctenopelmatinae sp.1* | 88 | P | **-86.47** | **d1. 41.56 ***** | **8.3** | **4.99** | **0.53** | **0.40** |  | 1.88 | - | - | - | - |  |
|  |  |  |  | **d2. -51.48 ***** | **11.3** | **-4.55** |  |  |  |  |  |  |  |  |  |
| *Ctenopelmatinae sp.7* | 73 | P | **-20.70** | **2.92 ***** | **0.7** | **4.45** | **0.18** |  |  | 2.41 | - | - | - | - |  |
| *Deuteragenia vechti* | 44 | P | **-6.11** | **-3.05 *** | **1.2** | **-2.62** | **0.14** |  |  | 0.11 | - | - | - | - |  |
| *Dolichoderus quadripunctatus* | 718 | NB | **-6.30** | **1.49 *** | **0.5** | **3.06** | **0.18** |  |  | -3.30 | 1.34 * | 0.6 | 2.38 | 0.12 |  |
| *Earinus gloriatorius* | 43 | NB | **-4.19** | **d1. 7.77 ***** | **1.7** | **4.50** | **0.21** | **0.42** |  | 2.57 | - | - | - | - |  |
|  |  |  |  | **d2. -9.29 ***** | **2.0** | **-4.73** |  |  |  |  |  |  |  |  |  |
| *Empria tridens* | 30 | P | **-2.72** | **-2.18 *** | **0.9** | **-2.33** | **0.11** |  |  | 1.07 | - | - | - | - |  |
| *Macrocentrus nitidus* | 42 | NB | **-3.08** | **d1. 5.12 ***** | **1.2** | **4.15** | **0.24** | 0.33 |  | -0.01 | - | - | - | - |  |
|  |  |  |  | **d2. -7.78 ***** | **1.6** | **-4.98** |  |  |  |  |  |  |  |  |  |

Tab. S8 continued

| **Guild / taxa** | **Abund.** | **Dist. fam.** | **Plot decline** | | | | | |  | **Stand decline** | | | | | |
| --- | --- | --- | --- | --- | --- | --- | --- | --- | --- | --- | --- | --- | --- | --- | --- |
|  |  |  | **Delta AICc** | **Estimate** | **S. E.** | **z or t value** | **Marginal R²** | **Max.** |  | **Delta AICc** | **Estimate** | **S. E.** | **z or t value** | **Marginal R²** | **Max.** |
| *Myrmecina graminicola* | 39 | P | **-10.86** | **d1. 12.48 **** | **3.8** | **3.29** | **0.30** | **0.51** |  | -3.03 | d1. 9.87 ** | 3.7 | 2.64 | 0.21 | 0.51 |
|  |  |  |  | **d2. -12.13 **** | **4.0** | **-3.04** |  |  |  |  | d2. -9.68 * | 4.1 | -2.39 |  |  |
| *Perilampus ruficornis* | 2,614 | NB | -8.50 | d1. 2.00 *** | 0.1 | 131.40 | 0.38 | 0.23 |  | **-22.79** | **d1. 3.53 ***** | **0.1** | **124.97** | **0.67** | **0.24** |
|  |  |  |  | d2. -4.36 *** | 0.1 | -287.19 |  |  |  |  | **d2. -7.43 ***** | **0.1** | **-262.53** |  |  |
| *Priocnemis perturbator* | 38 | P | **-4.30** | **d1. 9.73 *** | **3.9** | **2.47** | **0.17** | **0.58** |  | 1.33 | - | - | - | - |  |
|  |  |  |  | **d2. -8.37 *** | **3.9** | **-2.16** |  |  |  |  |  |  |  |  |  |
| *Strongylogaster multifasciata* | 52 | P | **-12.98** | **d1. 15.97 **** | **4.7** | **3.40** | **0.31** | **0.67** |  | -1.43 | - | - | - | - |  |
|  |  |  |  | **d2. -11.87 **** | **3.5** | **-3.39** |  |  |  |  |  |  |  |  |  |
| *Tiphia femorata* | 92 | P | **-2.44** | **2.39 *** | **1.1** | **2.21** | **0.04** |  |  | -2.17 | 4.24 * | 1.9 | 2.29 | 0.12 |  |
| *Vespa crabro* | 240 | NB | **-3.70** | **1.61 *** | **0.7** | **2.43** | **0.10** |  |  | -0.27 | - | - | - | - |  |
| *Xiphydria longicollis* | 149 | NB | **-3.51** | **-3.47 *** | **1.5** | **-2.33** | **0.35** |  |  | 2.57 | - | - | - | - |  |
| *Xorides praecatorius* | 30 | P | 1.37 | - | - | - | - |  |  | **-3.07** | **d1. 9.88 *** | **3.9** | **2.53** | **0.22** | **0.50** |
|  |  |  |  |  |  |  |  |  |  |  | **d2. -9.82 *** | **4.0** | **-2.46** |  |  |

Table S9. Effect of the proportion of declining oaks (at plot and stand scales) on the species richness of oak-dwelling Hymenoptera, and on the species richness of guilds (larval trophic guilds, larval nesting guild) and families (abundance > 30 ind.). GLMMs were fitted for the negative binomial family (NB), the Poisson family (P) or the log-normal family (Log), with forest and stand as random effects. ΔAICc = AICc (decline model) – AICc (null model). The best model between plot and stand scales is highlighted in bold. Only significant relationships are shown (*** *p* < 0.001, ** *p* < 0.01, * *p* < 0.05).

| **Guild / taxa** | **Species richness** | **Dist. fam.** | **Plot decline** | | | | | |  | **Stand decline** | | | | |
| --- | --- | --- | --- | --- | --- | --- | --- | --- | --- | --- | --- | --- | --- | --- |
|  |  |  | **Delta AICc** | **Estimate** | **S. E.** | **z or t value** | **Marginal R²** |  |  | **Delta AICc** | **Estimate** | **S. E.** | **z or t value** | **Marginal R²** |
| **Larval trophic guilds** |  |  |  |  |  |  |  |  |  |  |  |  |  |  |
| Carnivorous | 70 | P | **-3.60** | **0.57 *** | **0.23** | **2.54** | **0.05** |  |  | -3.23 | 0.63 * | 0.25 | 2.56 | 0.05 |
| Non-oak phyllophagous | 86 | Log | 1.62 | - | - | - | - |  |  | **-3.16** | **0.89 *** | **0.64** | **2.6** | **0.11** |
| Polyphagous | 25 | P | -2.66 | 0.56 * | 0.25 | 2.79 | 0.04 |  |  | **-3.24** | **0.66 *** | **0.28** | **2.40** | **0.05** |
| **Larval nesting guilds** |  |  |  |  |  |  |  |  |  |  |  |  |  |  |
| Generalist soil-nesters | 28 | P | **-2.13** | **0.53 *** | **0.24** | **2.19** | **0.03** |  |  | -0.68 | **-** | **-** | **-** | **-** |
| Generalist stem-nesters | 40 | P | -2.92 | 0.58 * | 0.24 | 2.39 | 0.04 |  |  | **-3.65** | **0.67 *** | **0.27** | **2.53** | **0.04** |
| Generalist wood-nesters | 32 | P | **-3,89** | **0.55 *** | **0.22** | **2.57** | **0.04** |  |  | -2,72 | 0.55 * | 0.24 | 2.32 | 0.03 |
| **Families** |  |  |  |  |  |  |  |  |  |  |  |  |  |  |
| Formicidae | 29 | P | **-2.08** | **0.49 *** | **0.23** | **2.15** | **0.03** |  |  | -1.54 | - | - | - | - |
| Pamphiliidae | 9 | P | -4.51 | 2.00 ** | 0.69 | 2.89 | 0.12 |  |  | **-7.45** | **2.58 **** | **0.79** | **3.29** | **0.17** |

**SUPPLEMENTARY DATA – FIGURES**

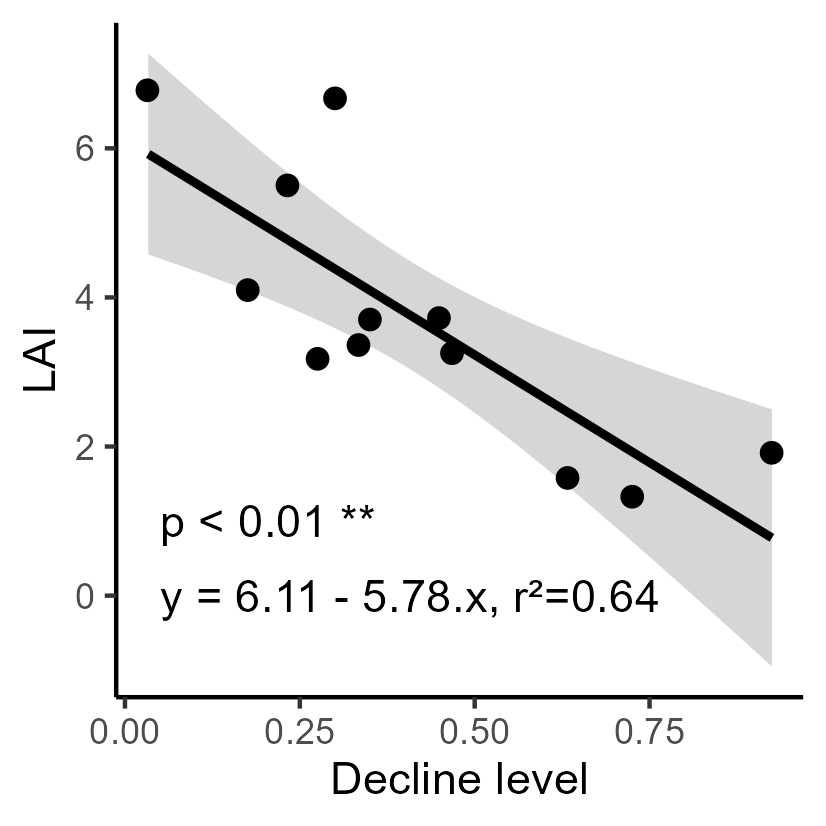

Figure S1. Relationship between mean stand Leaf Area Index (LAI) and tree decline rate at the stand scale.

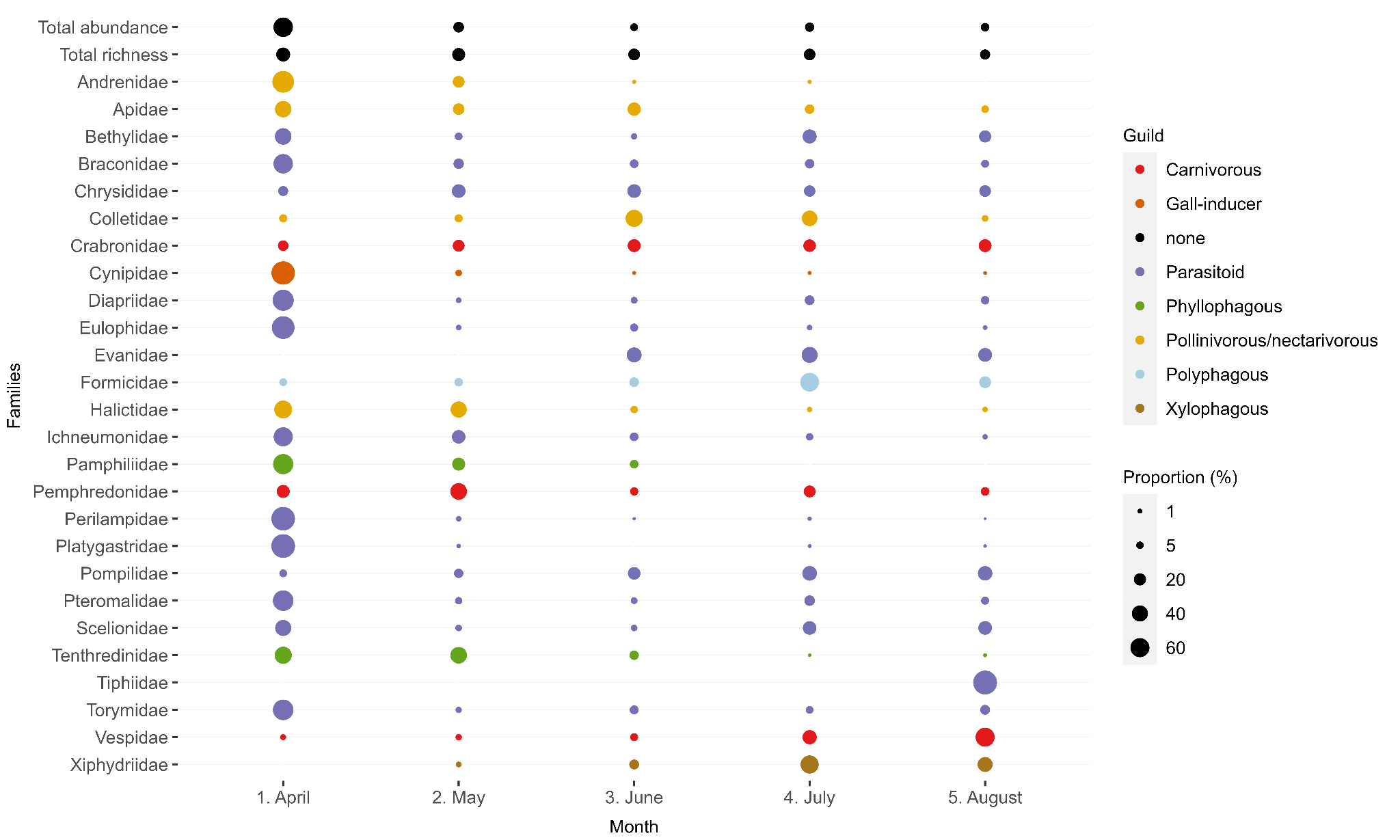

Figure S2. Seasonal activity (monthly relative proportion of individuals) of the Hymenoptera families dwelling in the canopy of the studied oak forests, arranged according to their dominant larval trophic guild. Only families with more than 30 individuals are shown.

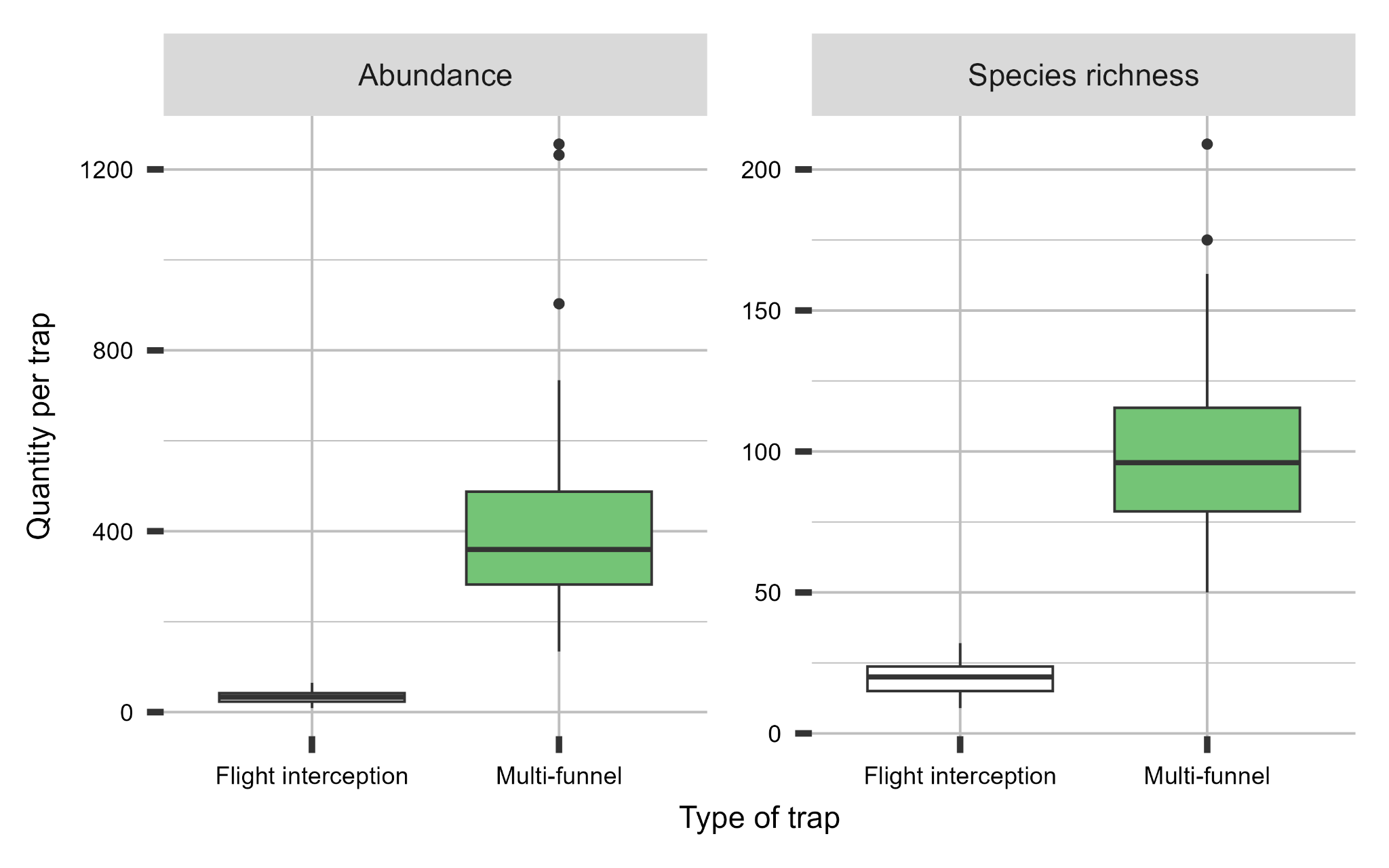

Figure S3. Mean abundance and species richness of Hymenoptera captured in transparent flight interception traps and in green multi-funnel traps.

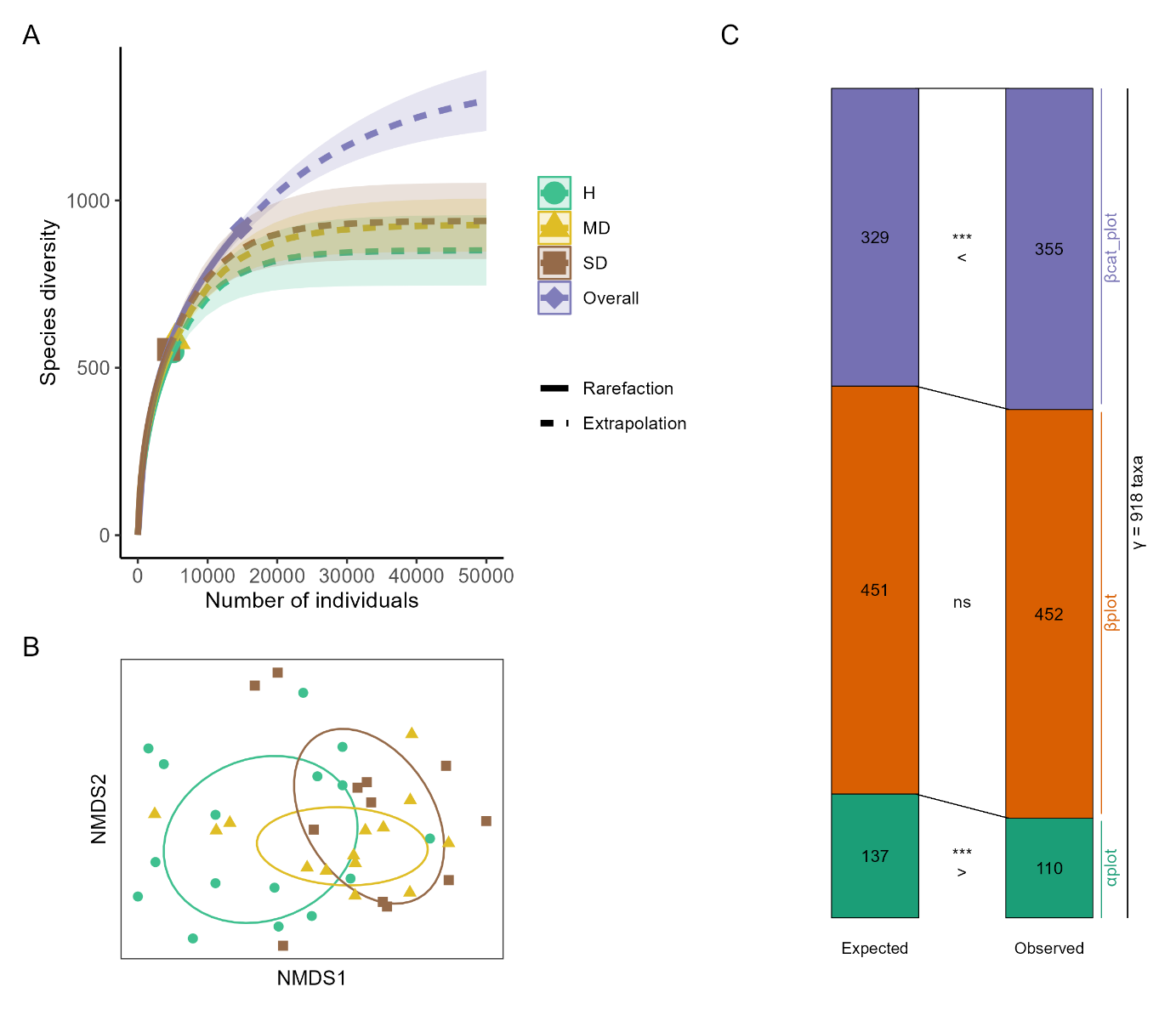
Figure S4. Responses of the community of canopy-dwelling Hymenoptera, collected from three oak forests, and 42 plots, to plot decline severity. **A)** Rarefaction curves for the overall dataset and for healthy (H: < 30% of trees are declining), moderately declining (MD: 30-60% of trees are declining) and severely declining (SD: > 60% of trees are declining) stands. **B)** NMDS ordination (k=3, stress=0.18) of species composition per plot, grouped by plot decline category. Species with less than 10 individuals were removed from the analysis. **C)** Global additive partitioning of species richness of the canopy-dwelling Hymenoptera in the three oak forests, at the plot scale. Three levels are represented: α plot (within plot), β plot (among plots), and β cat_plot (among level of plot decline). Significance levels correspond to the difference between expected and observed values, with ***: p < 0.001; . : p < 0.1.

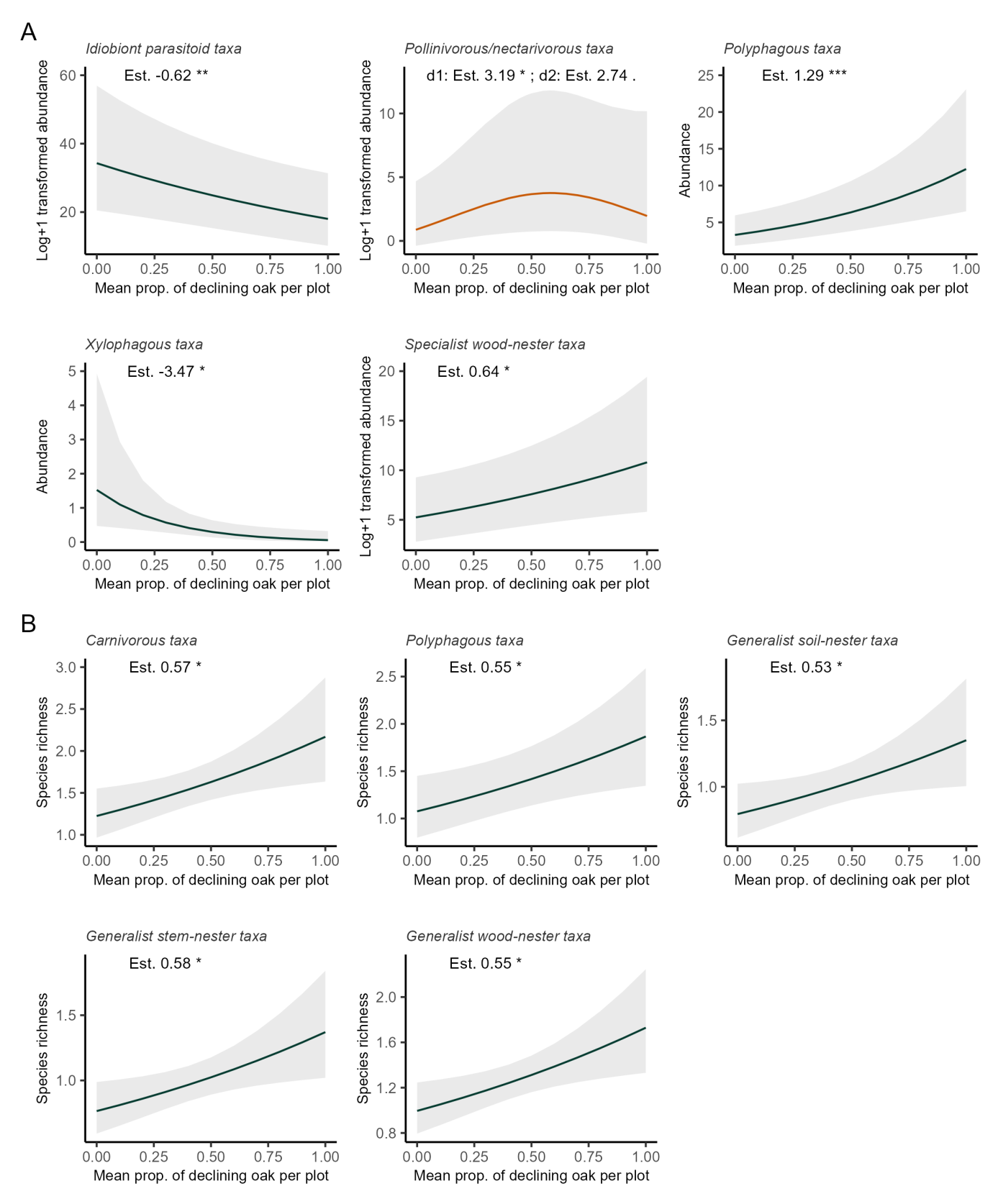
Figure S5. Graphical representations of the linear (in dark green) or quadratic (in orange) relationships between abundance **(A)** or species richness **(B)** of larval trophic and nesting guilds and the proportion of declining trees at the plot scale (10 trees). *ggeffects::ggpredict* was used to predict the values. The grey area corresponds to the confidence interval for the predicted values. When the quadratic model is the best model, d1 and d2 are displayed, with d1 corresponding to the linear form of the decline and d2 to the quadratic form of the decline. Est. corresponds to the estimate and only significant relationships are shown (*** *p* < 0.001, ** *p* < 0.01, * *p* < 0.05). Standard error, t and z value and marginal R² are available in the Tables S8 and S9.

**Figure S6**

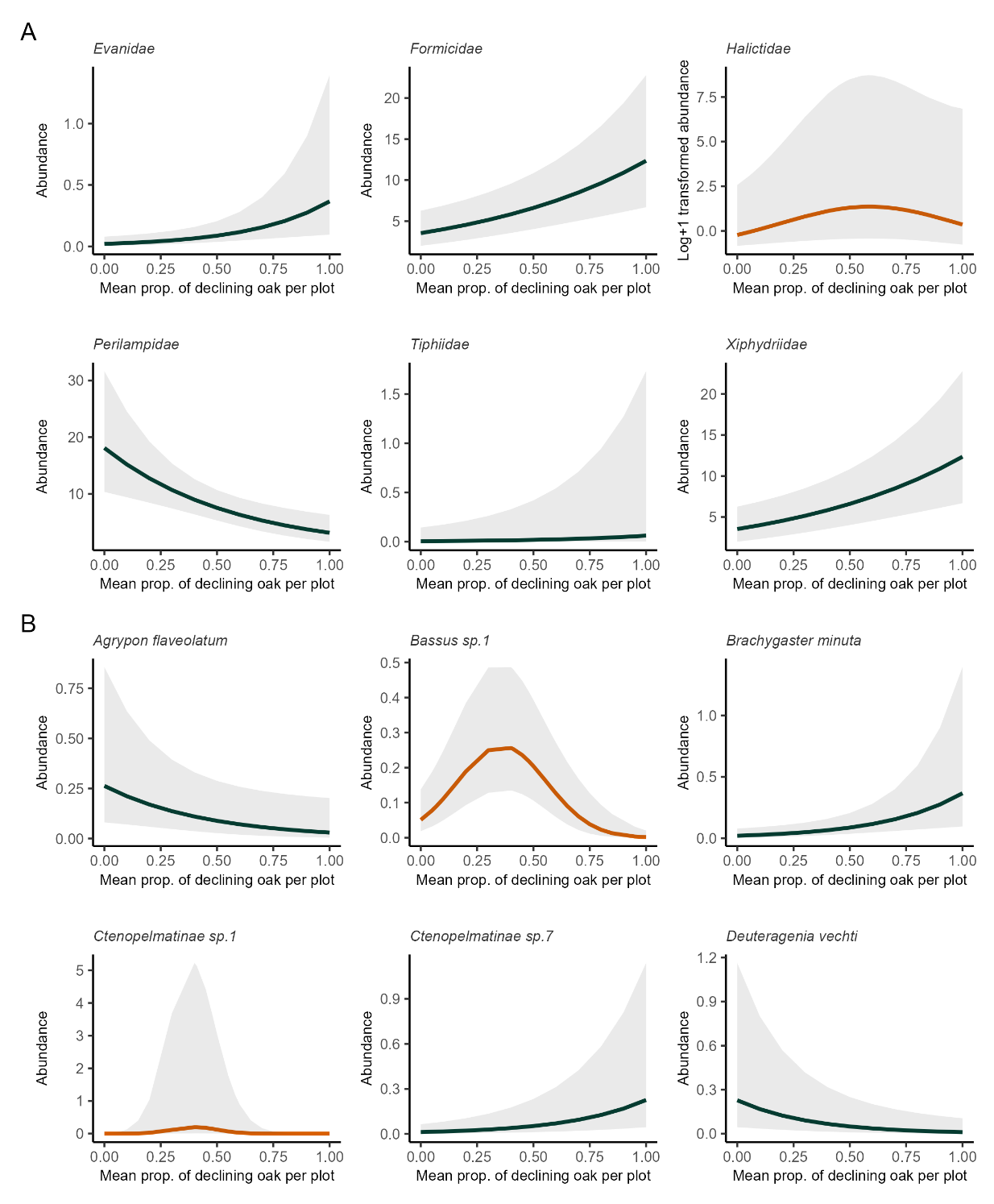

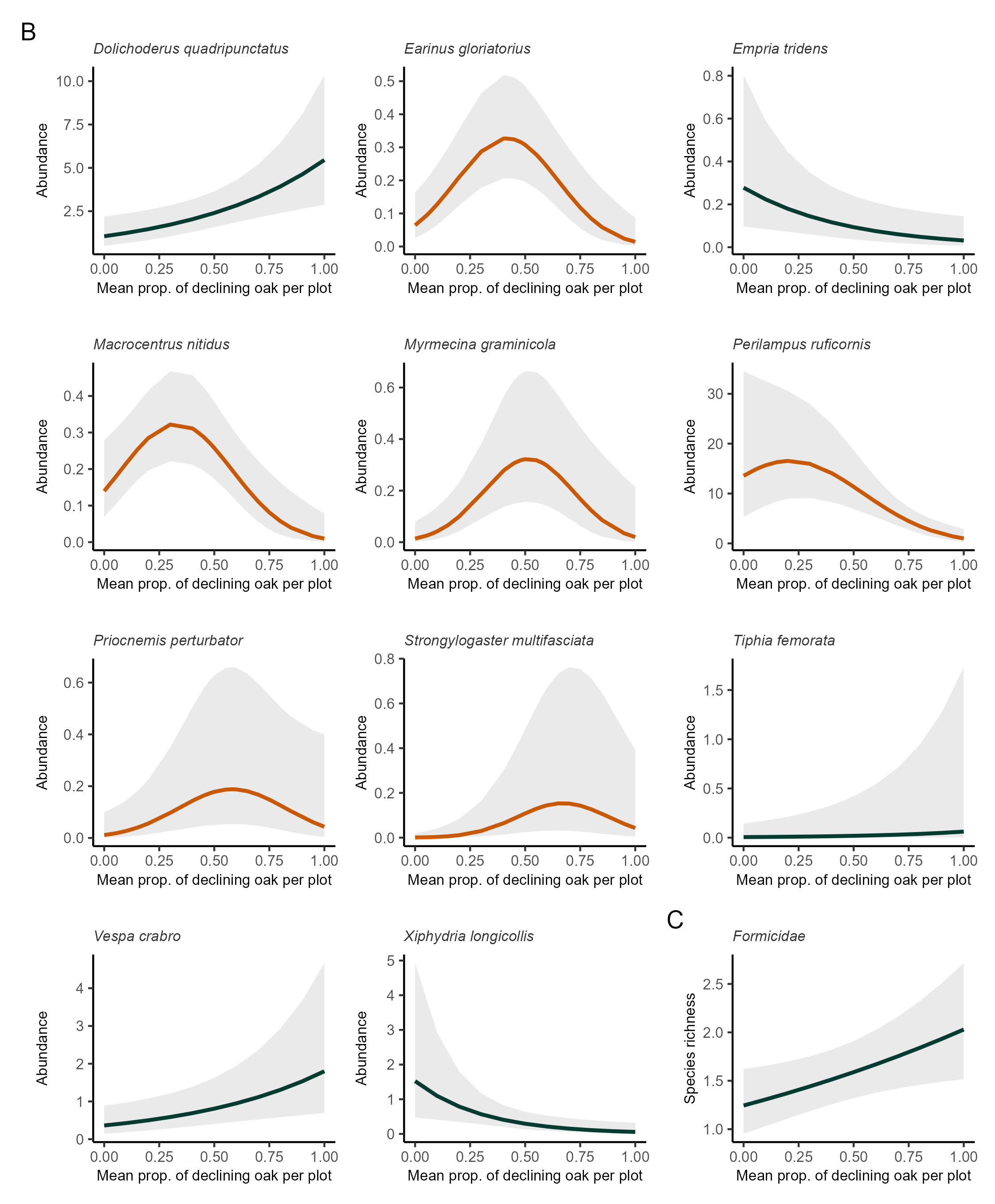

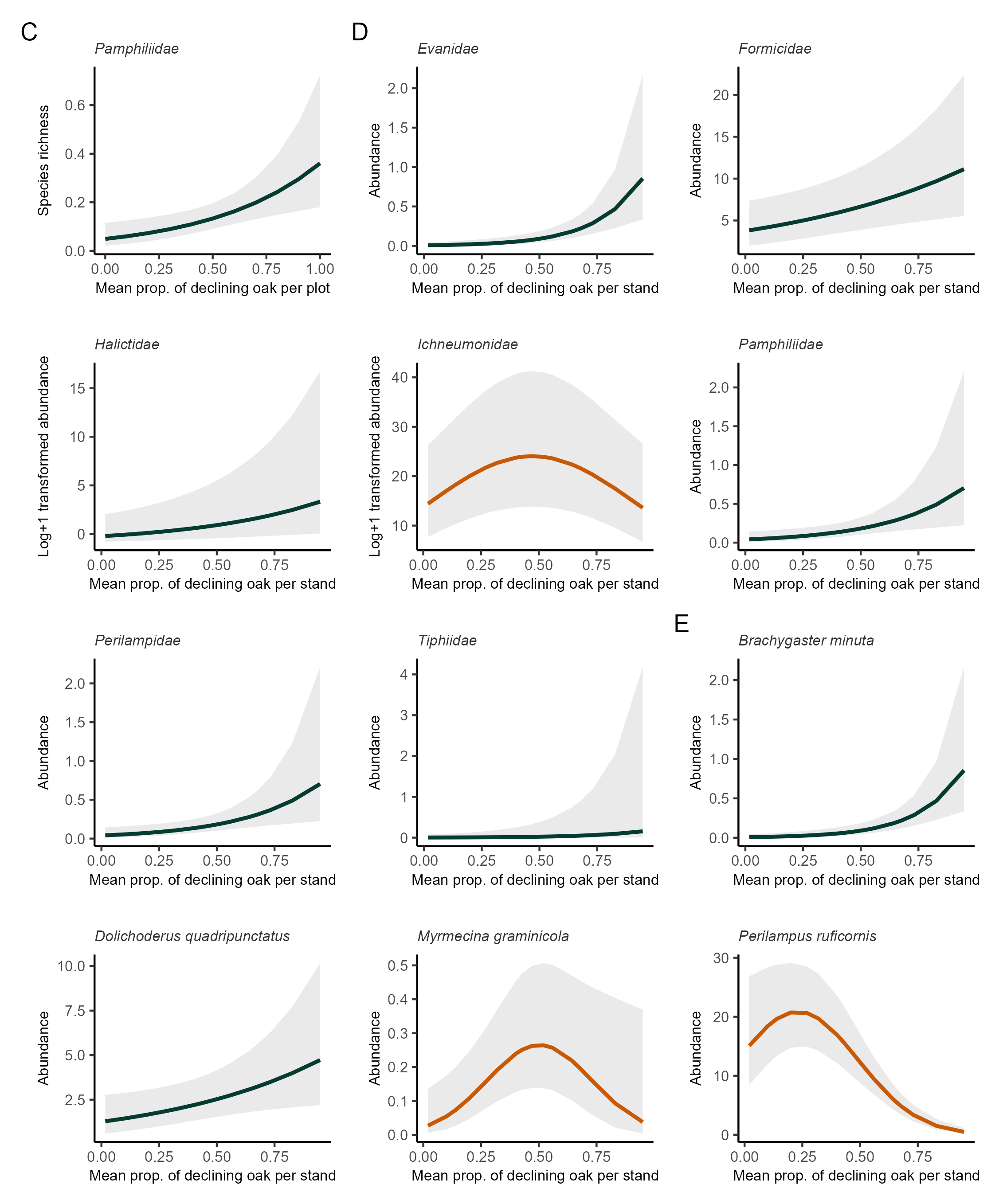

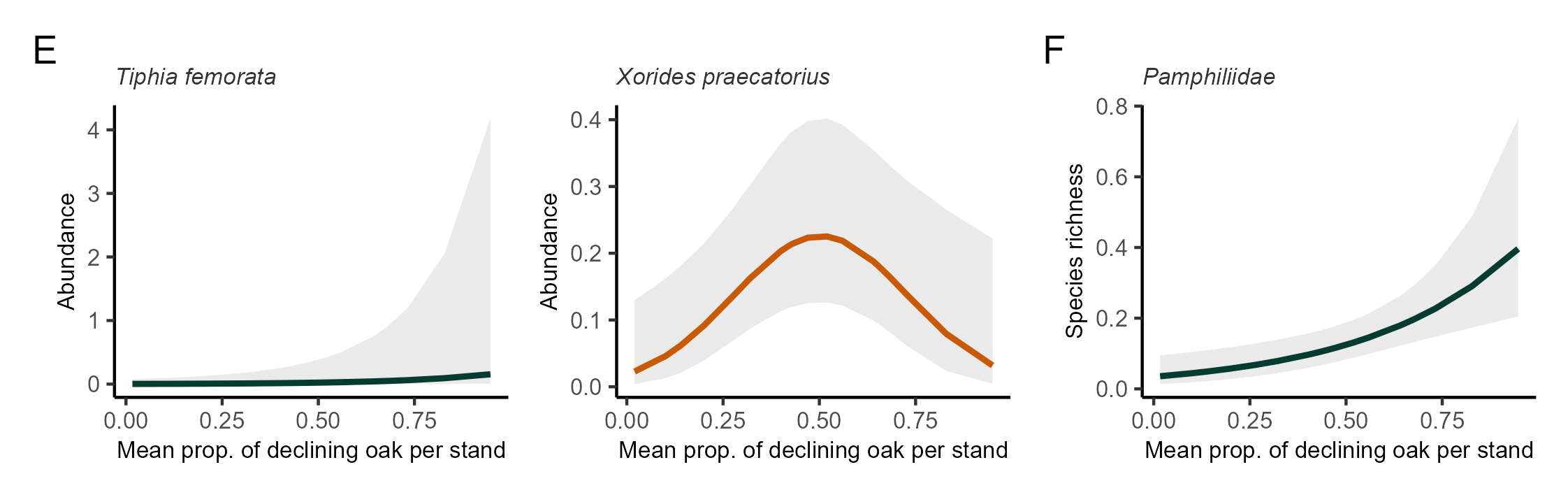

Figure S6. Graphical representations of the linear (in dark green) or quadratic (in orange) relationships between the abundance **(A, B, D, E)** of families and species or species richness **(C, F)** of families and the proportion of declining trees at plot **(A, B, C)** or stand **(D, E, F)** scales (10 and 30 trees, respectively). *ggeffects::ggpredict* was used to predict the values. The grey area corresponds to the confidence interval for the predicted values. Standard error, estimate, t and z value and marginal R² are available in the Tables S8 and S9.

**Figure S7**

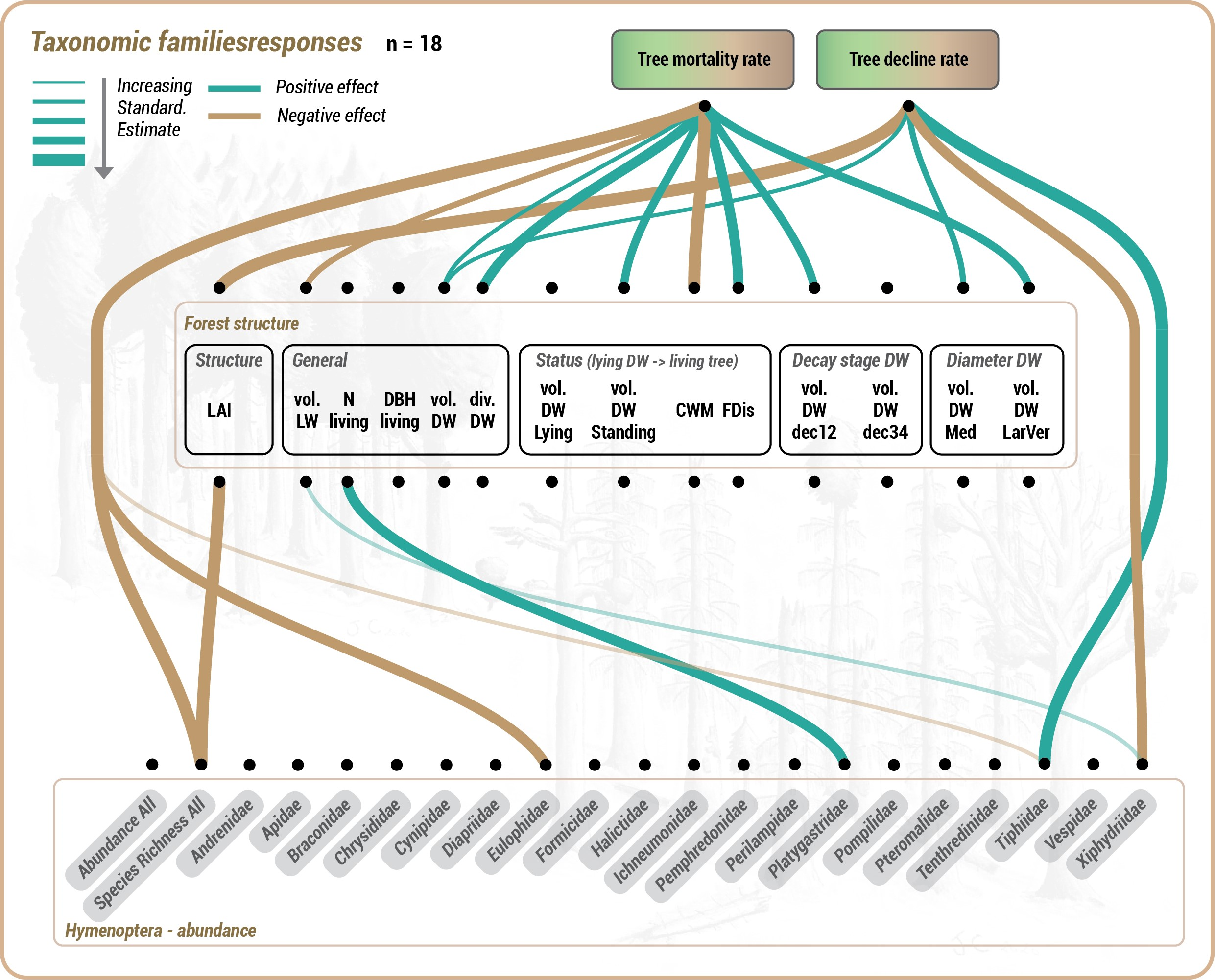

Figure S7. Results of the Structural Equation Modelling (SEM) for the effects of oak decline on the abundance of the main Hymenoptera taxonomic families through changes in forest structure. Since we tested 16 predictors, we used 0.0031 as the p.value (0.05/16). See the caption of Figure 6 for complementary information.

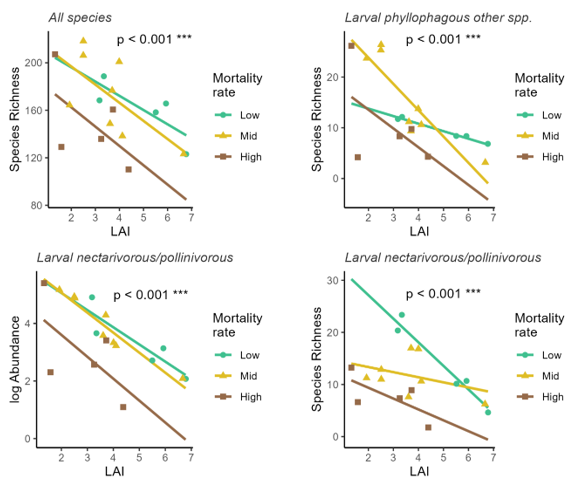

Figure S8. Relationships between overall Hymenoptera species richness and LAI (top left); log abundance of larval phytophagous species and LAI (top right); log abundance of larval species feeding on nectar and/or pollen (bottom left); and richness of larval species feeding on nectar and/or pollen (bottom right). All relationships are presented depending on the level of tree mortality rate (in colour).
